# A systematic analysis of brain tissue response to microelectrode material and size with single-cell spatial transcriptomics

**DOI:** 10.64898/2026.03.12.711361

**Authors:** Cort Thompson, Anirban Chakraborty, Lauren Wade-Kleyn, Mark Reimers, Erin Purcell

## Abstract

Implantable microelectrode arrays can interface with the central nervous system to record from and/or stimulate neural tissues to treat neurological disease and injury. The chronic tissue response to implanted electrodes is believed to be a driving factor behind microelectrode failure. Next-generation electrodes have been developed to attenuate the tissue response by reducing electrode size and/or incorporating softer materials. In this study, we used single-cell-resolution spatial transcriptomics to quantify the tissue response to implanted electrodes within custom-classified cell types in the rat brain. To test the effects of implant material and size, we assessed polyimide and silicon electrodes of 10 µm and 100 µm cross-sectional dimensions over 6-weeks post-implantation. Our data indicate that implants are associated with upregulation of inflammatory genes in glia that are coupled to damage-initiated losses in synaptic transmission and subsequent engagement of compensatory protective mechanisms (e.g., re-myelination, antioxidant production) to preserve local neurons. While bulk tissue analysis reinforced previously reported observations of glial scar consolidation over time, single cell analysis revealed an unexpected, progressive heightening of the expression of inflammatory genes in individual device-reactive astrocytes. With respect to design features, the impact of device dimensions more heavily influenced responses than material type, particularly by the 6-week time point. Our results add single-cell resolution observations to the growing use of transcriptomics to understand the biocompatibility of devices implanted in the brain.

## 1 Introduction

Implantable electrode arrays can interface with the central nervous system to record from and stimulate neural tissues to treat neurological disease and injury [1–14]. Recent innovations have aimed to improve these interfaces by enhancing device flexibility to better match the mechanical properties of brain tissue to decrease chronic inflammation and signal loss [15–17]. Nonetheless, the biological response to electrodes implanted in the brain remains poorly understood, which can undermine their use in research and clinical applications. For recording devices, the tissue response to implanted electrodes can limit the quality of the detected signal and functional longevity [15, 18–20]. For stimulating electrodes, cellular responses to the applied electric field can be remarkably complex, displaying cell type-specificity, “dose” dependence (i.e., reliance on the stimulation parameters used), and unique spatiotemporal patterns [21–24]. These intricacies, in turn, create challenges for defining safe stimulation protocols, optimizing therapies, and limiting off-target effects.

Initial characterizations of the tissue response were largely histology-based [19, 25, 26]. These observations revealed insertion-related vascular disruption, microglial activation, and astrocytic encapsulation of the electrode interface. The encapsulating glial scar, composed of microglia and astrocytes, can actively influence local neuronal populations through pro-inflammatory cytokine and neurotransmitter release while passively insulating the implanted electrode from local cells [15]. The observed glial activation during the tissue response typically coincides with a progressive loss of neuronal somata near the electrode interface within the recordable range of the device (within ∼ 100 *µ*m) [19, 27, 28]. More recent studies have identified additional factors that contribute to the tissue response through quantitative transcriptomics and in vivo imaging. Implantation of electrodes into cortical tissue damages cellular populations and the extracellular matrix, disrupts the blood-brain-barrier (BBB), and creates cellular debris, which initiates cytokine cascades and the innate immune response of the brain [29]. Both in vivo imaging and real-time polymerase chain reaction (rt-PCR) experiments confirmed that vascular damage and BBB disruption lead to perturbation in the expression of genes associated with tight junction and adherens junction proteins [30].

Recent studies have explored differentially expressed (DE) genes at the electrode interface that are involved with inflammatory cytokine cascades and the innate immune response, which contribute to the formation of glial scarring and cell death [30–36]. Oligodendrocytes, oligodendrocyte progenitor cells (OPCs), and NG2 glia are also influenced by BBB disruptions, inflammation, and the traumatic injury created by microelectrode insertion. Implantation of intracortical electrodes creates direct mechanical damage to oligodendrocytes and their corresponding myelin structure in addition to secondary damage caused through inflammatory mechanisms initiated by activated microglia and astrocytes [37]. In addition to neuronal loss, recent evidence shows structural and functional deficits in spared neurons, characterized by dendritic and spine loss, as well as reduced responsiveness to stimulation and the molecular underpinnings of spiking regularity [38]. These observations have motivated the development of “next-generation” electrodes designed with the goal of attenuating or circumventing the cortical tissue response [39, 40]. Common design strategies are to (1) reduce implant size, particularly using dimensions in the subcellular scale, and/or (2) employ softer, polymer-based materials with a Young’s modulus in closer alignment with the mechanical properties of brain tissue [41–43]. Based on these design strategies, recently published data have provided evidence that flexible devices provide reduced inflammatory responses at the histological level as well as enhanced recording stability [44]. Nonetheless, many unknowns remain with respect to optimization of design from the biological perspective [39].

In parallel with advancements on the device side, newer assays in molecular biology have revealed sophisticated classifications of neuronal and glial subtypes in the brain, as well as their unique responses to injury, disease, and inflammation. Reactive astrocytes and microglia can occupy a spectrum of heterogeneous neuroprotective to neurotoxic states, which can be understood through transcriptional profiling [45–49]. Neurotoxic astrocytes can upregulate pro-inflammatory genes such as *Serping1*, *Lcn2* and *C3*, and secrete TNF*α*, IL-6 and IL-1*β*. By contrast, IL-10 can bias resting astrocytes toward a phenotype that produces thrombospondins, IL-6, IL-10 and other neurotrophic and neuroprotective factors. Scar-forming proliferative reactive astrocytes are often found on the border of areas of degeneration and injury [45]. Microglia are likewise heterogeneous, adopting states ranging from protective to detrimental in response to disease and injury [50, 51]. Resting microglia adopt a pro-inflammatory state in response to classical stimuli such as IFN*γ* or lipopolysaccharide (LPS), wherein they exacerbate neuronal and oligodendrocyte injury through expression of TNF*α*, IL-1*β*, IL-6 and inducible nitric oxide synthase (iNOS). Conversely, IL-4, IL-13 and IL-10 promote a pro-regenerative state, exerting anti-inflammatory effects in injured brain tissue.

Recent transcriptomic analyses have identified hundreds of differentially expressed (DE) genes at the electrode-brain interface, implicating glial activation, BBB disruption, oxidative stress, iron metabolism and loss of neuronal function [33, 34, 52–54]. These studies identified upregulation of genes associated with reactive, neurotoxic glia (e.g., *Lcn2*, *C3*, *Itgam*, *C1q*, *Serping1*) and downregulation of genes associated with the BBB, suggesting compromised BBB integrity. Additionally, genes associated with neuronal function were modulated, particularly those with a role in cytoskeletal architecture (*Nefm*, *Nefh*, *Map4*), axonal transport (*Kif5a*, *Kif5b*, *Kif5c*, *Dctn1*), and synapse maintenance (*Cyfip2*, *Syn1*, *Stxbp1*, *Bsn*, *Arc*). Oligodendrocyte-associated genes (*Plp1*, *Mbp*, *Tf*, *Cnp*) were also differentially expressed at the device interface. While these data advance our understanding of the tissue response, there remains a fundamental knowledge gap regarding the individual cell types and reactive substrates that drive the results.

Here, we provide a first view of spatiotemporal transcriptional changes, registered at a single-cell resolution, as a function of electrode size and material. We used the *Xenium in situ* platform (10x Genomics) to assay a panel of 100 RNA-seq– identified genes [35,38] at single-cell resolution around implanted 10 *µ*m×10 *µ*m silicon probes (SI10), 10 *µ*m × 10 *µ*m polyimide probes (PI10), 100 *µ*m × 10 *µ*m silicon probes (SI100), and 100 *µ*m × 10 *µ*m polyimide probes (PI100). We report single-cell spatial transcriptomics from tissue implanted bilaterally with silicon or polyimide “Michigan”-style planar electrodes of subcellular or supracellular size. Fresh-frozen sections were collected at 1- and 6-weeks post-implantation to capture early- and near-chronic reactivity relative to unimplanted, naive controls. Sections were processed with the Xenium workflow and subsequently post-fixed and immunostained to enable secondary celltype identification, co-registered to the in situ data. We applied a DESeq2-based pipeline to evaluate broad transcriptional changes and implemented a complementary, custom-generated analysis pipeline leveraging cell-type-specific transcripts to quantify discrete differential expression between defined cell phenotypes at the device interface. These data provide high-resolution spatiotemporal patterns of gene expression related to neuronal function, myelination, cellular stress, glial activation, and cellular metabolism around implanted devices of varied dimensions and materials.

## 2 Methods

### 2.1 Electrode preparation

To examine device-related effects on the tissue response, polyimide and silicon devices with footprints of 10 *µ*m × 10 *µ*m and 100 *µ*m × 10 *µ*m were custom fabricated to match features of standard Michigan–style probes (A1×16-3mm, 10 *µ*m thick, 703 *µ*m**^2^** site size, 100 *µ*m spacing) (Fig. 1(a)). Silicon devices were supplied by NeuroNexus Technologies (Ann Arbor, MI). Polyimide devices were custom-fabricated and supplied courtesy of Dr. John Seymour (University of Texas Health Science Center) as previously described [38]. Devices were subsequently measured under light microscopy to verify device dimensions (Supplementary Figure 1). The assembled devices were then sterilized using ethylene oxide. Before surgical implantation, polyimide and silicon microelectrodes of all dimensions were reinforced with polyethylene glycol (PEG) 4000 to prevent buckling (Supplementary Figure 2). PEG 4000 was initially heated to 120°C and then cooled to 70°C, after which devices were dip-coated to the target thickness. To control for any potential PEG-4000-mediated effects on tissue response, all devices were coated with PEG 4000, regardless of initial dimension. To control for insertion-mediated effects on tissue response, all devices were coated to a total post-coating width of approximately 100-105 *µ*m.

**Fig. 1.**
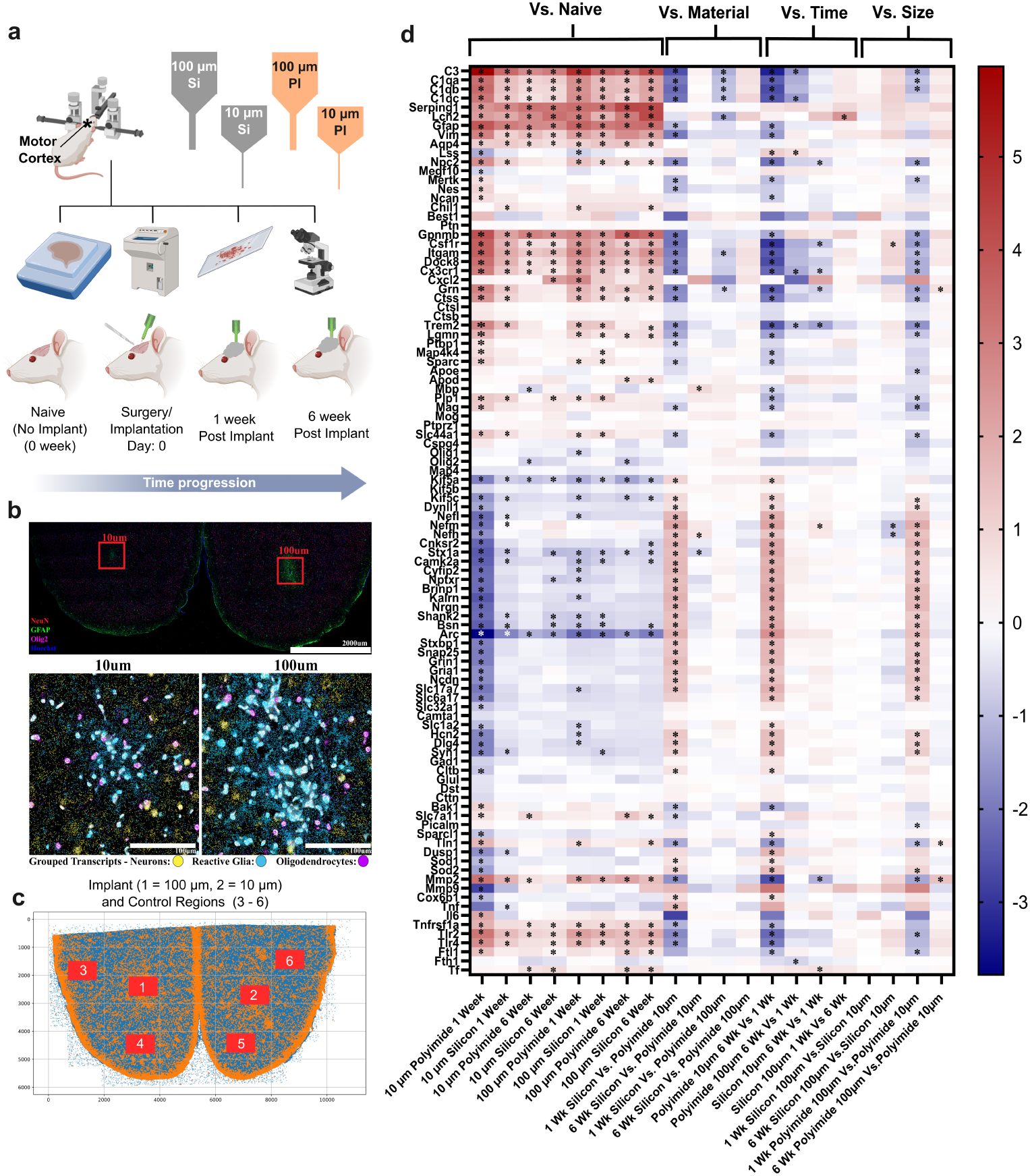
Experimental design, spatial transcriptomics workflow, and global gene expression changes at the electrode–brain interface. (a) Schematic illustrating implantation of Michigan-style microelectrode arrays into rat primary motor cortex, including electrode material (silicon or polyimide), feature size (10 µm or 100 µm), and post-implantation time points (1 week and 6 weeks). Illustrations created with BioRender.com (b) Xenium in situ spatial transcriptomics images (IHC) showing the spatial distribution of transcripts surrounding implanted electrodes. (c) Spatial mapping of transcript locations across the cortical tissue section, with regions of interest (ROIs) defined at the device interface and in distal control regions. (d) Heatmap summarizing DESeq2-generated “pseudobulk” differential gene expression in tissue regions surrounding devices across experimental contrasts, including implant versus naive tissue, material (silicon vs. polyimide), time (1 week vs. 6 weeks), and size (100 µm vs 10 µm). Colors indicate log_2_ fold change, and dots denote statistically significant differences after false discovery rate (FDR) correction.

### 2.2 Surgical implantation of electrodes

One nonfunctional device of each size (material matched, polyimide or silicon) was stereotaxically implanted bilaterally into the primary motor cortex (M1) of male Sprague–Dawley rats (aged 12–14 weeks). Animals were anesthetized with isoflurane, and a craniotomy was performed over M1 (anterior–posterior +3.0 mm, medio–lateral ±2.5 mm from Bregma). The dura was resected, and one electrode of each dimension (same material) was inserted into each hemisphere to a depth of 1.8 mm from the cortical surface to target deeper regions of the motor cortex, which are relevant for brain–machine interface applications. All devices were rapidly advanced into M1 to preserve structural integrity before the PEG shuttle softened due to dissolution. A dental-cement headcap secured the bilateral implants to two stainless-steel bone screws. Bupivacaine and Neosporin were applied topically around the incision to minimize discomfort and infection risk, and meloxicam (2 mg/kg) was administered by subcutaneous injection for postoperative analgesia. Devices remained implanted in each M1 for their designated time points (1 and 6 weeks). The Michigan State University Institutional Animal Care and Use Committee approved all surgical procedures.

### 2.3 Tissue extraction and preparation

At the terminal time point, animals were deeply anesthetized with an overdose of sodium pentobarbital and decapitated, and the brains were removed. Extracted brains were immediately cryopreserved by embedding in OCT and plunging into dry ice–cooled 2-methylbutane. Brains were cryosectioned at 10 *µ*m and mounted on Superfrost Plus slides for later histology. One transverse tissue section per animal per condition-time point (three sections total per Xenium slide) was placed horizontally on a Xenium (10x Genomics) slide within the fiducial frame. Tissue sections for Xenium analysis were chosen at an average depth of 800 *µ*m from the cortical surface to target deeper regions of the motor cortex which are relevant for brain-machine interface applications. and stored at −80°C for downstream *in situ* hybridization following delivery to the University of Michigan Advanced Genomics Research Core.

### 2.4 In situ Xenium analysis

Processing of tissue for imaging and analysis followed the manufacturer’s protocol. Fresh-frozen tissue was incubated at 37°C for 1 minute and subsequently fixed in chilled 70% methanol for 30 minutes at room temperature (RT). Fixed sections were rinsed in PBS, then permeabilized in 1% sodium dodecyl sulfate (SDS) for 2 minutes at room temperature. Slides were rinsed again and re-fixed in chilled 70% methanol for 1 hour. Sections were immersed in probe hybridization mix (10x Genomics) containing the gene panel and incubated at 50°C for 16-24 hours. The hybridization mix was removed, and sections were rinsed with PBS-T three times for 1 minute each. PBS-T was removed, and sections were incubated in ligation mix (10x Genomics) for 2 hours at 37°C. Before amplification, the ligation buffer was removed, and sections were rinsed in PBS-T three times for 1 minute each. Sections were then incubated in amplification buffer (10x Genomics) for 2 hours at 30°C. Tissue was rinsed in Tris-EDTA (Tris-Ethylenediamine tetra-acetic acid) buffer three times for 1 minute each, followed by autofluorescence quenching using 10X Genomics reducing agents and autofluorescence mix. Post-quenching, tissue was incubated in nuclear staining buffer (10x Genomics) for 1 minute and rinsed three times for 1 minute each before imaging on a Xenium Analyzer (10x Genomics).

### 2.5 Custom gene panel

For this study, we chose a panel of 100 RNA-seq-identified genes [34] which are associated with astrocytes, microglia, neurons (excitatory and inhibitory), and oligodendrocytes (Table 1). We also chose genes that are widely expressed and associated with biological processes such as innate immune response, cellular metabolism, cellular proliferation, and oxidative stress. Many of these genes were found to be differentially expressed in tissue surrounding implanted devices in previous bulk RNA-seq transcriptomic studies [34, 52]. Some genes in the panel were also chosen for exploratory purposes and/or for the identification of discrete cell types.

**Table 1.** Panel of gene probes utilized in this study. Highlighted genes were also utilized for cell-type identification in downstream analysis (Red = Excitatory Neurons, Pink = Inhibitory Neurons, Blue = Astrocytes, Orange = Oligodendrocytes, Teal = Microglia).

| Gene panel |  |  |  |  |  |  |  |  |  |
| --- | --- | --- | --- | --- | --- | --- | --- | --- | --- |
| *ApoE* | *ApoD* | *Arc* | *Aqp4* | *Bak1* | *Best1* | *Brinp1* | *Bsn* | *C3* | *Camk2a* |
| *Camta1* | *C1qa* | *C1qb* | *C1qc* | *Chil1* | *Cltb* | *Cnksr2* | *Cox6b1* | *Cspg4* | *Csf1r* |
| *Ctsb* | *Ctsl* | *Ctss* | *Cttn* | *Cxcl2* | *Cx3cr1* | *Cyfp2* | *Dlg4* | *Dock8* | *Dst* |
| *Dusp1* | *Dynll1* | *Fth1* | *Ftl1* | *Gad1* | *GFAP* | *Glul* | *GpnmB* | *Grin1* | *Gria1* |
| *Grn* | *Hcn2* | *Il6* | *Itgam* | *Kalrn* | *Kif5a* | *Kif5b* | *Kif5c* | *Lcn2* | *Lgmn* |
| *Lss* | *Mag* | *Map4* | *Map4k4* | *Mbp* | *Megf10* | *Mertk* | *Mmp2* | *Mmp9* | *Mog* |
| *Ncan* | *Ncdn* | *Nefh* | *Nefl* | *Nefm* | *Nes* | *Npc2* | *Nrgn* | *NptxR* | *Olig1* |
| *Olig2* | *Picalm* | *Plp1* | *Ptbp1* | *Ptn* | *Ptprz1* | *Serping1* | *Shank2* | *Slc1a2* | *Slc6a17* |
| *Slc7a11* | *Slc17a7* | *Slc32a1* | *Slc44a1* | *Snap25* | *Sod1* | *Sod2* | *Sparc* | *Sparcl1* | *Stx1a* |
| *Stxbp1* | *Syn1* | *Tf* | *Tln1* | *Tlr2* | *Tlr4* | *Tnf* | *Tnfrsf1a* | *Trem2* | *Vim* |

### 2.6 Immunofluorescence and imaging

Following Xenium analysis, slides were rinsed, quenched, and prepared for immunofluorescence. Tissue sections were hydrated in PBS, blocked with 10% normal donkey serum for 1 hour and incubated with primary antibodies (rabbit anti-NeuN Abcam AB177487 1:200, mouse anti-Olig2 MilliporeSigma ZMS1019 1:250, and chicken anti-GFAP Abcam AB4674 1:400) at 4°C overnight. After primary incubation, samples were rinsed three times in PBS for 10 minutes each, and secondary antibodies (Alexa Fluor 488, 594 and 647; 1:400; Thermo Fisher Scientific) were applied for 2 hours at room temperature. Slides were rinsed an additional two times for 10 minutes each, and nuclei were counterstained with 1 *µ*g/mL Hoechst. Images were acquired on a Nikon A1 inverted confocal microscope with a 20× Plan Fluor dry objective (NA 0.50) at 1024 × 1024 pixels (Supplementary Figures 3-6). For negative controls, one slide of sections per animal was processed without primary antibodies to assess autofluorescence and non-specific secondary binding.

### 2.7 Xenium analysis and statistical methods

Onboard Xenium in situ analysis and statistical methods for image processing, puncta detection, decoding, quality scoring, cell segmentation, clustering, and differential gene expression were performed as described by studies highlighted by 10x Genomics for Xenium methodologies [55, 56]. (Supplementary Figures 7-20)

To first verify that our Xenium data aligned with our previous transcriptomic studies using alternative assays, Xenium Explorer (10x Genomics) was used in conjunction with DESeq2 (R/Bioconductor) to broadly quantify differential gene expression across tissue samples using an ROI-based “pseudobulk” approach. Xenium Explorer is an image-based analysis environment that processes whole-tissue sections and reports the spatial distribution of all targets at subcellular resolution within the predefined panel. For each section, a 500 *µ*m × 500 *µ*m region of interest (ROI) centered on the implant site was defined, and per-gene transcript counts within the ROI were tabulated. Using these count values, DESeq2 estimated log**_2_** fold changes and associated wald-test *p*-values between experimental groups; *p*-values were adjusted for multiple testing using the Benjamini–Hochberg procedure. Genes with absolute log**_2_** fold change ≥ 0.6 and FDR-adjusted *p <* 0.05 were considered significantly differentially expressed (DE).

### 2.8 Identification and quantification of cell-type associated transcripts

To quantify gene expression within discrete cell-types, all single-cell-resolution analyses were performed using a custom MATLAB-based framework designed for spatial transcriptomics data generated by the 10x Genomics Xenium platform. We created a custom analysis platform, which integrates transcript localization, cell segmentation, cell-type identification, spatial region identification, expression quantification, distance-based analysis, and statistical testing of the results.

The pipeline was designed to preserve spatial information at all stages, allowing for observation of how gene expression changes within individual cells as a function of distance from implanted devices.

### 2.9 Estimation of local transcript densities

The on-board Xenium software delineates plausible estimates of cell boundaries from nuclei, assuming cell sections are convex and compact. However, most cells in the brain have complex morphologies, with many processes: approximately 90% of the volume of a neuron is in its dendrites and axons; roughly two-thirds of the volume of an astrocyte is in its processes; microglia and oligodendrocytes also have long processes that are entangled with other processes of other cells. These processes are much thinner than the 10 µm tissue section, thus many pixel locations comprise of tangled processes from dozens of different cells.

We estimate that the majority of the reads in more than half the pixel locations on the slide reflect genes expressed in neuropil rather than in cell bodies, motivating a customized approach.

Each Xenium “read” corresponds to a single detected RNA transcript and is represented by a gene name, an x-coordinate (µm), and a y-coordinate (µm). Thus, the raw dataset is a list of spatial points. No transcript is initially assigned to a specific cell; all our downstream analysis operates directly on these transcript coordinates.

To quantify gene expression, transcripts were treated as spatial point events. For any region of tissue, gene expression was first computed as simple transcript counts. For a given gene, the raw count in a region is the number of transcripts of that gene detected in the region. No assumptions about cell boundaries are made at this stage.

We observed a variation in read density of between 30% and 40% over spatial scales of ∼150 µm in a repeating grid pattern with a period of ∼650 µm, which we interpret as an artifact related to the reconstruction (stitching) of individual fields of view during Xenium imaging (Supplementary Figure 30). The abundance of most genes correlated well with the overall read density on spatial scales under 1 mm. Although some gradients of gene expression follow putative cortical layers and some reflect rostro-caudal and medio-lateral gradients of gene expression, the most plausible major source of variation on spatial scales between 50 µm and 1 mm is an uneven representation of transcripts across the surface of the slide.

To estimate how many transcripts are present locally, we computed total transcript density using kernel density estimation. In simple terms, this helps us to understand how many transcripts are nearby to this location, on average.

Using MATLAB’s ksdensity() function, a grid spacing of 40 µm was used and a smoothing bandwidth of 80 µm was chosen. This produces a smooth map, where each grid point contains the estimated number of transcripts per square micron.

In simple notation, the total density at a point is equal to the average number of all transcripts nearby. This coarse scale was chosen to smooth over the blood vessels and local variations in cell packing and the gaps in the segmentation.

#### 2.9.1 Estimation of gene-specific densities

For each gene, we computed a gene-specific density map on a finer grid by using a grid spacing of 5 µm and a bandwidth of 7 µm. This revealed how concentrated each specific gene is near a specific location. Each density value is expressed as reads per square micron, not raw counts.

#### 2.9.2 Density normalization

To remove spatial artifacts as discussed above, we normalized gene densities by the local total density. We interpolated the coarse estimate of total read density onto the finer grid and then divided the local gene density estimate by the ratio of the local total read count density to the average read count density. That means if a region has unusually high overall density, gene counts are scaled down, and if a region has unusually low overall density, gene counts are scaled up. The reference average total density was typically between 1.8 and 3.2 reads per µm**^2^**.

After this correction, the grid artifacts disappeared, and the spatial patterns aligned better with biological structures (Supplementary Figure 30).

### 2.10 Cell type inference at locations

#### Step 1: Select marker genes for density computation

From the 100 genes profiled in our Xenium data, we identified ‘marker genes’: transcripts annotated in prior sources as highly expressed in one cell type but expressed at low levels in other cell types. For each cell type, we further selected from these markers those transcripts that were highly clustered spatially; finally, we selected from those the markers that were mutually highly correlated.

These genes were organized by cell type:

- **Excitatory neurons:** “Brinp1”, “Bsn”, “Cyfip2”, “Grin1”, “Kalrn”, “Ncdn”, “Nptxr”, “Slc17a7”, “Slc6a17”, “Stx1a”, “Stxbp1”, “Syn1”
- **Inhibitory neurons:** “Gad1”, “Slc32a1”, and genes selected for neurons as above
- **Astrocytes:** “Apoe”, “Aqp4”, “Glul”, “Slc1a2”
- **Oligodendrocytes/OPCs:** “Mag”, “Mog”, “Tf”, “Apod”, “Slc44a1”
- **Microglia:** “C1qa”, “C1qb”, “C1qc”, “Csf1r”, “Ctss”, “Cx3cr1”

For microglia and oligodendrocytes, the distribution of genes was very sparse: almost all transcripts were within 15 µm of a nucleus center. For neurons, the distribution of genes was more graded: about 70% of marker gene transcripts were within 15 µm of the most likely nucleus center. This likely reflects the distribution of RNA both in the soma and in dendritic processes. For astrocytes, the marker distributions were spread out, with less than half of the four marker gene transcripts within 15 µm of a plausible nucleus. Nevertheless, the four genes selected were better correlated with each other (*r >* 0.65) than they were with *GFAP*, *Mertk*, or other annotated astrocyte-specific genes.

#### Step 2: Construct marker gene expression densities

For each region, the local expression densities of each gene were constructed on a grid with a spacing of 5 µm, using the ksdensity() function in MATLAB, with a Gaussian kernel bandwidth of 7 µm. This function computes a weighted average of counts extending over a disc of radius roughly 10 µm around each point. The unitsum density returned by ksdensity() was multiplied by the number of reads in the square to get a density expressed in reads per square micron. We normalized the raw gene density by dividing the gene density, for each gene at each point, by the ratio of the local density of all reads, at that point, to the reference density, which we set at 2.5/µm**^2^**, close to the median density for all samples.

For each of the genes considered to be a reliable cell type marker, we constructed an expression density on a finer grid of spacing 2.5 µm, using ksdensity() with a kernel bandwidth of 3.5 µm. We normalized the same way as above.

#### Step 3: First determination of cell type by marker gene densities

Many distributions of density values (maps), especially for microglia, showed a strong right skew. Biologically, most pixels don’t have transcripts of markers for a given cell type, while a few pixels, which overlap a cell body, have very high transcript counts. If we simply summed or averaged these raw values, a few extreme pixels would skew the final score and lead to inaccurate cell-type assignments. We transformed raw transcript densities into a limited score between 0 and 1 to standardize marker gene evidence across various genes and cell types. First, we set the bottom 50% of pixel gene densities to 0 (showing absence) and capped a small proportion of the highest densities at 1 (showing high-confidence presence); that proportion reflected the expected spatial area occupied by each cell type. Values falling between these cutoffs were scaled to form an exponential distribution with decay parameter K = 2 from 0 to 1, preserving their rank. This adjustment compresses extreme outliers while preserving a moderate right skew. For example, a 99th-percentile pixel may have three times more transcripts than a 97th-percentile pixel, but our scoring ensures that both only represent a high-confidence cell presence. This prevents a small number of pixels with an extremely high density of transcript pixels from influencing the composite cell-type score. Locations where multiple cell-type marker scores were high were labeled as ambiguous and were therefore excluded from analysis..

#### Step 4: Estimating fractions of somata within the slice

Using principal component analysis (PCA), we combined the customized marker-gene scores for each cell type to produce a single composite score. PCA was applied to a matrix of per-pixel marker scores, with rows representing pixels and columns representing marker genes specific to that cell type. Because the first principal component (PC1) includes a significant portion of the variance that is shared among markers co-localizing in soma-like areas, we used the normalized PC1 loadings as marker-gene weights to compute a continuous per-pixel cell-type score, resulting in a distinct spatial score map for each of the five cell types. Following this PCA-weighted construction, specific adjustments were made to improve data interpretability. Scores were adjusted to better separate excitatory and inhibitory neurons, and microglia scores were transformed with a square root to account for their sparse signals.

To assign cell types to the segmented nuclei, each nucleus centroid was projected into the fine grid, and we calculated the mean cell type score in a local neighborhood of 3×3 pixels around that centroid. This produced a 5-element score vector per nucleus. Each nucleus was assigned to the cell type with the highest score (argmax decision rule).

Finally, to quantify assignment reliability, we developed a confidence metric defined as the difference between the highest and second-highest scores for a given nucleus. Assignments with a small numerical difference between these top scores were classified as less confident or ambiguous in downstream analysis.

For analyses of the spatial distribution of gene expression in relation to the implants, the distance of every transcript and grid point from the implant surface was computed, taking into account the implant geometry. Expression and density values were then grouped into distance bins, enabling estimation of spatial gradients of gene expression and statistical testing.

### 2.11 Statistical analysis of cell-type-specific gene expression estimates

All further statistical analyses were performed on our estimates of normalized gene density values and cell-type-specific relative expression to assess whether gene expression varied across experimental conditions; we tested for the effects of implant size, implant material, time point, and implant presence. All comparisons were conducted at the animal level; individual cells were not treated as independent biological replicates.

We tested genes only within 1 or 2 cell types, in which they were highly expressed (at least 2 times more) relative to the other cell types; if a gene was expressed highly in 3 or more cell types, it was excluded from cell-type-specific analysis. For each tested gene and cell type, the estimated expression values were averaged for all cells within the specified spatial regions in each animal; these averages were the numbers used in the ANOVA procedure. Differences of these cell-specific averages between experimental groups were evaluated using analysis of variance (ANOVA) with factors representing implant presence/absence, device material, and implant size. Separate analyses were performed for the 1-week and 6-week time points.

To compensate for multiple hypothesis testing across many gene–cell-type pairs, thresholds for p-values from each ANOVA contrast were set using the Benjamini–Hochberg false discovery rate (FDR) control procedure [57]; genes that met the threshold were considered to have statistically significant condition-dependent expression.

## 3 Results

### 3.1 Description of the Xenium data

The Xenium processed data files usually reported between 80 million and 120 million reads per slide. The brain slice occupied a typical area of ∼ 40 mm**^2^** with average densities of 2–3 transcripts per *µ*m**^2^**. Four genes were disregarded in our custom analysis pipeline (*Best1*, *Il6*, *Cxcl2*, and *Tnf*) due to the consistently poor quality of these probes across all samples. All reads were included in this analysis, where the vast majority (90%) exceeded a quality value score of 20.

### 3.2 Implants alter gene expression within single cells in the brain

In this study, we have identified the spatial expression of 100 genes around implanted polyimide and silicon devices with global (tissue level, cell-type agnostic) and single cell resolution (associated with discrete nuclei) (Fig. 2). At the histological level, we qualitatively observed an expected increase in GFAP immunofluorescence and reduction in NeuN immunofluorescence around implanted electrodes. We also observed greater GFAP immunofluorescence around 100 µm relative to 10 µm implants. Then, we conducted pseudobulk DESeq2 analysis on 500 *µ*m × 500 *µ*m tissue regions surrounding implants to validate that our Xenium data aligned with differential expression from our prior transcriptomics studies [34, 52].

**Fig. 2.**
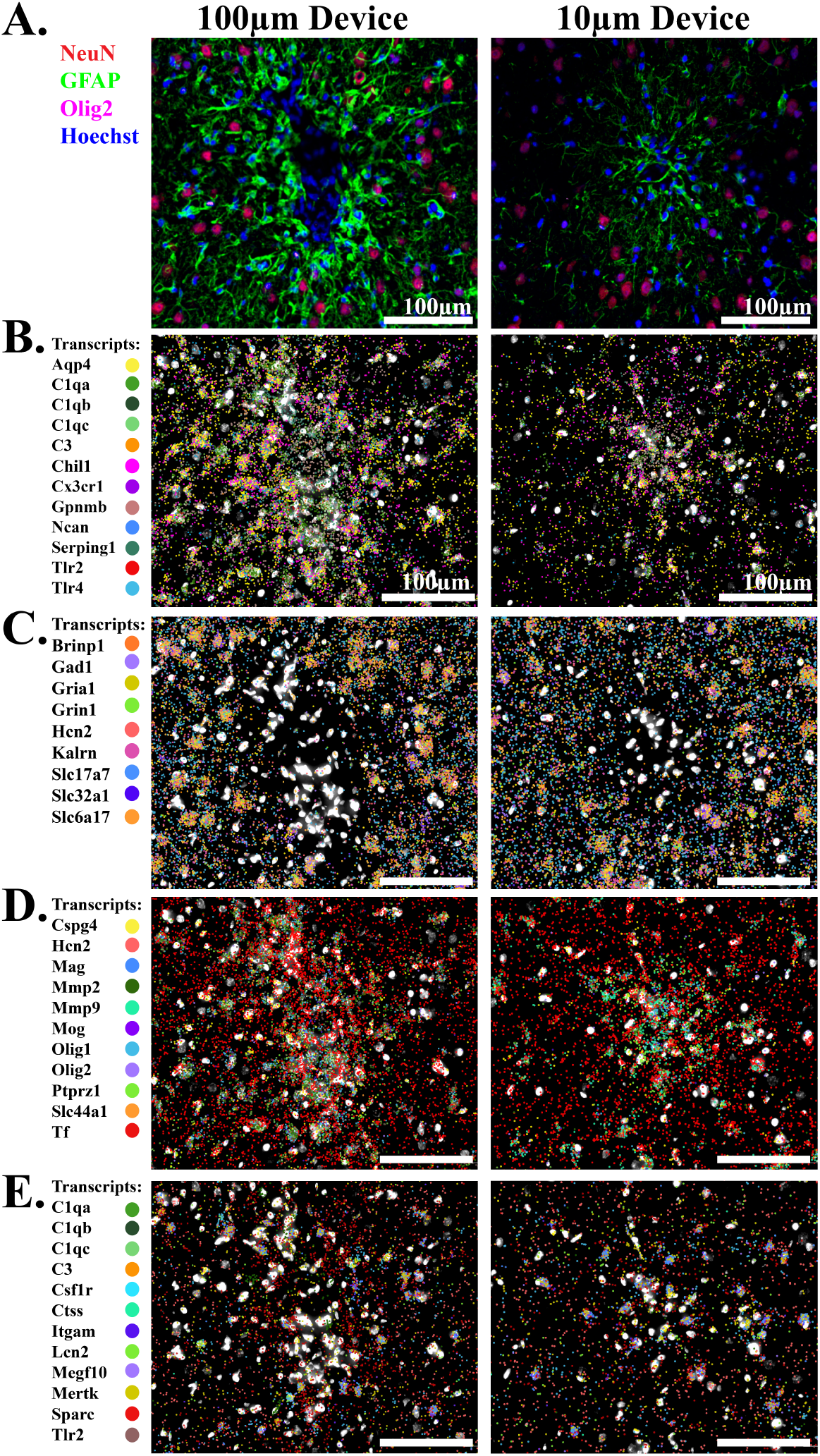
In situ spatial transcriptomics enables visualization and quantification of transcripts within individual cells around implanted electrodes. Representative confocal immunofluorescence images registered to spatial transcriptomics data from an animal with bilateral polyimide implants (10 µm and 100 µm) 6-weeks post-implantation that contain transcripts associated with reactive glia within the glial scar (B), neurons (C), oligodendrocytes (D), and modulatory glia which are not present directly within the glial scar (E). Scale bar = 100 µm.

Our initial DESeq2 results identified elevated expression of genes associated with inflammation, the complement system, glial activation, and cellular remodeling around both large and small polyimide and silicon devices, in alignment with previous reports (Fig. 1d, [34, 52]). Across conditions, implanted tissue, regardless of material or size, exhibited significant upregulation of tissue response programs. At both time points, polyimide and silicon MEAs of 10 and 100 *µ*m generated increases in genes with known associations to inflammatory responses in reactive glia (*C3*, *C1qa*, *C1qb*, *C1qc*, *Serping1*, *Lcn2*, *GFAP*, *Vim*, *Aqp4*, *Npc2*, *Gpnmb*, *Csf1r*, *Itgam*, *Dock8*, *Cx3cr1*, *Cxcl2*, *Grn*, *Ctss*, *Trem2*, *Lgmn*, *Tnfrsf1a*, *Tlr2*, *Tlr4*) and significant decreases in neuronal genes (*Kif5a*, *Kif5c*, *Dynll1*, *Stx1a*, *Bsn*, *Arc*, *Syn1*) in comparison to unimplanted tissue (Fig. 1(d)). These results align with our previous observations. However, this type of analysis falls short of identifying the individual cell types that contribute to the expression of these genes, as well as the classification of specific cellular reactive sub-states and their spatial distribution surrounding devices. Additional onboard Xenium based clustering results can be found in Supplementary Figures 21-29.

To achieve greater resolution of the transcriptional tissue response we developed in-house analysis algorithms to associate individual transcripts with individual cell types, and we identified differentially expressed genes associated with astrocytes, microglia, neurons (excitatory and inhibitory), and oligodendrocytes at the device interface. Compared to unimplanted tissue, all implants, regardless of material or dimension, elicited significantly increased gene expression associated with glial activation, glial proliferation, inflammation, remyelination, oxidative stress, and loss of neuronal function (Fig. 3). Effects were evident at 1-week post-implantation and persisted at the 6-week timepoint: reactive glia displayed pronounced per-cell upregulation of many inflammatory-related genes at both timepoints. Relatively large increases in *GFAP*, *Serping1*, and *Gpnmb* expression in astrocytes and *C3* and *Gpnmb* in microglia were conserved across timepoints. Expression of oligodendrocyte-associated genes was more nuanced and time-dependent, with a mixture of minor increases and decreases in gene expression noted at each timepoint. Notably, *Slc44a1* was upregulated and *Olig1* was downregulated at both time points. Neuronal gene expression was universally downregulated at the 1-week time point, and typically across both excitatory and inhibitory neurons (*Kif5a*, *Kif5c*, *Dlg4*, and *Cyfip2*) (Fig. 4). *Stx1a* and *Kalrn* were downregulated only in excitatory neurons. However, by 6 weeks neuronal genes were nearly universally upregulated (increases in *Gria1*, *Brinp1*, *Sod2*, *Cox6b1*, *Nefl* and *Nefh* in excitatory neurons; *Stx1a* was decreased in excitatory neurons, and *Kif5a* was decreased in both interneurons and excitatory neurons). Unlike reactive glial genes, patterns of neuronal gene expression were notably distinct across time points, both in terms of the genes expressed and the sign of expression (up or down).

**Fig. 3.**
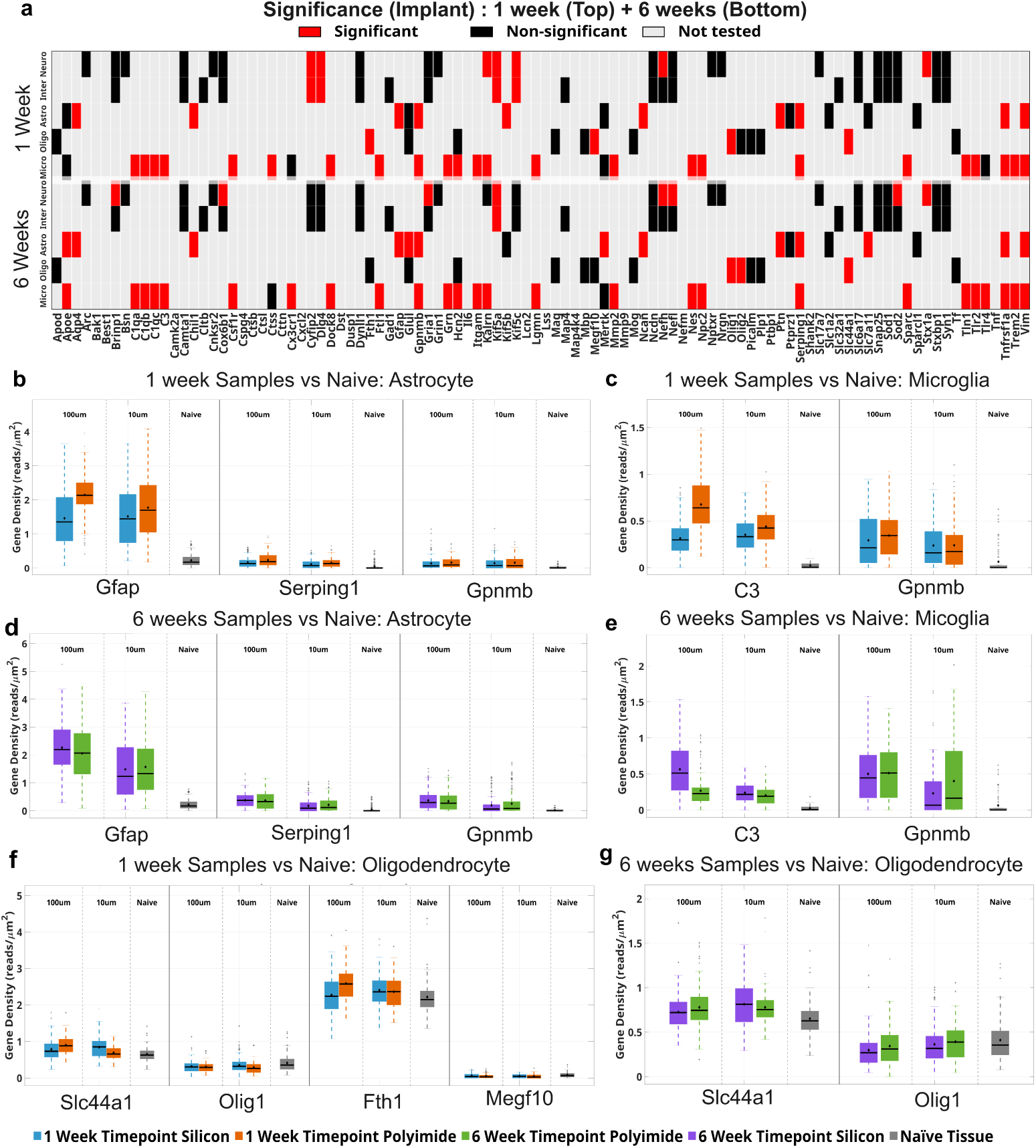
Implant-driven cell-type-specific expression changes in glia. Implant-associated differential gene expression across major glial cell types. (a) Heatmap showing statistically significant implant-driven changes in gene expression at 1 week (top) and 6 weeks (bottom) post-implantation across all cell types relative to naive tissue. Statistical significance was determined using ANOVA with Benjamini–Hochberg FDR correction (adjusted *q <* 0.05). Colors indicate status: red = significant, black = non-significant, grey = not tested due to lack of cell-type expression. (b–c) Astrocyte and microglial gene expression changes at 1 week post-implantation, highlighting increased expression of inflammatory and reactive markers. (d–e) Astrocyte and microglial gene expression changes at 6 weeks post-implantation, showing reactivity profiles. (f–g) Oligodendrocyte gene expression changes at 1 week and 6 weeks post-implantation. Boxplots represent normalized gene density (reads/ µm^2^), measured within a 500 *µ*m *×* 500 *µ*m region of interest centered on the implant, with points indicating individual samples. Sample sizes: 1-week polyimide (n = 2), 1-week silicon (n = 3), 6-week polyimide (n = 3), 6-week silicon (n = 3), naive controls (n = 3).

**Fig. 4.**
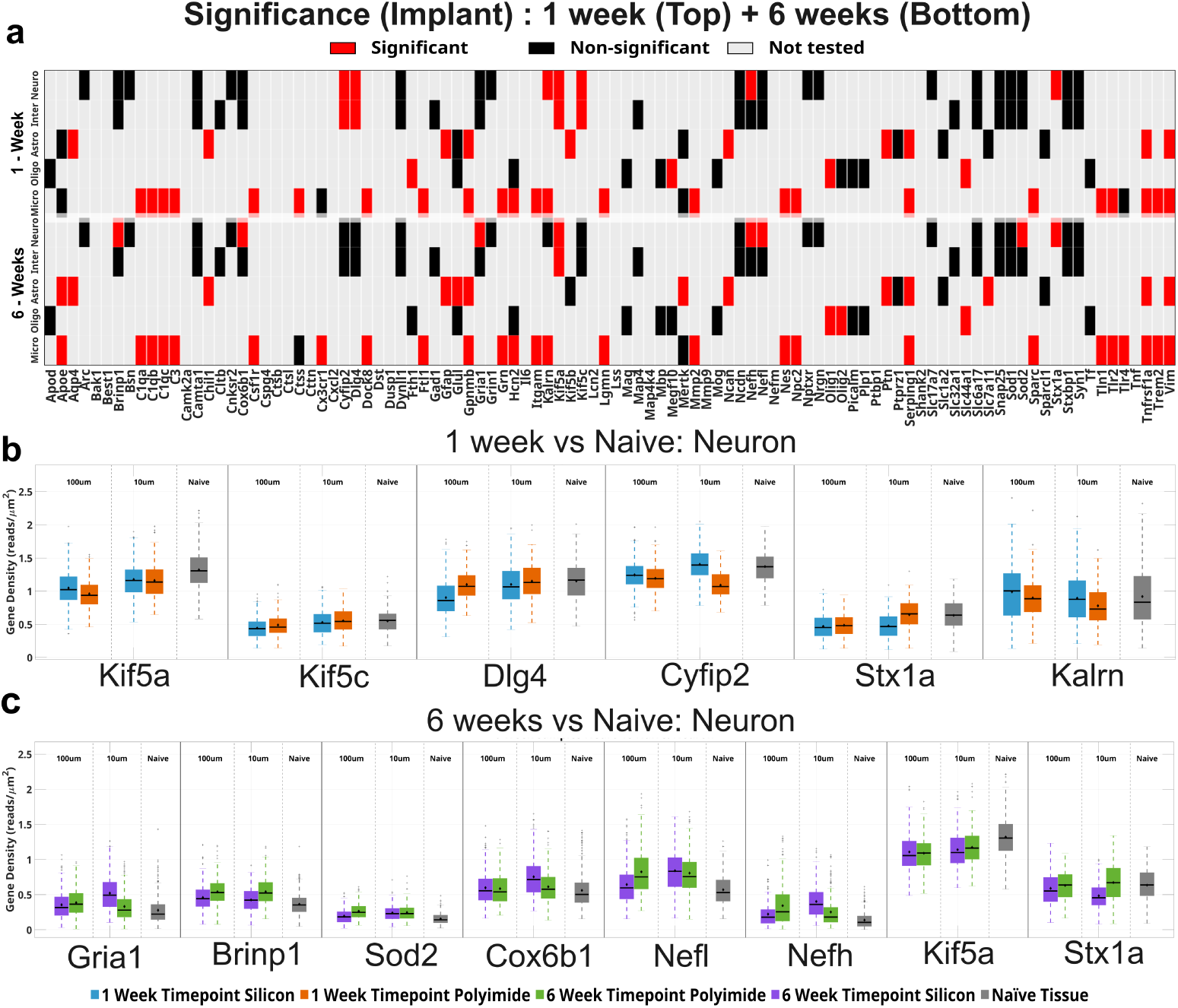
Implant-driven cell-type–specific gene expression changes in neurons. Implant-associated differential gene expression across neurons. (a) Heatmap showing statistically significant implant-driven changes in gene expression at 1 week (top) and 6 weeks (bottom) post-implantation across all cell types relative to naive tissue. Statistical significance was determined using ANOVA with Benjamini–Hochberg FDR correction (adjusted *q <* 0.05). Colors indicate status: red = significant, black = non-significant, grey = not tested due to lack of cell-type expression. (b–c) Neuronal gene expression changes at 1 week and 6 weeks post-implantation. Boxplots represent normalized gene density (reads/ µm^2^), measured within a 500 *µ*m *×* 500 *µ*m region of interest centered on the implant, with points indicating individual samples Sample sizes: 1-week polyimide (n = 2), 1-week silicon (n = 3), 6-week polyimide (n = 3), 6-week silicon (n = 3), naive controls (n = 3).

### 3.3 Implant dimensions alter gene expression in single cells in the brain

Following an initial characterization of all implant-associated effects, we contrasted effects between the microelectrode array (MEA) implant conditions—10 *µ*m polyimide, 10 *µ*m silicon, 100 *µ*m polyimide, and 100 *µ*m silicon—to reveal effects of electrode design features on gene expression at 1- and 6-weeks within specific cell types post-implantation against unimplanted naive tissue (Fig. 5.).

**Fig. 5.**
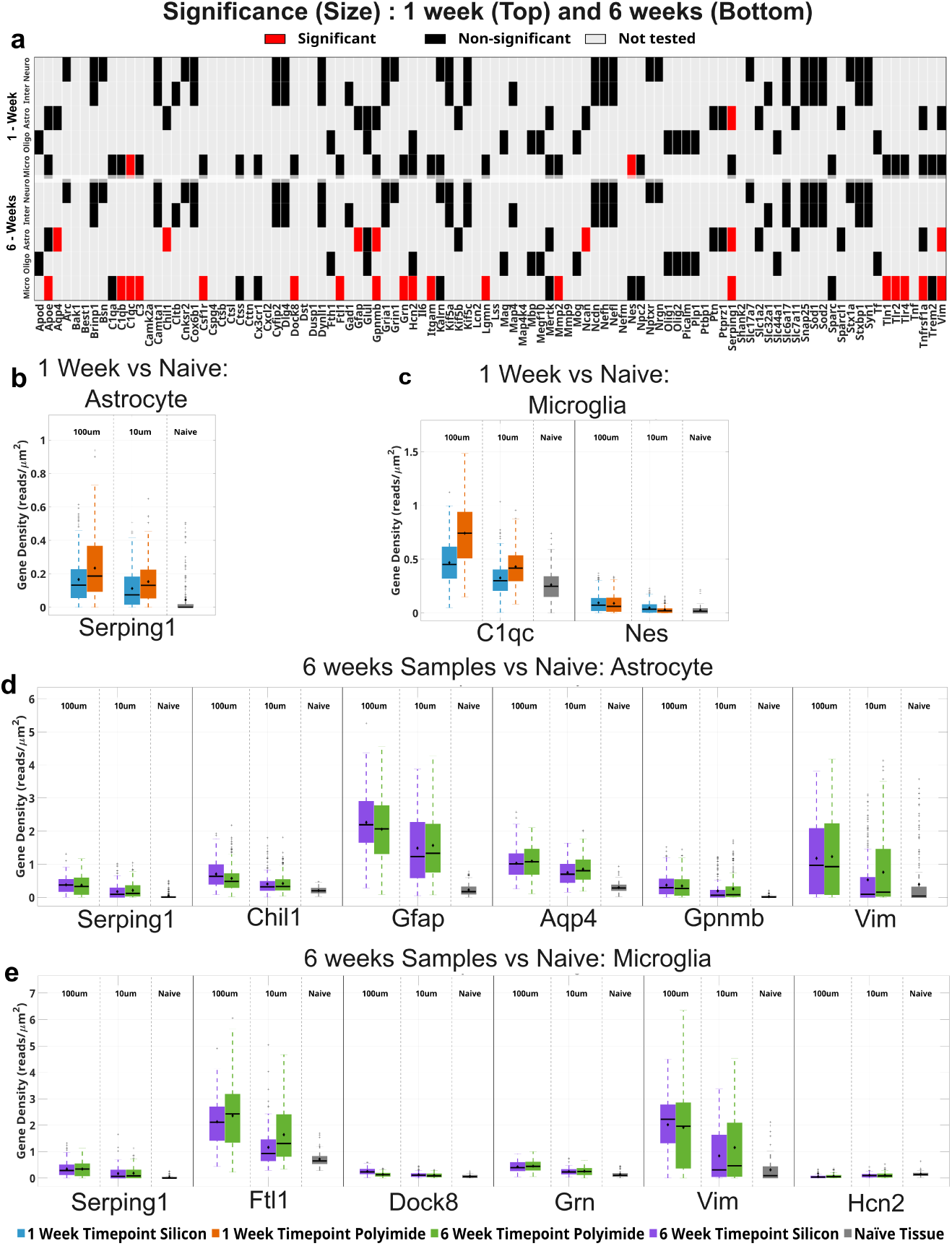
Size-dependent gene expression changes at the electrode–brain interface. (a) Heatmap showing statistically significant size-dependent differences in gene expression between 10 *µ*m and 100 *µ*m at 1 week (top) and 6 weeks (bottom) post-implantation across all cell types relative to naive tissue. Statistical significance was determined using ANOVA with Benjamini–Hochberg FDR correction (adjusted *q <* 0.05). Colors indicate status: red = significant, black = non-significant, grey = not tested due to lack of cell-type expression. (b–c) Astrocyte and microglial gene expression differences at 1 week post-implantation. (d–e) Astrocyte and microglial gene expression differences at 6 weeks post-implantation. Boxplots represent normalized gene density (reads/ µm^2^), measured within a 500 *µ*m *×* 500 *µ*m region of interest centered on the implant, with points indicating individual samples. Sample sizes: 1-week polyimide (n = 2), 1-week silicon (n = 3), 6-week polyimide (n = 3), 6-week silicon (n = 3). Each sample had both the 100 µm and 10 µm devices implanted.

We first contrasted gene expression between microelectrode arrays (MEAs) of different widths within each time point to evaluate size-dependent effects. Starting with the global assessment of all transcripts near the device using DESeq2, we found no significant difference in gene expression between 100 and 10 *µ*m silicon MEAs at 1-week post-implantation. However, at 6 weeks, *Csf1r* expression was significantly higher and *Nefm* and *Nefh* were significantly lower around 100 *µ*m silicon devices. We next compared 10 and 100 *µ*m polyimide devices within each time point. Relative to 10 *µ*m polyimide MEAs, 100 *µ*m polyimide implants exhibited significantly reduced expression of several tissue–response genes (*C3*, *C1qa*, *C1qb*, *Npc2*, *Mertk*, *Gpnmb*, *Csf1r*, *Itgam*, *Dock8*, *Cx3cr1*, *Grn*, *Ctss*, *Trem2*, *Lgmn*, *Apoe*, *Slc44a1*) and a significant increase in gene expression associated with neuronal function (*Camk2a*, *Nptxr*, *Brinp1*, *Kalrn*, *Nrgn*, *Shank2*, *Bsn*, *Arc*, *Stxbp1*, *Snap25*, *Grin1*, *Gria1*, *Ncdn*, *Slc17a7*, *Slc6a17*, *Hcn2*, *Dlg4*, *Syn1*) and neuronal structure (*Kif5c*, *Dynll1*, *Nefm*, *Nefh*, *Cnksr2*, *Stx1a*, *Cyfip2*) within identified neurons. Additionally, *Picalm*, *Dusp1* and *Mmp2* were downregulated around 100 *µ*m polyimide devices relative to 10 *µ*m polyimide devices at 1 week post-implantation. It’s possible that the smaller polyimide devices may have incurred greater insertional damage, thus explaining the heightened expression of inflammatory genes 1-week post-implantation. At the 6-week time point, the relatively few genes that were differentially expressed were upregulated surrounding the larger polyimide implants (*Mmp2*, *Tln1*, *Grn*).

We next utilized our custom scripts to assess effects within individual cell types. When evaluating effects at single-cell resolution, only 3 significant effects were noted between implants of large and small size: a minor increase in the expression of glial reactivity genes was noted around larger devices (*C1qc* and *Nes* in microglia and *Serping1* in astrocytes). At the 6-week time point, many more genes were significantly differentially expressed, and all of them were associated with the expression of reactivity and inflammatory genes in microglia and astrocytes (Fig. 6). The reactivity genes were upregulated surrounding larger devices, with the exception of *Hcn2*, which was downregulated in microglia. Inhibition of Hcn2 channels suppresses inflammation in microglia [58]. No effect of device dimension on gene expression at the single cell level was observed in neurons or oligodendrocytes.

**Fig. 6.**
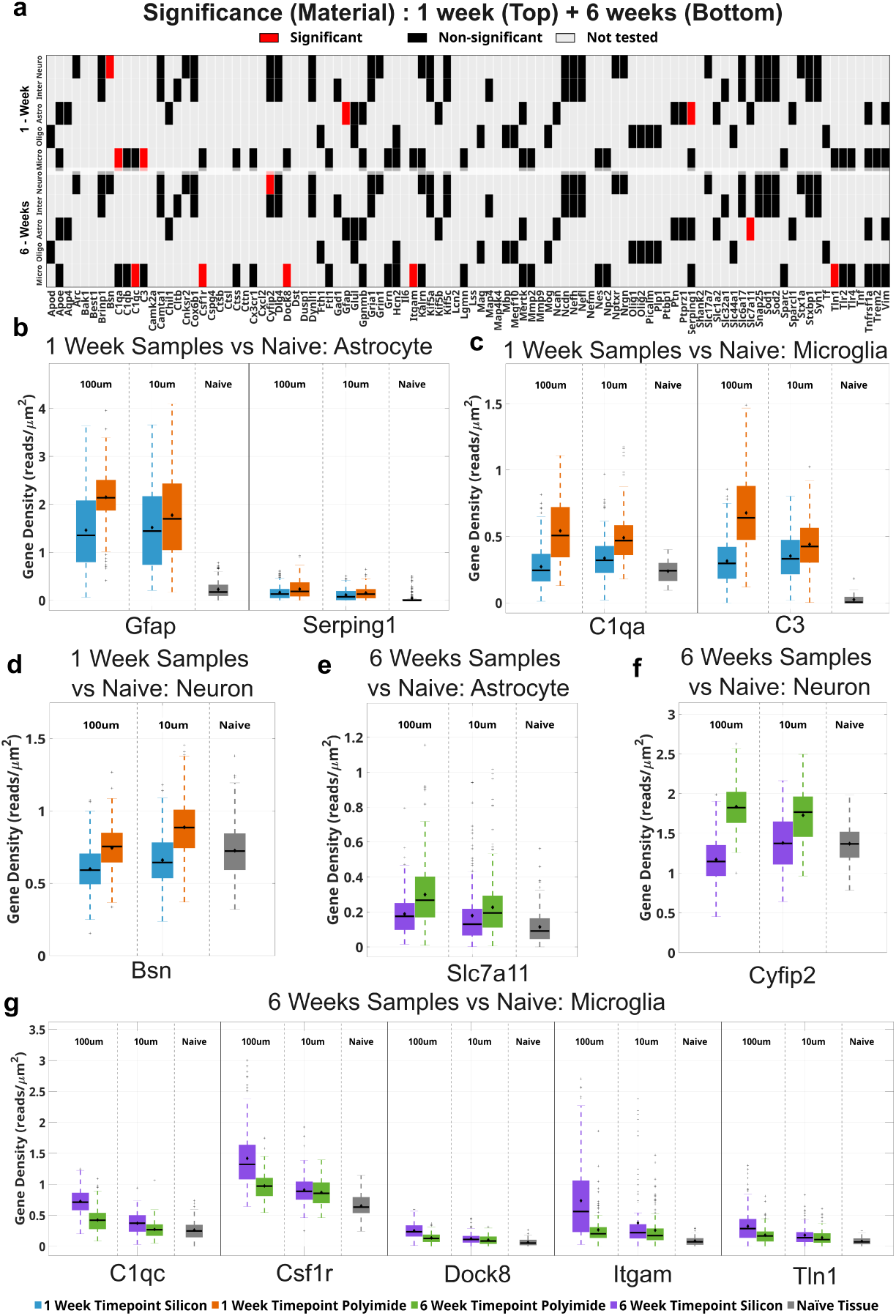
Material-dependent transcriptional responses to implanted microelectrode arrays. (a) Heatmap showing statistically significant material type-driven changes in gene expression at 1 week (top) and 6 weeks (bottom) post-implantation across all cell types relative to naive tissue. Statistical significance was determined using ANOVA with Benjamini–Hochberg FDR correction (adjusted *q <* 0.05). Colors indicate status: red = significant, black = non-significant, grey = not tested due to lack of cell-type expression. (b–c) Astrocyte and microglial gene expression differences at 1 week post-implantation, indicating enhanced early inflammatory signaling around polyimide implants. (d–f) Neuronal and astrocytic material-dependent expression changes at 6 weeks post-implantation. (g) Microglial gene expression differences at 6 weeks post-implantation, high-lighting modest but significant material-specific effects. Boxplots represent normalized gene density (reads/ µm^2^), measured within a 500 *µ*m *×* 500 *µ*m region of interest centered on the implant, with points indicating individual samples. Colors denote implant material and time point. For material comparisons, silicon (n = 6) was compared to polyimide devices (n=5) across different timepoints.

### 3.4 Implant material alters gene expression in single cells in the brain

We contrasted gene expression between silicon and polyimide microelectrode arrays (MEAs) of identical dimensions at each time point to assess material-driven differences at the device interface. In the assessment of global changes in transcription (Fig. 1d), between-material contrasts revealed that at 1 week post-implantation, 10 *µ*m polyimide MEAs elicited a more pronounced upregulation of tissue-response genes than their 10 *µ*m silicon counterparts. These genes encompassed a broad spectrum of effects, including downregulation of neuronal genes and upregulation of genes involved in inflammatory and glial reactivity. These effects were muted by 6 weeks, when the two materials produced very similar results: *Mbp* and *Nefh* were upregulated, and *Stx1a* was downregulated, surrounding small polyimide implants in comparison to silicon. For the 100 *µ*m implants, inflammatory responses were relatively increased surrounding silicon devices in comparison to their polyimide counterparts (upregulation of *C3*, *C1qa-c*, *Lcn2*, *Itgam*, and *Grn*) at 1-week, and no significant differences based on material were evident at 6 weeks. When examining effects at the single-cell level, relatively few genes showed modulated expression as a function of material. At 1-week, on a percell basis, glial reactivity genes were upregulated surrounding polyimide devices (*C1qa*, *C3* in microglia; *GFAP* and *Serping1* in astrocytes). The presynaptic marker Bassoon (*Bsn*) was also upregulated surrounding polyimide devices. At 6 weeks, reactivity markers were more predominant surrounding silicon devices, with per-cell upregulation of microglial genes including *C1qc*, *Csf1r*, *Dock8*, *Itgam*, and *Tln1*. *Cyfip2*, which regulates translation at the synapse and supports synaptic maintenance, was downregulated in excitatory neurons surrounding silicon devices, and *Slc7a11*, which can promote glutamate release [59], was downregulated in astrocytes surrounding silicon devices.

### 3.5 Time point influences gene expression in single cells in the brain

We then evaluated each MEA implant condition across time points to determine whether tissue-response gene expression persists or diminishes as devices remain in the cortex. We first evaluated the global changes in gene expression surrounding devices. In the silicon implant conditions, there were relatively few temporal differences in expression, whereas tissue with polyimide devices showed more pronounced time-dependent changes. For 10 *µ*m silicon implants, contrasts between 6 and 1 week post-implantation revealed differential expression in genes associated with glial activation, oxidative stress, cholesterol homeostasis and cell survival. Relative to 1 week, tissue surrounding 10 *µ*m silicon implants at 6 weeks exhibited significantly reduced expression of *Npc2*, *Csf1r*, *Cx3cr1*, *Grn*, *Trem2* and *Mmp2*; only *Nefm* and *Tf* were upregulated. For 100 *µ*m silicon devices, contrasts between 6 weeks and 1 week post-implantation showed essentially no differences within the selected gene panel, except *Lcn2*, which was significantly upregulated. For 10 *µ*m polyimide devices, the 1-week time point displayed increased expression of a broad range of inflammatory and glial reactivity genes, while neuronal genes were downregulated in expression, in comparison to the 6-week time point. For larger polyimide devices, *Fth1*, *Trem2*, *Cx3cr1*, *C3*, and *C1qc* were downregulated at 6 weeks in comparison to 1-week, while *Lss* was the only upregulated gene in the comparison.

Effects on a single-cell basis displayed both overlapping and distinct effects in comparison to global effects, implying an influence of cell number and/or long-range projections (in the case of neurons) in discrepancies. At 6-weeks in comparison to 1-week, individual glial cells displayed increased expression of reactivity genes, including *Serping1* in microglia and *Gpnmb* in astrocytes. The single largest magnitude effect in this comparison was a downregulation of the *Glul* gene in astrocytes, which encodes the glutamine synthetase enzyme, which otherwise plays a protective role against excessive extracellular glutamate [60]. These effects indicate progressive heightening of glial reactivity over time within single cells, particularly in astrocytes, between the 1 and 6-weeks time points. However, the microglial markers *Csf1r* and *Trem2* were decreased in expression over time: this pattern supports a shift from initial microgliosis to astrogliosis over time, despite the overall appearance of glial scar consolidation with time. Neurons and oligodendrocytes displayed unique effects when comparing the 6-week and 1-week time points. A large reduction in *Fth1*, which encodes the heavy subunit of the iron-binding protein ferritin, occurred in oligodendrocytes. Ferritin reduction in oligodendrocytes is associated with cellular dysfunction and impaired myelination [61]. Oligodendrocytes exhibited increased expression of *Tf* and *Apod*, which were relatively small effects in magnitude. Both excitatory and inhibitory neurons displayed increased *Sod2* expression, indicating engagement of a protective mechanism to neutralize damaging reactive oxygen species. Synaptic and axonal transport markers were also increased (*Syn1* and *Dlg4* in excitatory neurons; *Kif5c* in both neuronal classes). Thus, whereas global effects on gene expression, which include the effect of cell number, indicate a reduction in the inflammatory state of glial reactivity between the 1- and 6-week points, individual glia show some evidence of a progressively more inflammatory state that is accompanied by signs of a recovery of neuronal function.

### 3.6 Summary of single-cell analysis

We examined the effects of implant material and size on the transcriptional tissue response within individual cell types at the device interface using our custom software to identify individual cell types and quantify transcriptional expression from within the nucleus of excitatory neurons, interneurons, astrocytes, microglia, and oligodendrocytes at the device interface. The summary of the results is given in Fig. 8. Our analysis indicates that much of the tissue response appears to be primarily driven by the presence of an indwelling device, with relatively few changes in the transcriptional response attributable to device material or size. In the context of electrode material and size, the most differential gene expression was observed at the 6-week time point. Normalized gene density based boxplots across distinct cell types varied based on implant material, time point, and distance from the device are given in the supplementary figure 31).

**Fig. 7.**
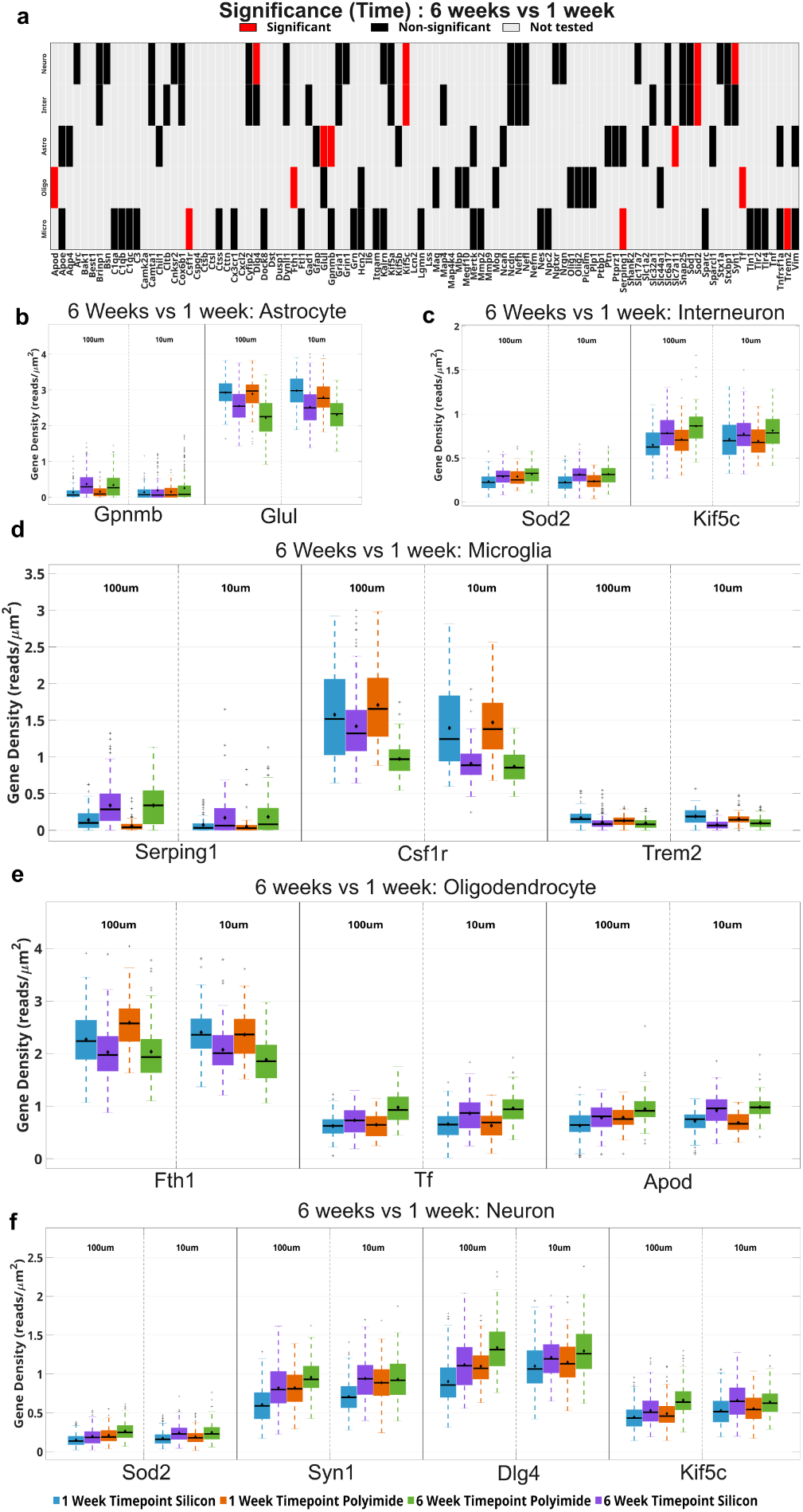
Temporal evolution of cell-type-specific gene expression at the electrode interface. (a) Heatmap illustrating genes that are differentially expressed at 6 weeks compared to 1 week post-implantation across all implant conditions (grey bars = not tested due to lack of cell-type expression, black bars = not significant, red = significant). Statistical significance was determined using ANOVA with Benjamini–Hochberg FDR correction (adjusted *q <* 0.05). (b–d) Astrocyte and microglial temporal expression changes. (e) Oligodendrocyte temporal expression changes. (f) Neuronal temporal expression changes at 6 weeks relative to 1 week post-implantation. Normalized gene density values are shown; statistical significance was determined after false discovery rate (FDR) correction. Comparisons were performed between independent animals. Sample sizes: 1-week polyimide (n = 2), 1-week silicon (n = 3), 6-week polyimide (n = 3), 6-week silicon (n = 3).

**Fig. 8.**
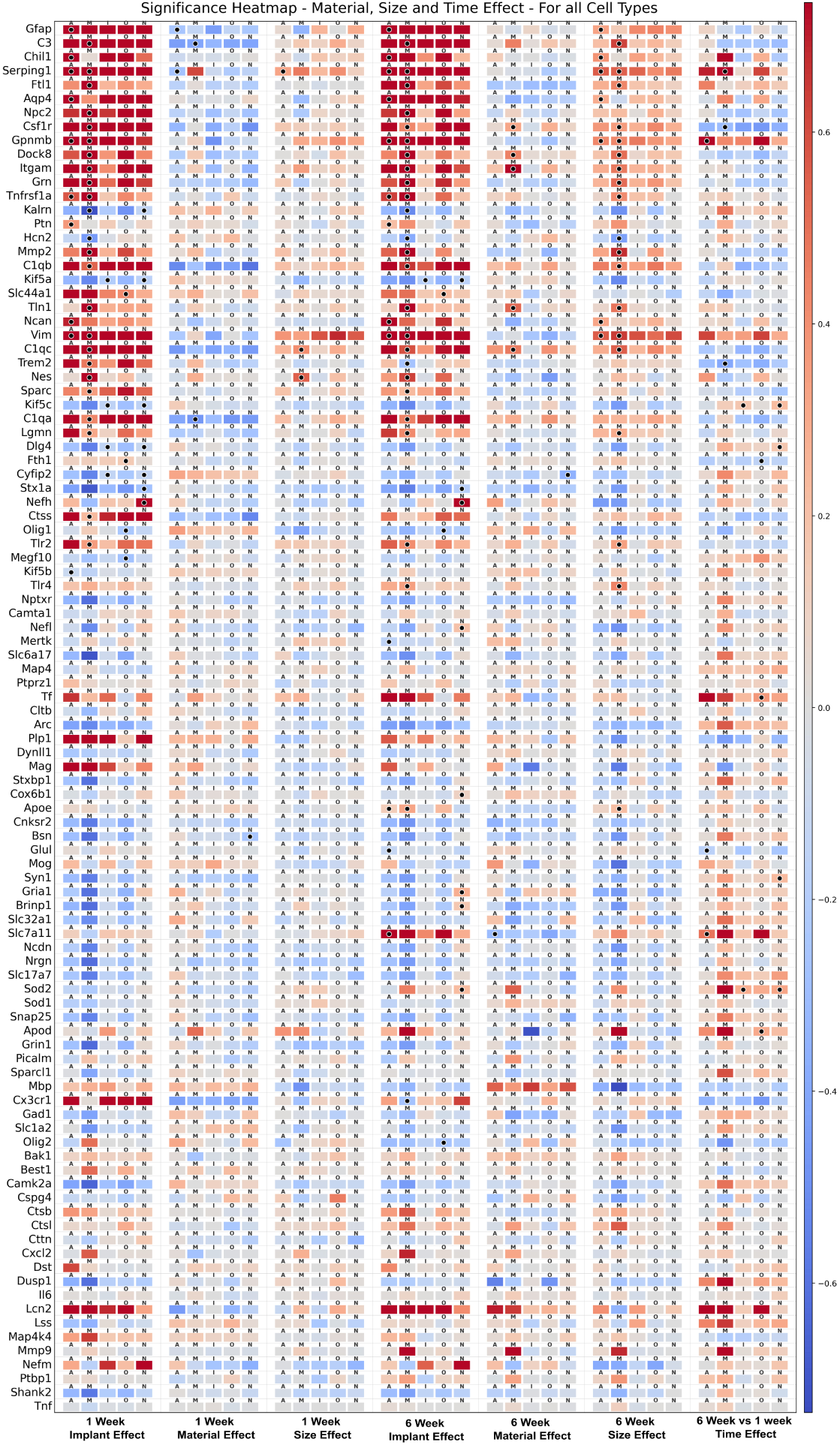
Heatmap showing implant, material, size, and time effects on gene expression across all cell types. Letters above each box indicate cell type (A=Astrocytes, M=Microglia, I=Interneuron, O=Oligodendrocyte, N=Excitatory Neuron). Color intensity represents the relative change in average cell expression score (calculated as the ANOVA-derived estimate normalized by baseline control expression). Estimates were derived using analysis of variance (ANOVA) as described in 2.11, evaluating the concurrent effects of device presence, material (Silicon vs. Polyimide), and size (big vs. small) at 1-week and 6-week timepoints, as well as the effect of time (6-weeks vs. 1-week) for implanted devices. *n* = 2 to 3 animals per condition; single-cell expression values were averaged per Region of Interest (ROI) within each animal before running the ANOVA. Black dots denote statistical significance derived from the ANOVA model (Adjusted *q <* 0.05, corrected via the Benjamini-Hochberg FDR procedure).

### 3.7 Single-cell-resolution gene expression effects as a function of distance from the implant

Next, we selected genes that exhibited the largest material- and size-dependent effects within specific cell types: microglia (*C3*, *Dock8*, *Grn*, *Itgam*, *Trem2*), astrocytes (*Gpnmb*, *Serping1*), oligodendrocytes (*Hcn2*, *Slc44a1*), excitatory neurons (*Kalrn*, *Lcn2*), and interneurons (*Syn1*). For each gene, we quantified relative expression as a function of distance from the implanted device across all implant conditions (Fig. 9 (a-b)). Distances were measured from the device interface (0 µm) to the furthest analyzed point (375 µm). In microglia, distance-dependent expression profiles were evaluated for *C3*, *Dock8*, *Itgam*, *Grn*, and *Trem2*. Expression of *C3*, *Dock8*, *Itgam*, and *Grn* was highest nearest the implanted microelectrode arrays (MEAs). *Dock8* and *Itgam* showed pronounced elevation within 100 µm of the device interface, while *C3* remained strongly expressed out to approximately 250 µm. By contrast, *Trem2* exhibited only modest elevation near the implant relative to distal regions. In astrocytes, *Gpnmb* and *Serping1* expression was markedly increased at the device interface and gradually declined with increasing distance, persisting out to 375 µm. Within oligodendrocytes, *Hcn2* expression was strongly reduced at the device interface and increased monotonically with distance from the implant. Conversely, oligodendrocyte *Slc44a1* expression was elevated nearest the device and decreased as a function of distance. In excitatory neurons, *Kalrn* expression remained relatively consistent out to approximately 225 µm from the device interface. Neuronal *Lcn2* expression was slightly elevated within 100 µm of the implant but fell to near-background levels beyond this distance. Across all samples, the standard error of the distance-dependent analysis indicated consistent gene expression trends (Fig. 9(b)).

**Fig. 9.**
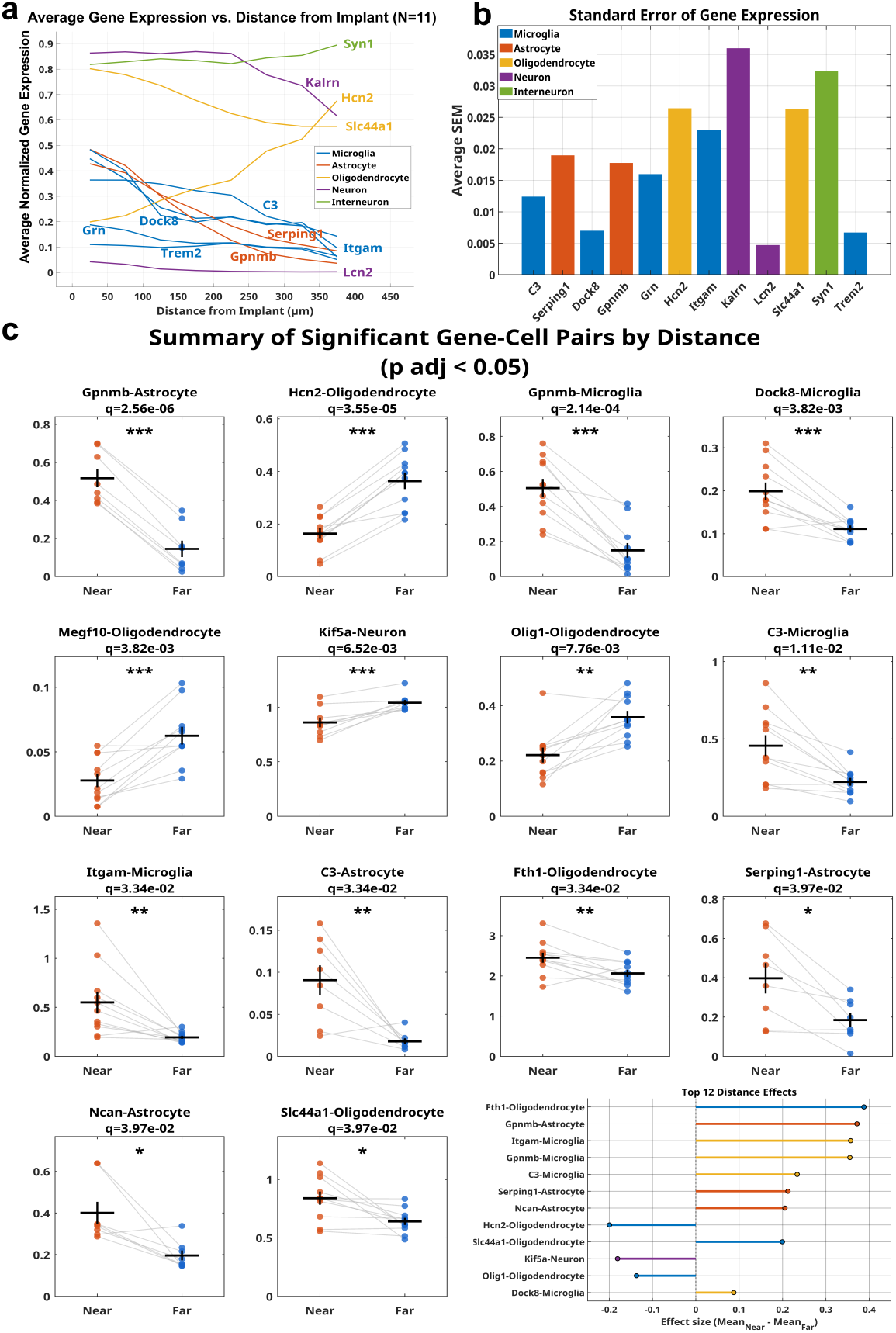
Cell type-specific gene expression as a function of distance from the implant site. Custom analysis pipeline identifies that tissue response genes are spatially expressed around implanted electrodes. (a) Average normalized gene expression as a function of distance from the implant surface (0–350 µm) across all implanted samples (*N* = 11). Neuron-, interneuron-, microglia-, astrocyte-, and oligodendrocyte-associated genes are spatially increased or decreased near the implant or far from the implant. (b) Standard error of spatial gene expression from (a), which illustrates the very small variation across samples. (c) Significant gene-cell pairs illustrate cell-type-specific differences in gene expression in near versus far comparisons using paired *t*-tests. In the figure, the points represent individual samples, and the lines indicate paired measurements. The horizontal lines show mean values. Reported *q*-values reflect false discovery rate (FDR)–adjusted *p*-values (*p*_adj_ *<* 0.05). The bottom-right panel summarizes the top 12 distance-dependent effects ranked by effect size (mean_near_ *−* mean_far_). Positive values indicate increased expression in the near region, whereas negative values indicate enrichment far from the implant.

To further quantify spatial effects, we compared cell-type–specific gene expression averaged across all implanted samples within 60 µm of the device interface (near) to tissue located 150 µm from the implant (far) (Fig.9(c)). These distances were chosen for comparison based on a combination of inspection of the data and reported spatial ranges associated with detection of large-amplitude, well-isolated signals (within approximately 50 µm of the electrode) and non-detection of unit activity above noise (greater than approximately 150 µm from the electrode) [28, 62]. Twelve gene–cell-type combinations exhibited the largest effect sizes in this near-versus-far comparison. In pooled samples, the strongest distance-dependent effects in astrocytes and microglia reflected increased expression of inflammation-associated genes in the near condition. Within microglia, *Gpnmb*, *Dock8*, *C3*, and *Itgam* were significantly upregulated near the implant. In astrocytes, *Gpnmb*, *C3*, and *Ncan* were significantly increased in tissue proximal to the device. Distance-dependent effects in neurons and oligodendrocytes were characterized by reduced expression of genes associated with normal cellular function and homeostasis and increased expression of stress-related genes. In oligodendrocytes, *Hcn2*, *Megf10*, *Olig1* were reduced near implants relative to distal tissue, whereas *Fth1* and *Slc44a1* exhibited increased expression in the near condition. In neurons, *Kif5a* expression was significantly reduced near implants compared to far tissue. Collectively, these results demonstrate that the transcriptomic tissue response to implanted microelectrodes is spatially structured and highly cell-type specific. The observed distance-dependent expression patterns highlight distinct cellular programs that extend beyond the immediate electrode interface and the glial scar, providing insight into how implant-induced molecular perturbations propagate through surrounding cortical tissue. Detailed distance-dependent trend plots for the relative gene expression, including paired t-test comparisons for selected gene-cell type pairs, are provided in the supplementary figures 32-43.

## 4 Discussion

Understanding the tissue response to indwelling electrodes is essential for improving the functional longevity and biocompatibility of neural prostheses. In this study, we have provided new transcriptional analyses of the tissue response to implanted electrodes with single-cell resolution. We used single-cell spatial transcriptomics to investigate hypotheses that MEAs fabricated with smaller feature sizes and softer materials may attenuate the inflammatory tissue response, thus improving the functional longevity of implanted MEAs *in vivo*. We first tested the differential expression of genes within 500 µm of the implant versus unimplanted tissue, and we confirmed the upregulation of inflammatory-associated genes and downregulation of neurotransmission-associated genes surrounding devices, similarly to previous reports [34]. Differential expression analysis indicated that the bulk of the tissue response is primarily driven by the presence of any implant, regardless of bulk material or size, and persists for at least 6-weeks post-implantation.

To statistically test the influence of implants, electrode material, electrode size, and time point on gene expression in single cells, we developed a customized computational analysis pipeline. We used cell type-specific transcripts (marker genes) to identify distinct populations of excitatory neurons, interneurons, astrocytes, microglia, and oligodendrocytes, and the association of specific transcripts with the individual cell types. When observing gene expression changes in individual cell types due to the presence of an implant, we observed increased reactivity genes in astrocytes and microglia that were conserved across time points. We also observed an initial downregulation of synaptic genes in neurons. However, neuronal gene expression shows partial recovery by 6-weeks, albeit in a subset of genes. Plasma levels of neurofilament correlate with the degree of axonal damage following brain injury [63], and the increases in neurofilament gene expression observed here are corroborated by our previous immunohistochemistry studies [64]. Coupled with observable damage to dendritic arbors [38], the per-cell increase in neurofilaments may reflect an internal compensatory mechanism to repair insertional injury. Similarly, the increase in *Cox6b1* and *Sod2* expression in excitatory neurons, which encode anti-apoptotic and reactive oxygen species-neutralizing enzymes [65,66], reinforces the engagement of protective mechanisms in neurons surrounding chronic implants. *Stx1a*, which encodes the Syntaxin-1a synaptic vesicle fusion protein, was consistently decreased in neurons at both time points, and downregulation of the neuron-specific kinesin *Kif5a* was also conserved across time points. Reduced expressions of *Stx1a* and *Kif5a* indicate reduced synaptic transmission and mitochondrial transport in neurons at the electrode interface, consistent with our previous observations of impaired excitatory neurotransmission near implants [38]. In oligodendrocytes, a choline transporter crucial for myelination (*Slc44a1*) was upregulated at both time points, while downregulation of *Olig1* may be consistent with differentiation of progenitors to mature myelinating oligodendrocytes [67, 68]: both of these effects are consistent with neuronal repair. Thus, our data indicate that implants generally cause upregulation of inflammatory genes in astrocytes and microglia that are coupled to damage-initiated losses in synaptic transmission and subsequent engagement of compensatory protective mechanisms to preserve local neurons.

Examination of temporal effects showed that tissue from the 6-week time point revealed fewer differentially expressed genes than the 1-week time point in the global comparison of 500 *µ*m regions of tissue; generally, the bulk analyzed data support a shift toward dampening glial reactivity over time. However, results at the single-cell level revealed more complexity in the underlying mechanisms. Effects that were notably large in magnitude included downregulation of *Glul* in astrocytes and downregulation of *Fth1* in oligodendrocytes. Reduced *Glul* expression is associated with astrogliosis and subsequent neurodegeneration and neuronal hyperexcitability [69].

Although recent seminal studies have begun to explore the impacts of device implantation on oligodendrocytes [70, 71], these cells historically have received less attention in reported literature in comparison to astrocytes and microglia. Oligodendrocytes rely on iron metabolism for cellular maintenance and myelination, and the reduction in *Fth1* over time implies a shift toward dysfunction and loss of myelin [61]. Further, since local glia, and astrocytes in particular, are the primary source of ferritin iron to oligodendrocytes [72], this result implicates astroglial reactivity as a potential source of decreased *Fth1* in oligodendrocytes. Reduced *Fth1* in astrocytes results in hypomyelination and, more broadly, oxidative stress in the brain [72]. In parallel, our results show that neurons upregulate superoxide dismutase, a defense mechanism against oxidative stress, over time following implantation. Taken together, our data indicate that the initial activation of microglia is followed by astrocyte activation (in accordance with previous reports) [29], but although glial reactivity begins to consolidate spatially around the device by 6-weeks, transcriptional data reveal a progressive increase in reactivity in individual astrocytes over time. Iron metabolism in oligodendrocytes, which relies on astroglial delivery of ferritin iron, is impaired, while compensatory mechanisms are activated in neurons to counteract a state of oxidative stress [31]. The increased expression of synaptic machinery in neurons *(Syn1*, *Dlg4*), and increased expression of *Slc7a11* in astrocytes, which can promote glutamate release in response to oxidative stress [73], are both indicative of an engagement of recovery mechanisms following the neuronal damage and dysfunction that occur post-implantation [38].

Device effects on astrocytes, oligodendrocytes, and neurons occur in parallel to a relatively large reduction in the expression of *Csf1r* in microglia at 6-weeks, supporting a gradual reduction in the inflammatory state of these cells. Inhibition of *Csf1r* is associated with reduced neuronal degeneration [69]. Signs of neuronal recovery at 6-weeks coincide with our prior observations of changes in intrinsic excitability and a reduction of spontaneous excitatory postsynaptic currents in individual neurons at the device interface at that time point [38]. Thus, while the gene expression data show a shift toward reparative mechanisms at the 6-week time point, the increase in the expression of synapse-associated genes occurs at a time point when structural damage and functional deficits are evident in neurons. Based on our results, we infer that at the later time points, a reduced inflammatory state in microglia is associated with a minor recovery in synapse-associated genes in neurons, but that the heightened reactivity of astrocytes coupled with oligodendrocyte dysfunction produces new challenges to neuronal health and function in the chronic implant phase.

Design features influenced gene expression more predominantly at the tissue level than on a single cell basis, meaning that the underlying mechanisms driving improved tissue integration of smaller/more flexible devices are more closely related to changes in the numbers of cells than per-cell gene expression changes. An associated reduction in insertion associated damage to vasculature and a reduced release of activating blood-derived factors are plausible explanations for the source of the effect on glial activation and proliferation [74]. However, there were a few notable effects at the single-cell level. At the 1-week time point, larger devices were associated with mildly increased expression of glial reactivity genes (e.g., complement *C1qc* in microglia, and *Serping1* in astrocytes). By the 6 week time point, many more reactivity genes, particularly those associated with astrocytes (*Aqp4*, *Chil1*, *GFAP*, *Gpnmb*, *Ncan*, *Serping1*, *Vim*) were upregulated surrounding larger implants. Thus, larger implants seem to exacerbate astroglial reactivity in individual cells over time, even as the glial scar consolidates spatially around the implant. This observation would seem to implicate non-insertion damage-related mechanisms. The adherence of macrophages to material fibers *in vitro* has been observed to increase with increasing dimensions (12 to 25 microns in diameter, above and below the 15 micron threshold for phagocytic activity) [75]. Taken together, the data are potentially explainable by the idea that devices of larger dimensions promote increased numbers of reactive cells via blood-derived factors driving proliferation, followed by an increasingly reactive state of glia at the single cell level promoted by adherence-related mechanisms and recognition of the foreign body. No size effects were evident in individual neurons or oligodendrocytes, indicating that the influence of dimensions on biocompatibility is astrocyte and microglia-centric and is relatively decoupled with any coordinated influence on neuronal structure, function, re-myelination, and/or other repair mechanisms.

Material effects were intertwined with size at the 1-week time point and were likely largely influenced by the extent of insertional damage for the small polyimide implants. Material effects were generally absent by the 6-week time point: on a per-cell basis in the chronic injury phase, our data indicate that this design feature had an unremarkable influence on tissue integration. These results are reinforced by our previous electrophysiological and morphological assessments at the single cell level [38], which showed very similar effects irrespective of electrode material. Although these results may appear to be antithetical to the hypothesis that more flexible polymer materials improve brain tissue integration [39, 76], our results can be reconciled with prevailing trends in design if both size and material modulus are considered: A coordinated reduction in size alongside increased material flexibility may be required to reduce the overall bending stiffness and support integration [41].

The data collected from this study illustrate new single-cell resolution transcriptomics surrounding implanted electrodes of varying materials and dimensions. In situ spatial transcriptomics paired with our custom analysis has enabled the spatiotemporal quantification of gene expression within individual cell types in intact tissue sections implanted with microelectrodes. Analysis of the tissue response to implanted electrodes identified that many genes within our custom panel are differentially expressed within discrete cell types in response to electrode implantation. We observed a progressive heightening of the reactive state of individual astrocytes over time against dampening inflammation in individual microglia, the emergence of specific reparative mechanisms in oligodendrocytes and neurons, and the relationship between device features and tissue versus single-cell-level effects. These observations provide new mechanistic insights into device-tissue interactions in the brain at the single-cell level.

## 5 Code Availability

Custom MATLAB scripts used for region selection, transcript density estimation, marker-based cell-type inference, distance-to-implant computation, and summary plotting are available on GitHub at the following location: https://github.com/msureil/ SpatiaXen-Analysis.

## 6 Data Availability

The datasets supporting the results of this study are available upon contacting the authors.

## 7 Author contributions

**Cort Thompson** and **Anirban Chakraborty** contributed equally to this work. **Cort Thompson**: Conceptualization, Methodology, Formal analysis, Visualization, Writing – original draft. **Anirban Chakraborty**: Methodology, Investigation, Formal analysis, Visualization, Data analysis and Results generation, Writing – original draft. **Lauren Wade-Kleyn**: Investigation, Data curation, Proofreading. **Mark Reimers**: Conceptualization, Methodology, Data Analysis, Supervision, Writing – review & editing. **Erin Purcell**: Conceptualization, Supervision, Funding acquisition, Project administration, Writing – original draft, Writing – review & editing.

## 9 Acknowledgements

The authors would like to thank the University of Michigan Genomics Core for their assistance with Xenium processing. We are grateful to Dr. Melinda Frame for her technical support with microscopy. We gratefully thank Mohammad Ghajar and Dr. John Seymour (University of Texas Health Science Center and Rice University; Houston, TX) for fabricating the custom polyimide devices used in this study. This work was supported by an NSF CAREER award (PI: Purcell, #1943716).

