## Supplementary File 1 for "A systematic analysis of brain tissue response to microelectrode material and size with single-cell spatial transcriptomics"

### Supplementary File 01

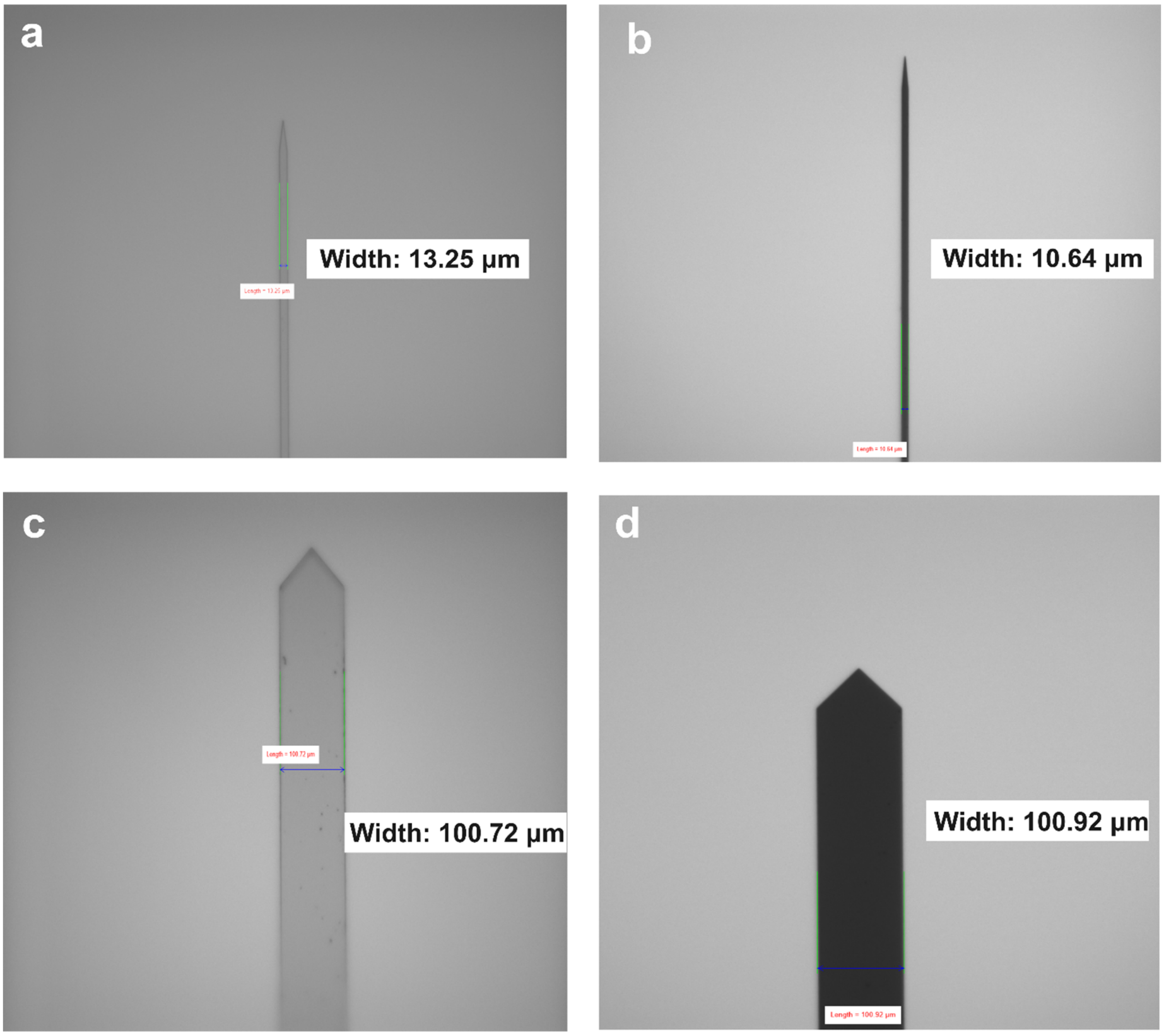


**Supplementary Figure 1:** Representative bright field images of implants used for this study prior to shuttling with polyethylene glycol. (a) PI10, (b) SI10 (c) PI100 and (d) SI100 implants.

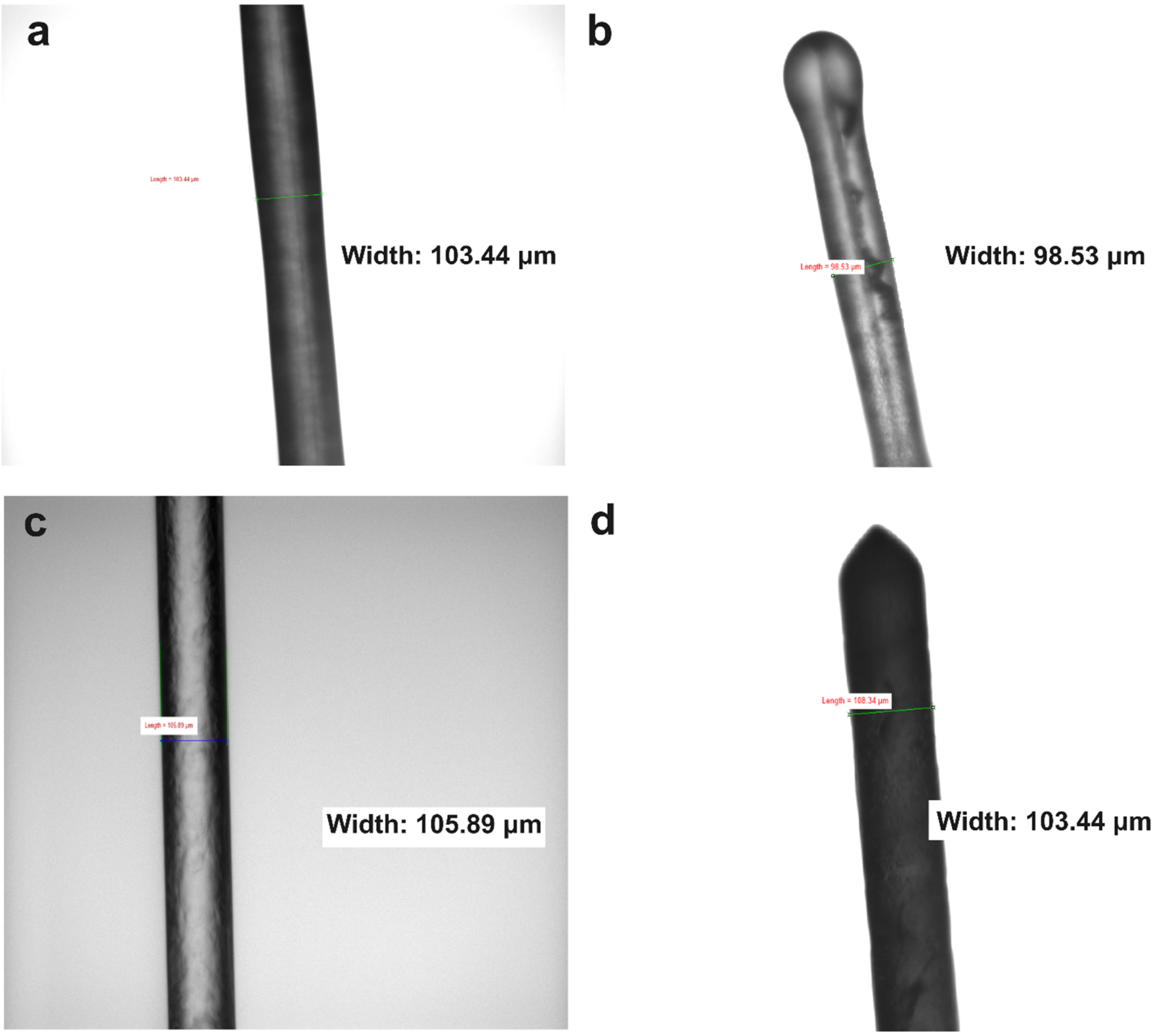


**Supplementary Figure 2:** Representative bright field images of implants used for this study after shuttling with polyethylene glycol. (a) PI10 , (b) SI10 , (c) PI100 , (d) SI100 implants.

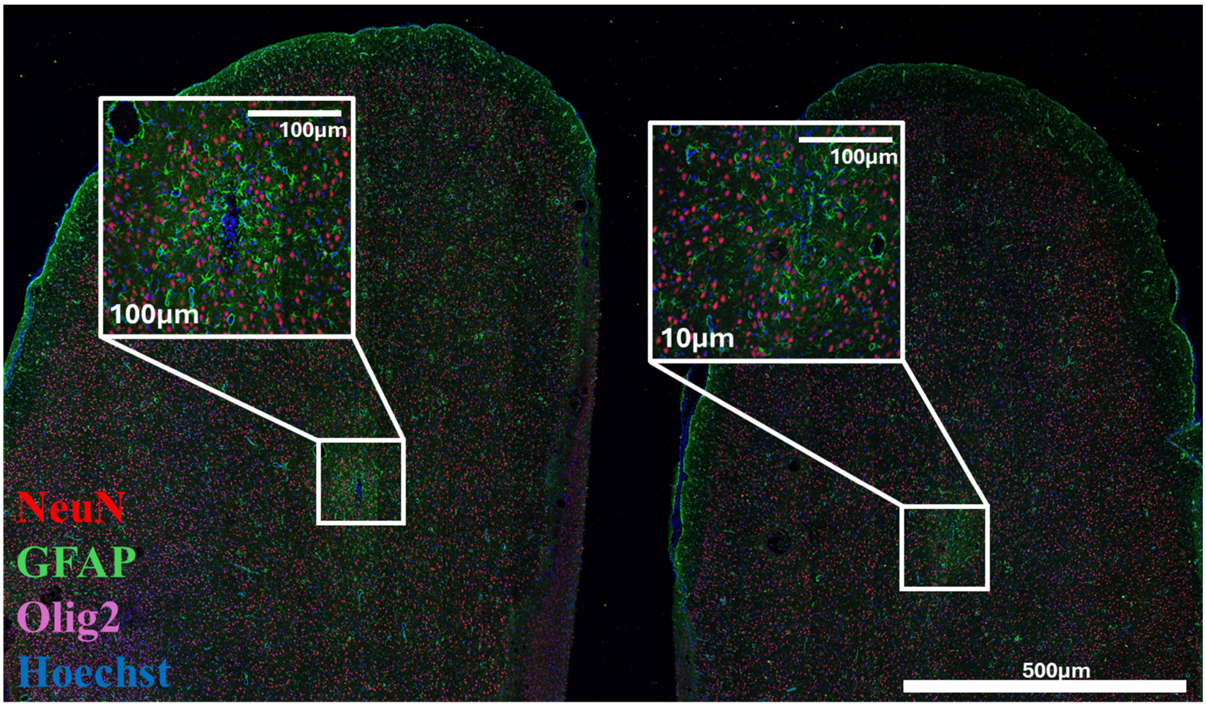


**Supplementary Figure 3:** Representative whole-section immunofluorescence from a 1-week sample with bilateral silicon probe injuries (left: 100 µm, right: 10 µm).

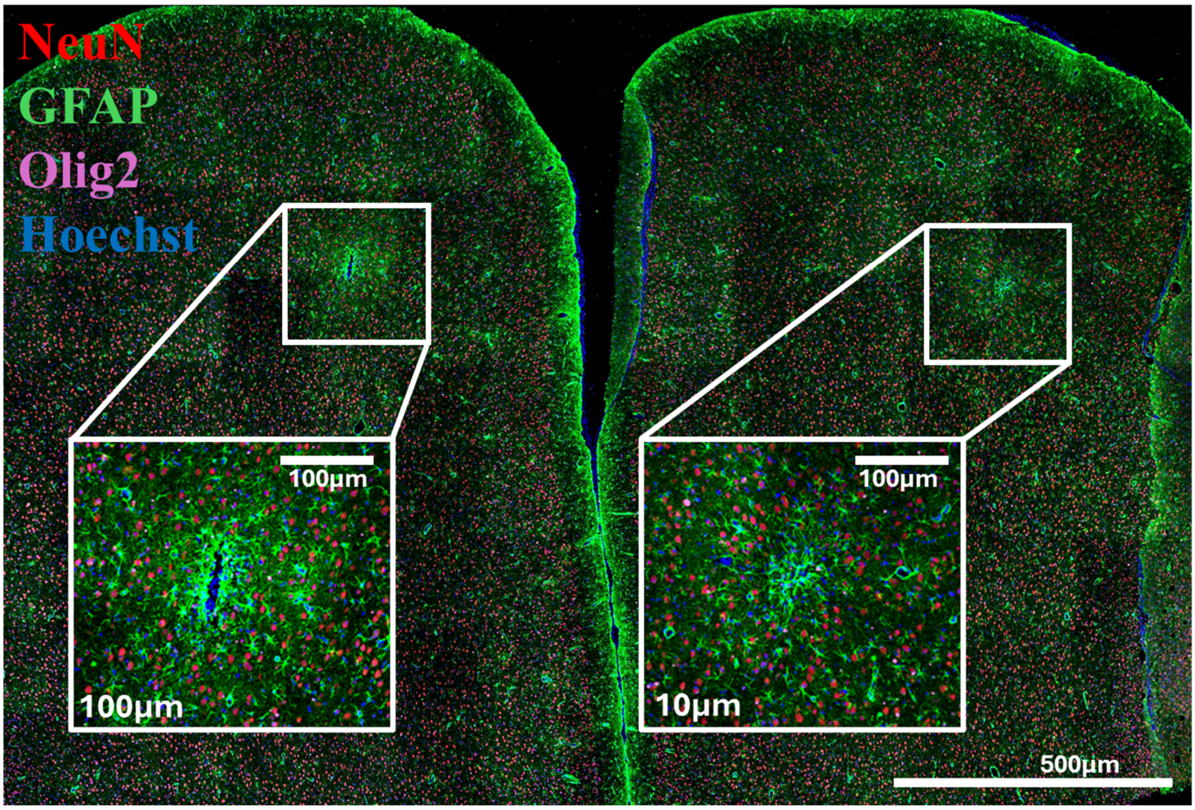


**Supplementary Figure 4:** Representative whole-section immunofluorescence from a 1-week sample with bilateral polyimide probe injuries (left: 100 µm, right: 10 µm).

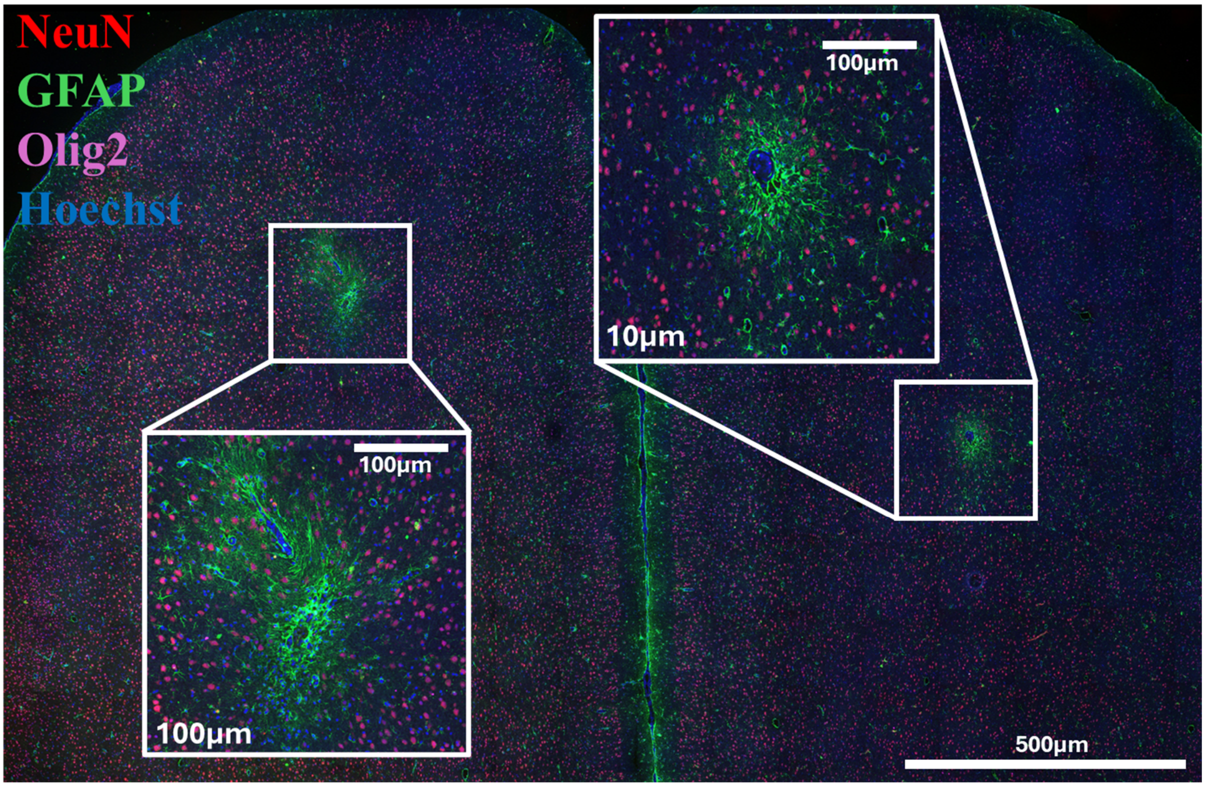


**Supplementary Figure 5:** Representative whole-section immunofluorescence from a 6-week sample with bilateral silicon probe injuries (left: 100 μm, right: 10 μm).

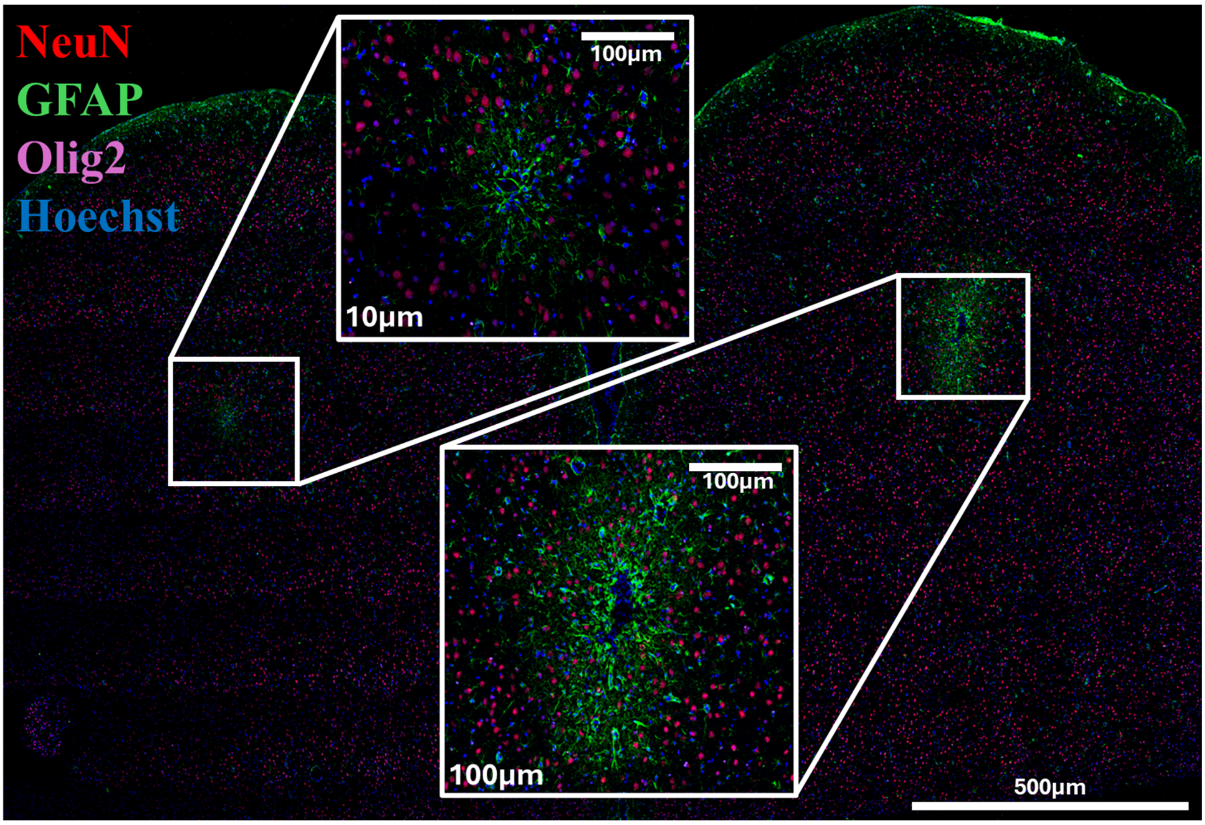


**Supplementary Figure 6:** Representative whole-section immunofluorescence from a 6-week sample with bilateral polyimide probe injuries (left: 100 µm, right: 10 µm).

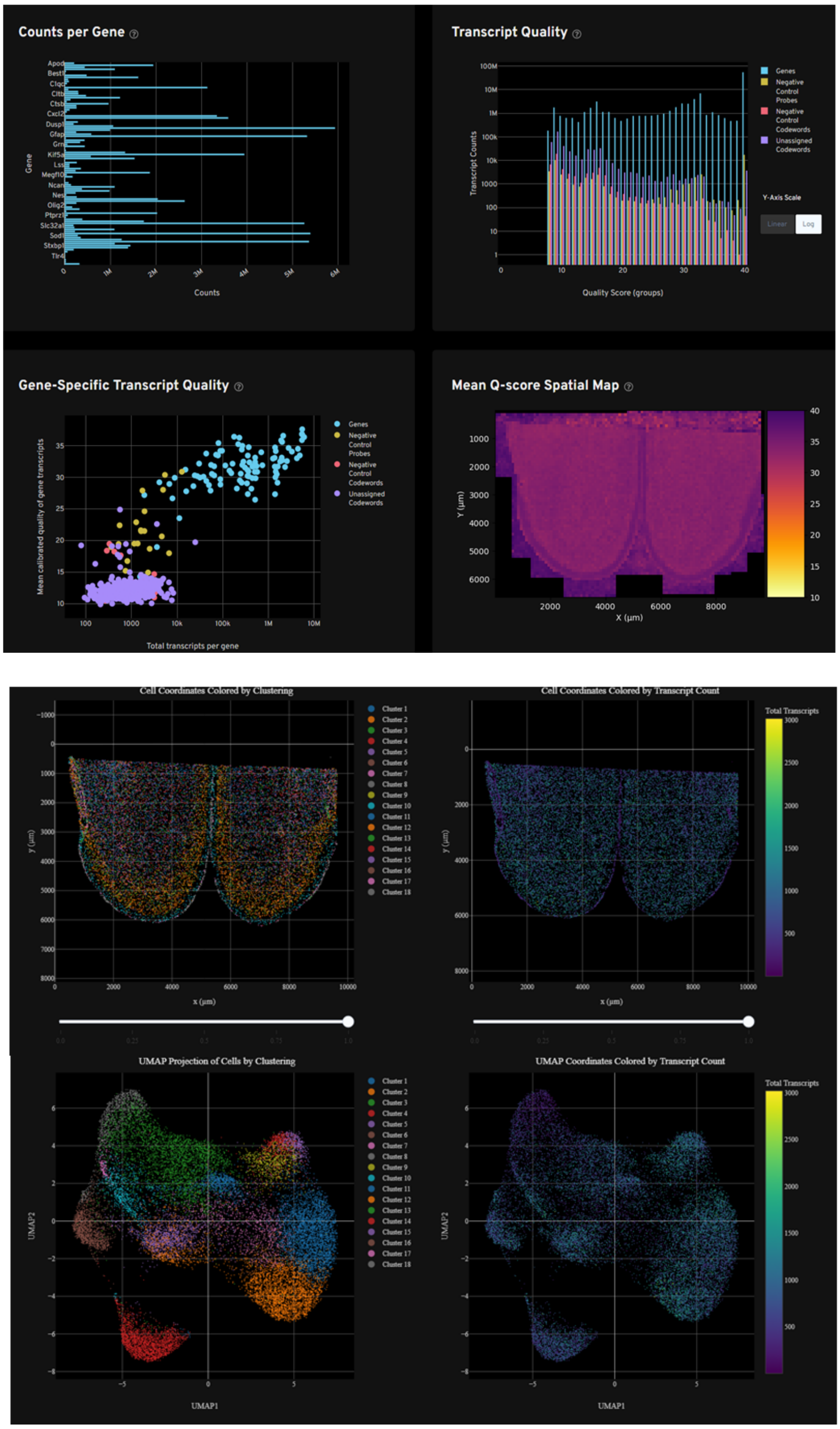


**Supplementary Figure 7:** Quality control analysis from the first naïve sample (ROI A) on one xenium slide. Quality control analysis indicates counts per gene, transcript quality, mean Q-score, cellular clusters, counts per cell assigned to spatial coordinates, and graph-based cluster UMAPs organized by clusters and transcript counts.

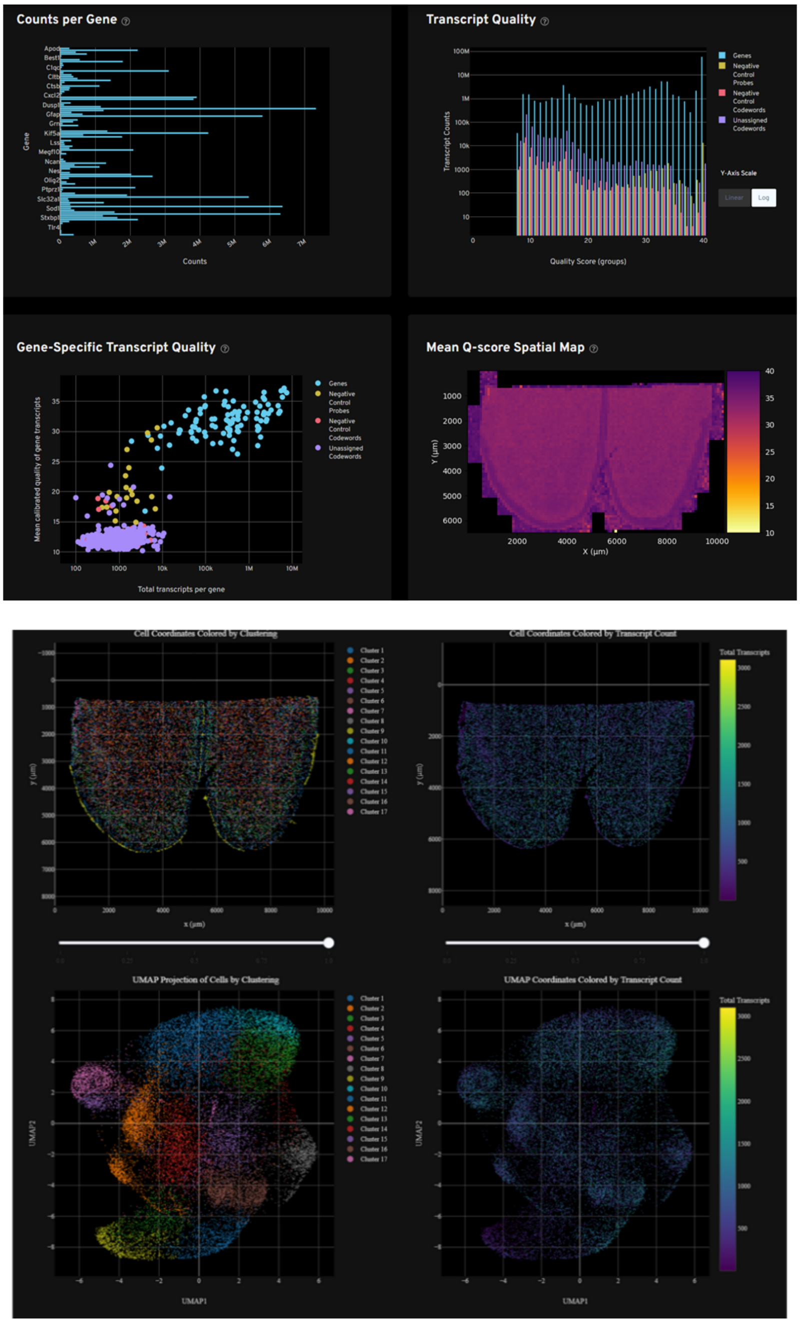


**Supplementary Figure 8:** Quality control analysis from the first naïve sample (ROI B) on one xenium slide. Quality control analysis indicates counts per gene, transcript quality, mean Q-score, cellular clusters, counts per cell assigned to spatial coordinates, and graph-based cluster UMAPs organized by clusters and transcript counts.

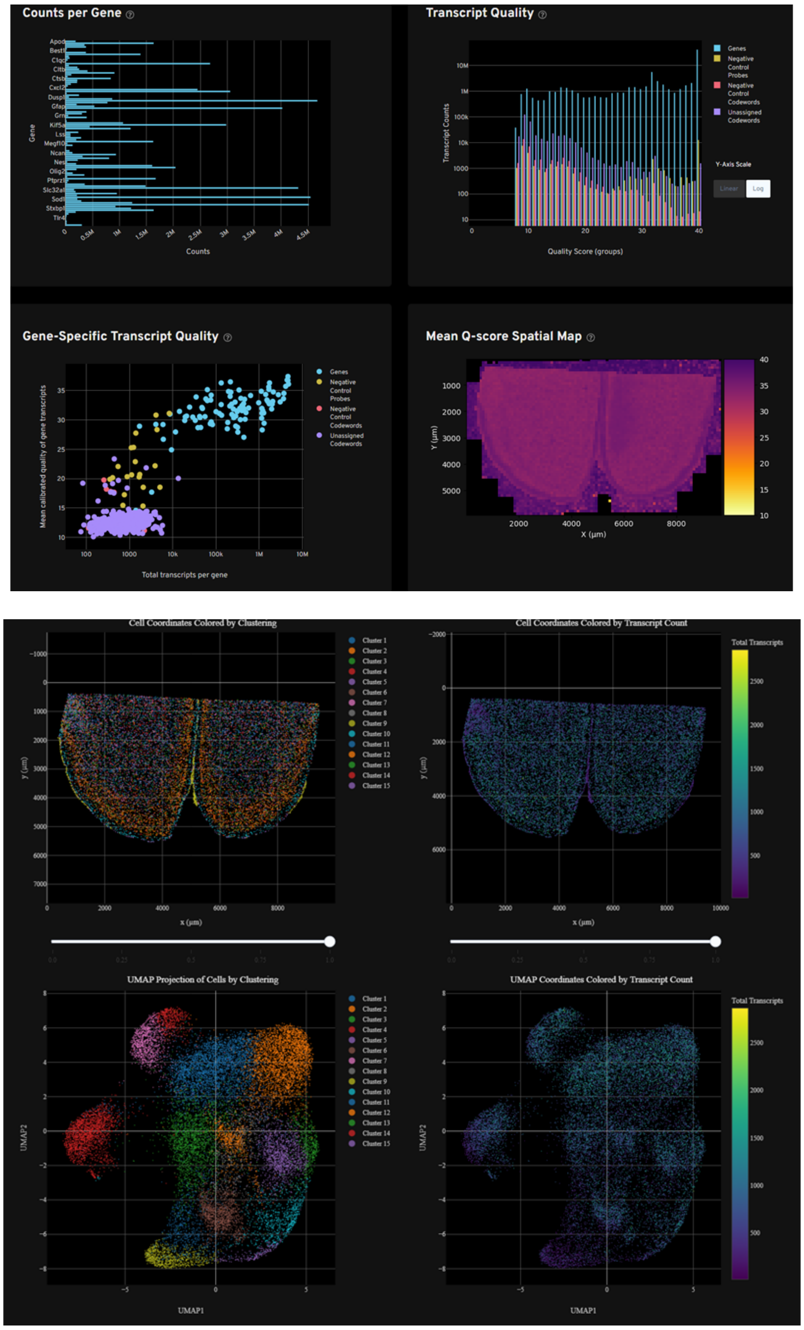


**Supplementary Figure 9:** Quality control analysis from the first naive sample (ROI C) on one xenium slide. Quality control analysis indicates counts per gene, transcript quality, mean Q-score, cellular clusters, counts per cell assigned to spatial coordinates, and graph-based cluster UMAPs organized by clusters and transcript counts.

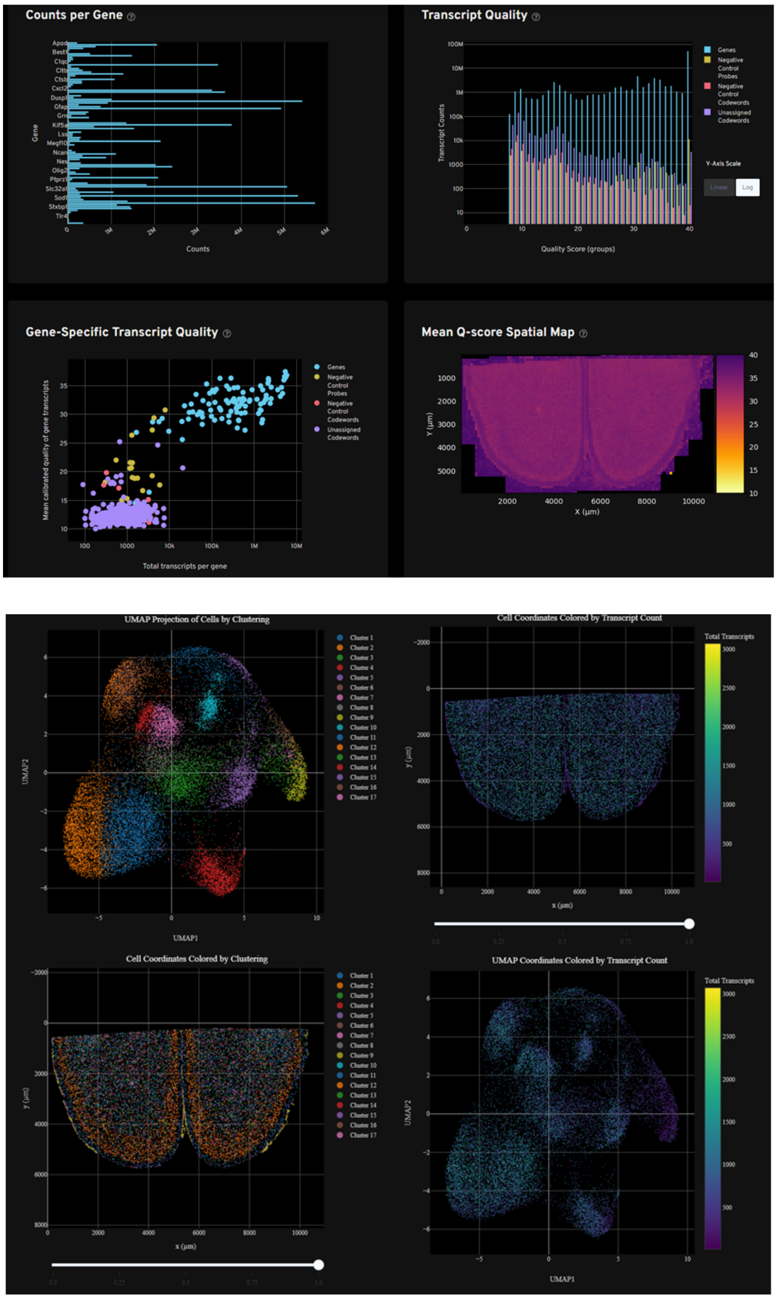


**Supplementary Figure 10:** Quality control analysis from the first bilateral silicon sample (ROI A) on one xenium slide. Quality control analysis indicates counts per gene, transcript quality, mean Q-score, cellular clusters, counts per cell assigned to spatial coordinates, and graph-based cluster UMAPs organized by clusters and transcript counts.

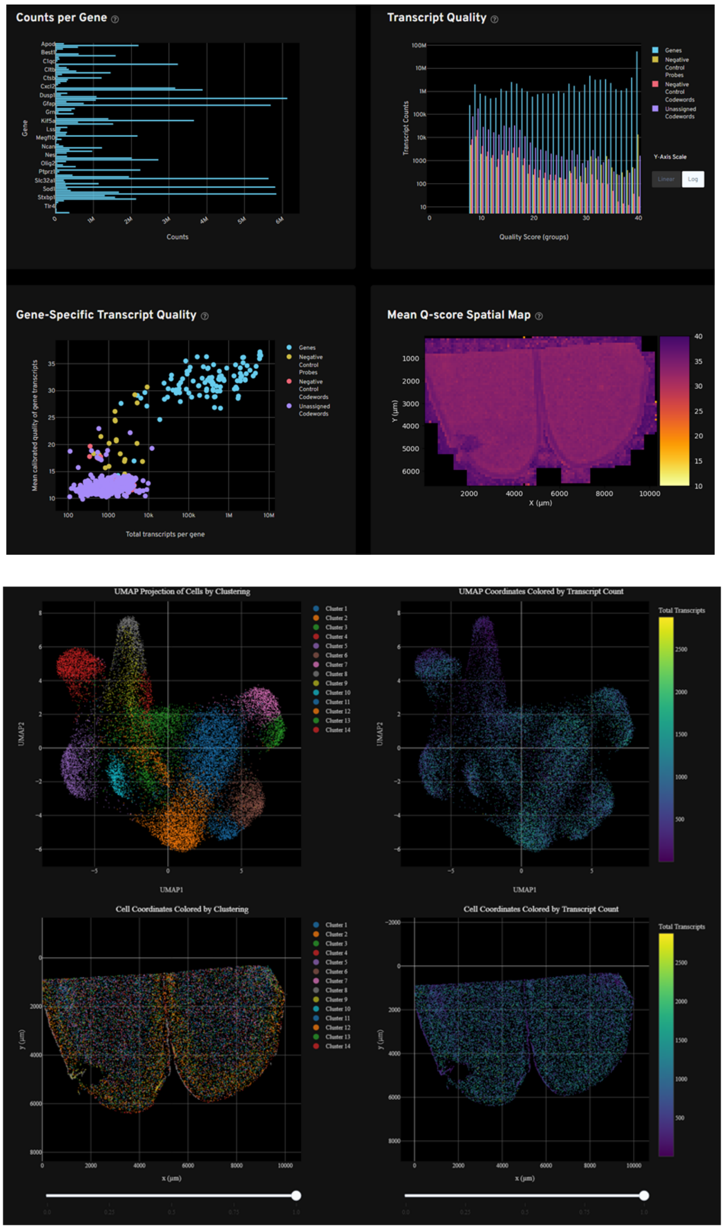


**Supplementary Figure 11:** Quality control analysis from the second bilateral silicon sample (ROI B) on one Xenium slide 1-week post-implantation. Quality control analysis indicates counts per gene, transcript quality, mean Q-score, cellular clusters, counts per cell assigned to spatial coordinates, and graph-based cluster UMAPs organized by clusters and transcript counts.

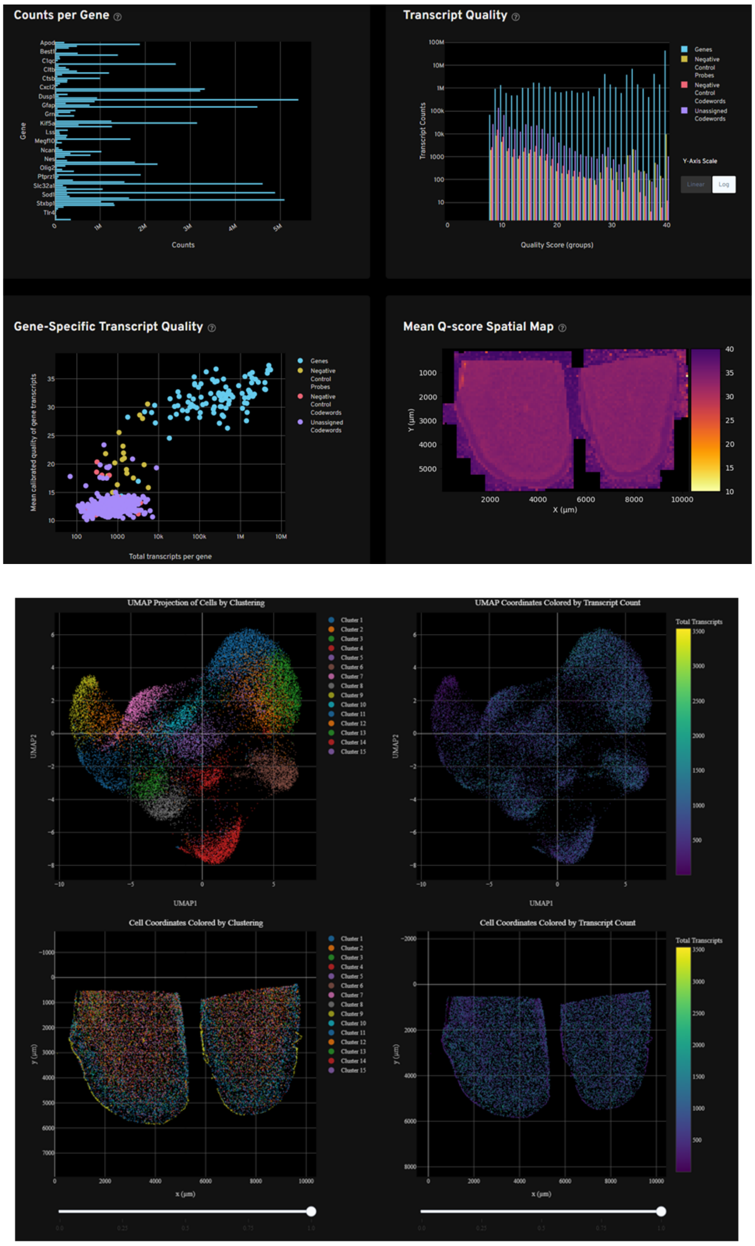


**Supplementary Figure 12**: Quality control analysis from the third bilateral silicon sample (ROI C) on one xenium slide 1-week post-implantation. Quality control analysis indicates counts per gene, transcript quality, mean Q-score, cellular clusters, counts per cell assigned to spatial coordinates, and graph-based cluster UMAPs organized by clusters and transcript counts.

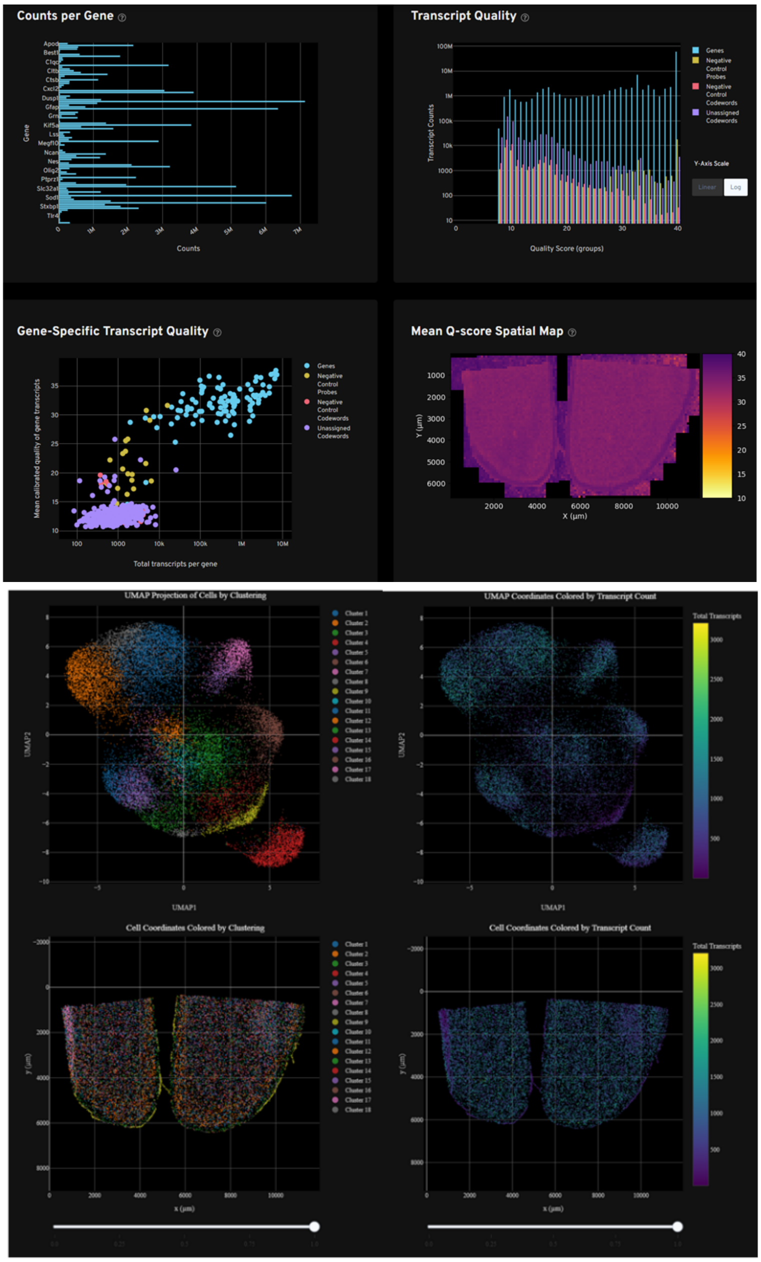


**Supplementary Figure 13:** Quality control analysis from the first bilateral polyimide sample (ROI B) on one Xenium slide 1 week post-implantation. Quality control analysis indicates counts per gene, transcript quality, mean Q-score, cellular clusters, counts per cell assigned to spatial coordinates, and graph-based cluster UMAPs organized by clusters and transcript counts.

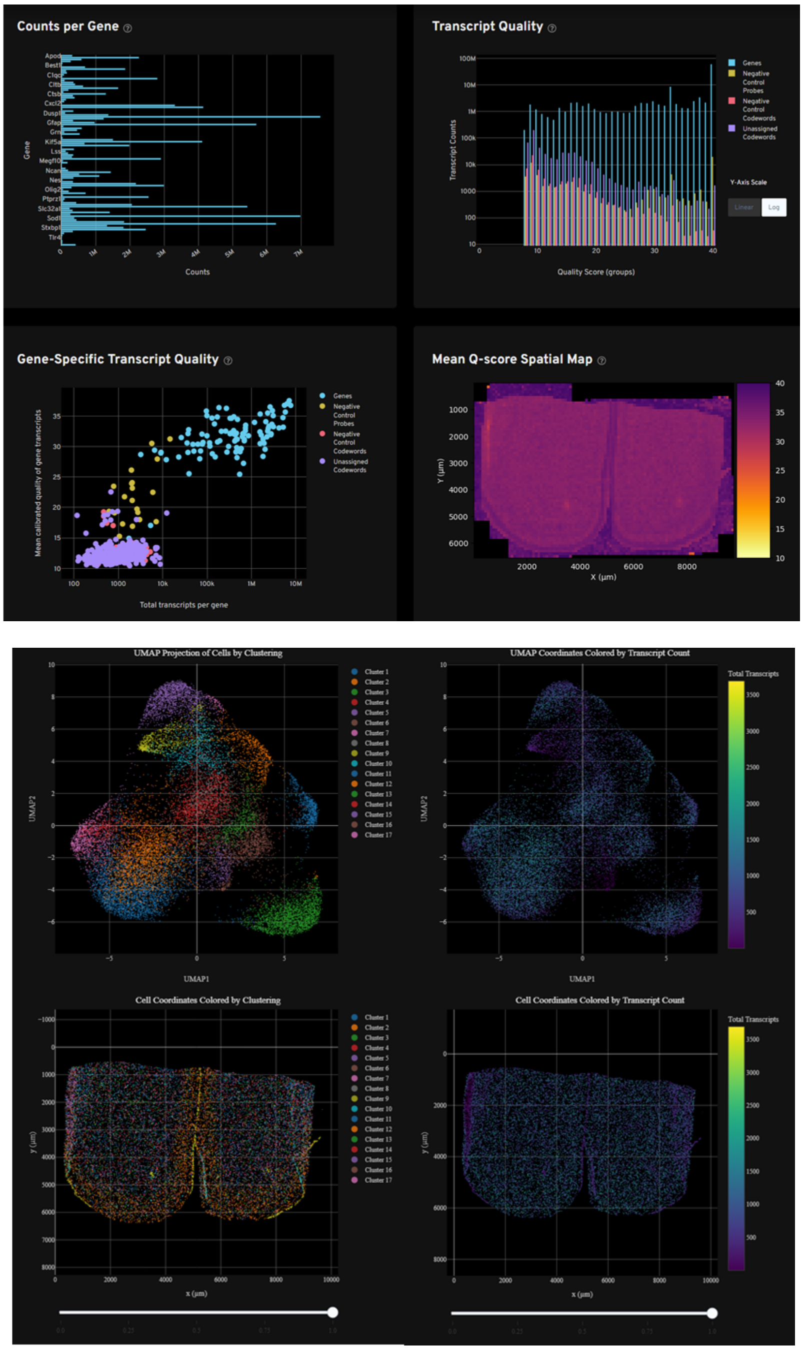


**Supplementary Figure 14:** Quality control analysis from the second bilateral polyimide sample (ROI C) on one xenium slide 1-week post-implantation. Quality control analysis indicates counts per gene, transcript quality, mean Q-score, cellular clusters, counts per cell assigned to spatial coordinates, and graph-based cluster UMAPs organized by clusters and transcript counts.

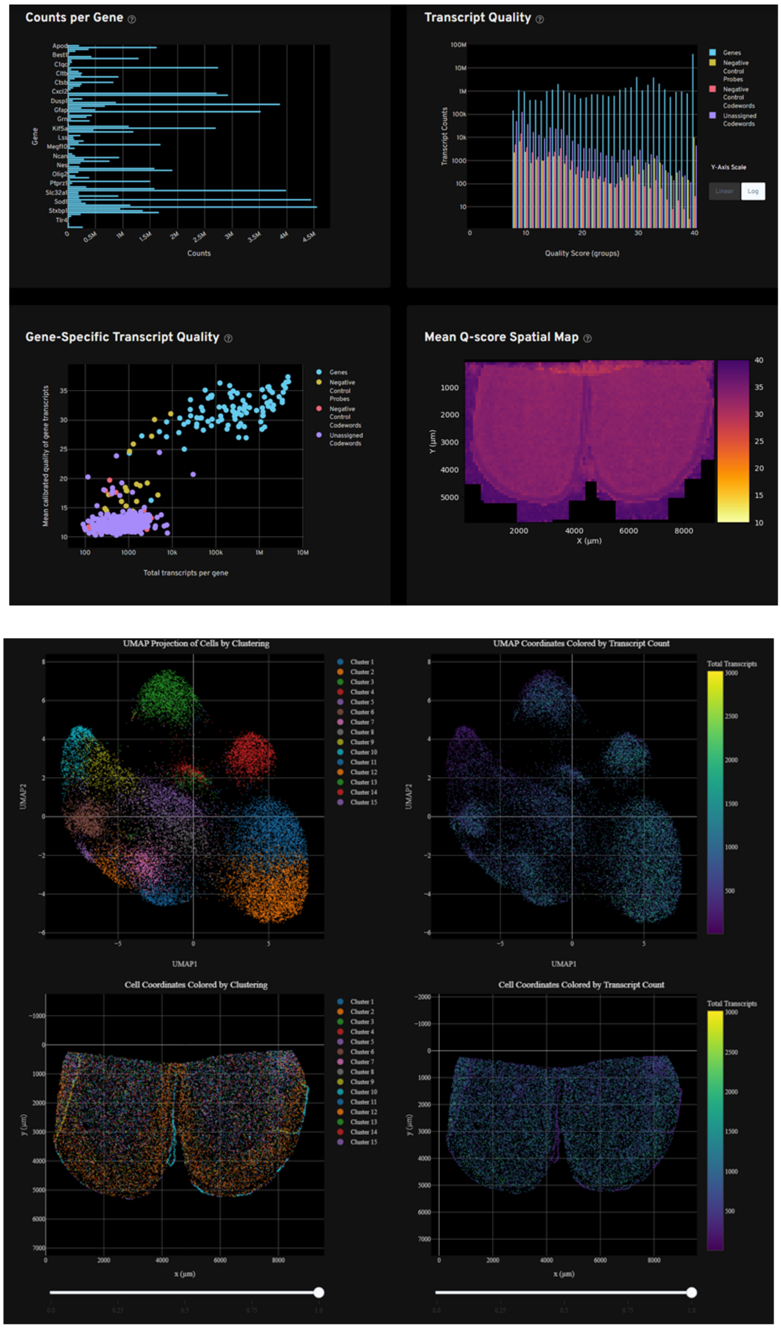


**Supplementary Figure 15:** Quality control analysis from the first bilateral silicon sample (ROI A) on one Xenium slide 6-weeks post-implantation. Quality control analysis indicates counts per gene, transcript quality, mean Q-score, cellular clusters, counts per cell assigned to spatial coordinates, and graph-based cluster UMAPs organized by clusters and transcript counts.

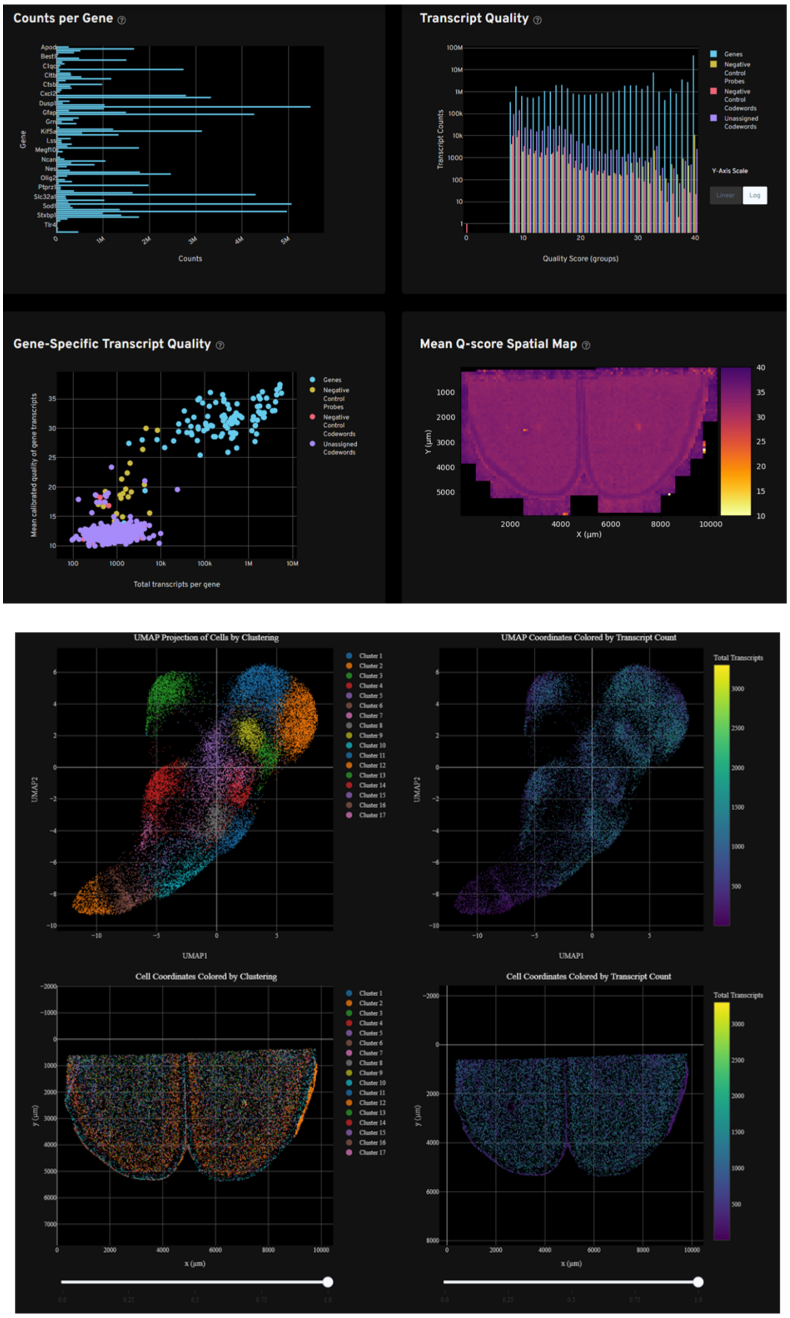


**Supplementary Figure 16:** Quality control analysis from the second bilateral silicon sample (ROI B) on one Xenium slide 6 weeks post-implantation. Quality control analysis indicates counts per gene, transcript quality, mean Q-score, cellular clusters, counts per cell assigned to spatial coordinates, and graph-based cluster UMAPs organized by clusters and transcript counts.

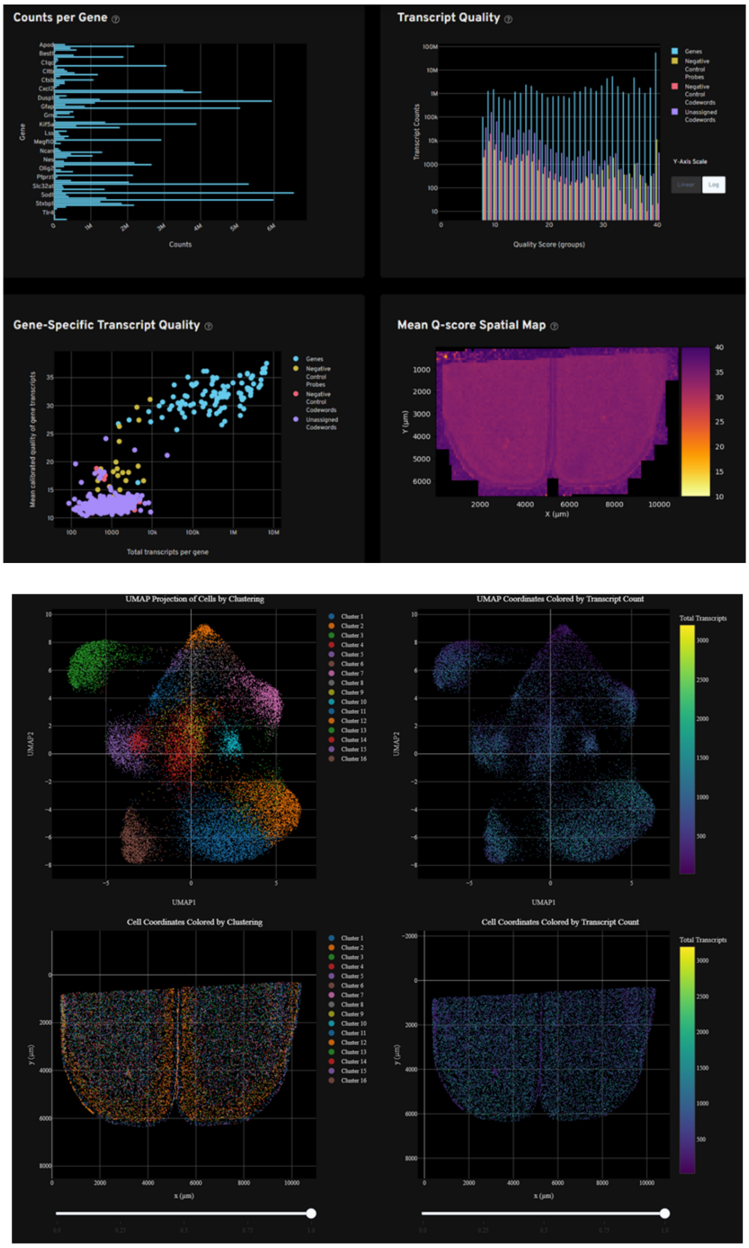


**Supplementary Figure 17:** Quality control analysis from the third bilateral silicon sample (ROI C) on one Xenium slide 6 weeks post-implantation. Quality control analysis indicates counts per gene, transcript quality, mean Q-score, cellular clusters, counts per cell assigned to spatial coordinates, and graph-based cluster UMAPs organized by clusters and transcript counts.

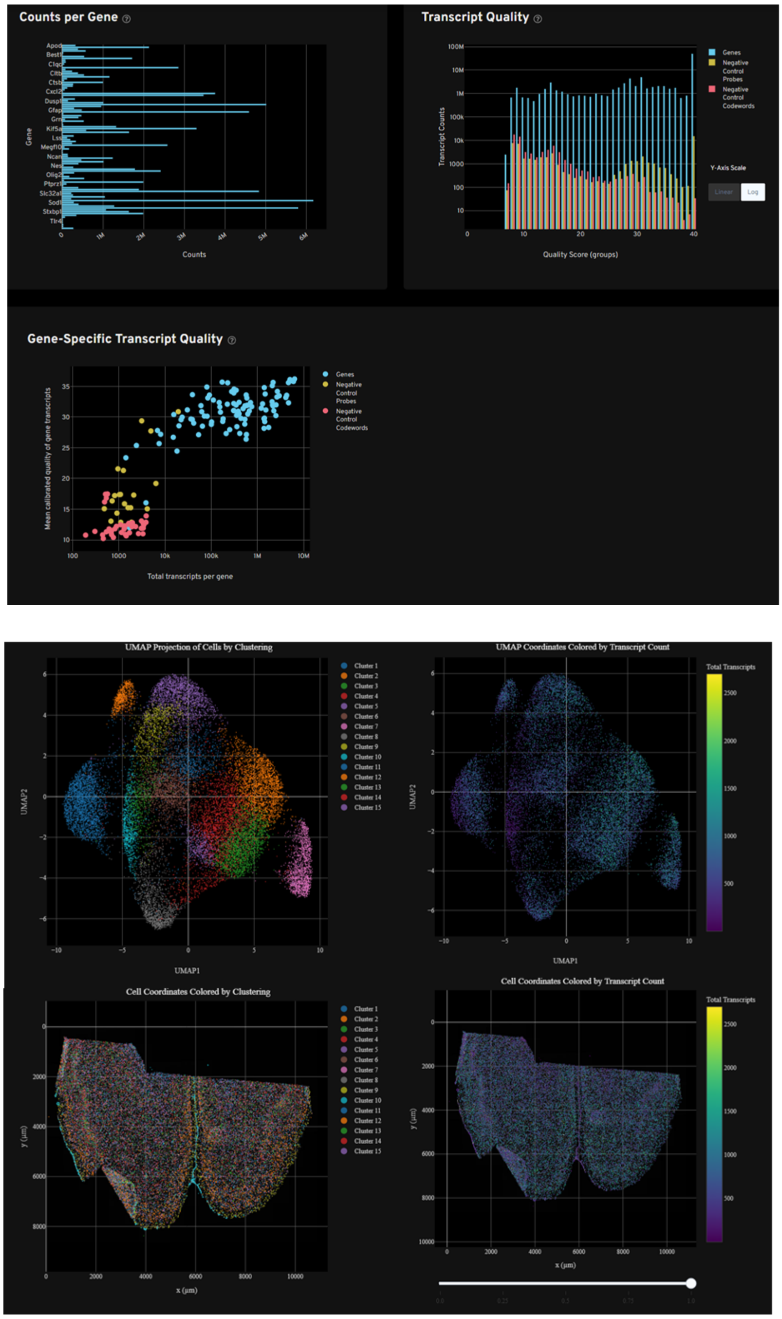


**Supplementary Figure 18:** Quality control analysis from the first bilateral polyimide sample (ROI A) on one Xenium slide 6 weeks post-implantation. Quality control analysis indicates counts per gene, transcript quality, cellular clusters and counts per cell assigned to spatial coordinates and graph-based cluster UMAPs organized by clusters and transcript counts.

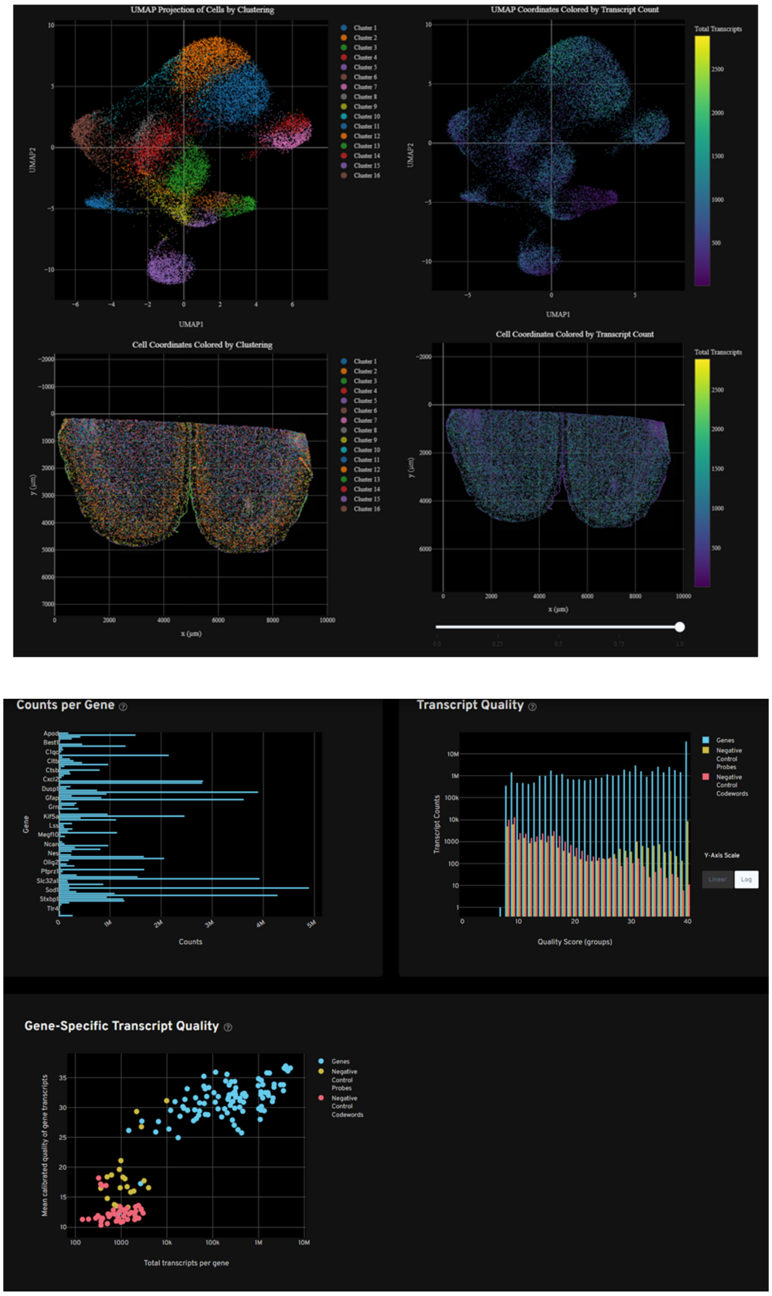


**Supplementary Figure 19:** Quality control analysis from the second bilateral polyimide sample (ROI B) on one Xenium slide 6-weeks post-implantation. Quality control analysis indicates counts per gene, transcript quality, cellular clusters and counts per cell assigned to spatial coordinates and graph-based cluster UMAPs organized by clusters and transcript counts.

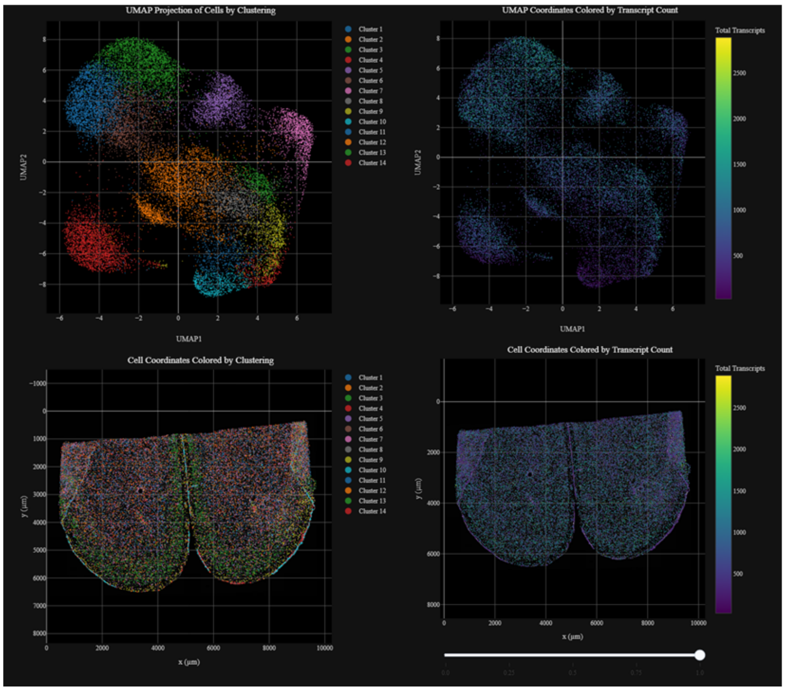


**Supplementary Figure 20:** Quality control analysis from the third bilateral polyimide sample (ROI C) on one Xenium slide 6-weeks post-implantation. Quality control analysis indicates counts per gene, transcript quality, cellular clusters and counts per cell assigned to spatial coordinates and graph-based cluster UMAPs organized by clusters and transcript counts.

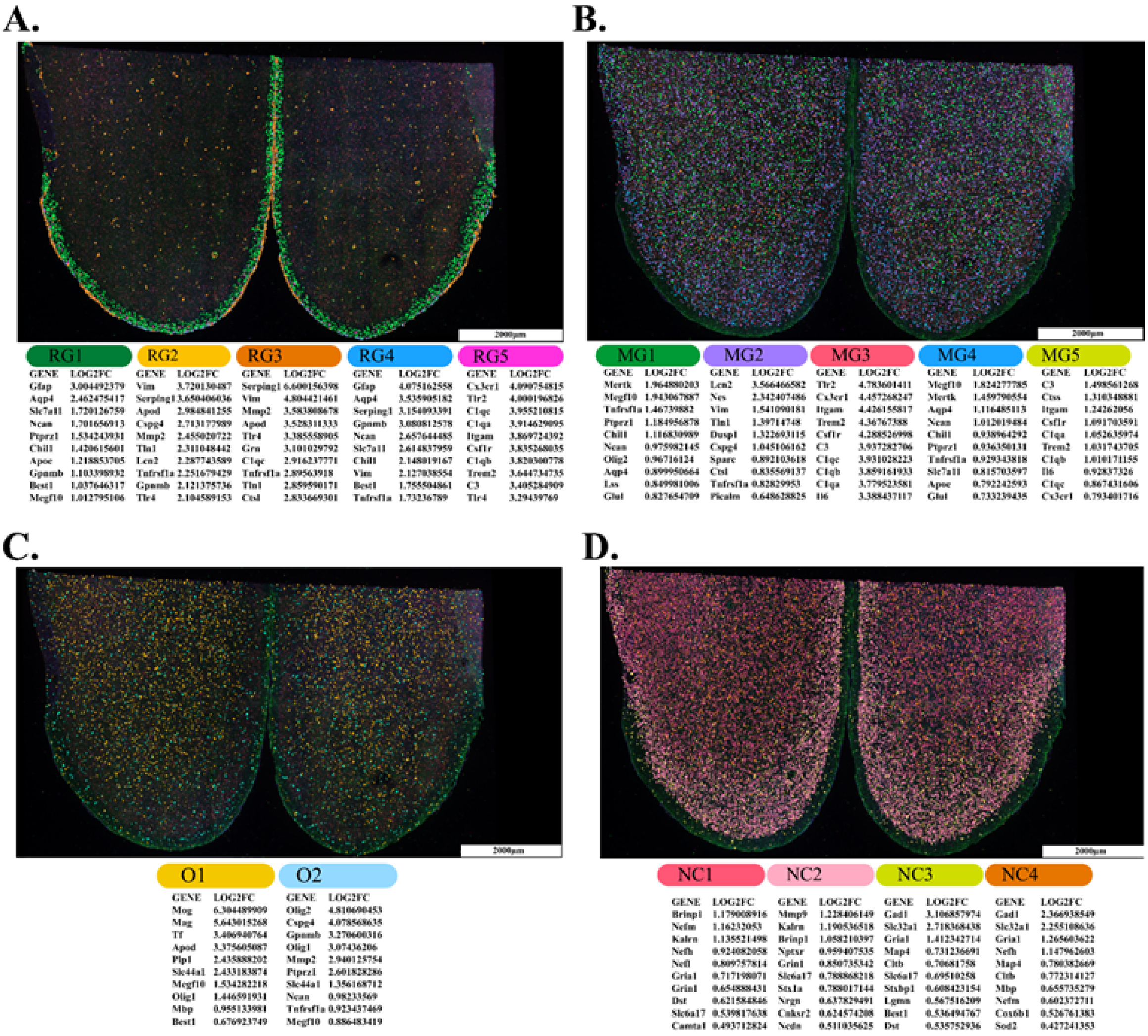


| RG1 GENE | RG1 LOG2FC | RG2 GENE | RG2 LOG2FC | RG3 GENE | RG3 LOG2FC | RG4 GENE | RG4 LOG2FC | RG5 GENE | RG5 LOG2FC |
|---|---|---|---|---|---|---|---|---|---|
| Gfap | 3.004492379 | Vim | 3.720130487 | Serping1 | 6.600156398 | Gfap | 4.075162558 | Cx3cr1 | 4.090754815 |
| Aqp4 | 2.462475417 | Serping1 | 3.650406036 | Vim | 4.804421461 | Aqp4 | 3.535905182 | Tlr2 | 4.000196826 |
| Slc7a11 | 1.720126759 | Apod | 2.984841255 | Mmp2 | 3.583808678 | Serping1 | 3.154093391 | C1qc | 3.955210815 |
| Ncan | 1.701656913 | Cspg4 | 2.713177989 | Apod | 3.528311333 | Gpnmb | 3.080812578 | C1qa | 3.914629095 |
| Ptprz1 | 1.534243931 | Mmp2 | 2.455020722 | Tlr4 | 3.385558905 | Ncan | 2.657644485 | Itgam | 3.869724392 |
| Chil1 | 1.420615601 | Tln1 | 2.311048442 | Grn | 3.101029792 | Slc7a11 | 2.614837959 | Csf1r | 3.835268035 |
| Apoe | 1.218853705 | Lcn2 | 2.287743589 | C1qc | 2.916237771 | Chil1 | 2.148019167 | C1qb | 3.820300778 |
| Gpnmb | 1.103398932 | Tnfrsf1a | 2.251679429 | Tnfrsf1a | 2.89563918 | Vim | 2.127038554 | Trem2 | 3.644734735 |
| Best1 | 1.037646317 | Gpnmb | 2.121375736 | Tln1 | 2.859590171 | Best1 | 1.755504861 | C3 | 3.405284909 |
| Megf10 | 1.012795106 | Tlr4 | 2.104589153 | Ctsl | 2.833669301 | Tnfrsf1a | 1.73236789 | Tlr4 | 3.29439769 |

| MG1 GENE | MG1 LOG2FC | MG2 GENE | MG2 LOG2FC | MG3 GENE | MG3 LOG2FC | MG4 GENE | MG4 LOG2FC | MG5 GENE | MG5 LOG2FC |
|---|---|---|---|---|---|---|---|---|---|
| Mertk | 1.964880203 | Lcn2 | 3.566466582 | Tlr2 | 4.783601411 | Megf10 | 1.824277785 | C3 | 1.498561268 |
| Megf10 | 1.943067887 | Nes | 2.342407486 | Cx3cr1 | 4.457268247 | Mertk | 1.459790554 | Ctss | 1.310348881 |
| Tnfrsf1a | 1.46739882 | Vim | 1.541090181 | Itgam | 4.426155817 | Aqp4 | 1.116485113 | Itgam | 1.24262056 |
| Ptprz1 | 1.184956878 | Tln1 | 1.39714748 | Trem2 | 4.36767388 | Ncan | 1.012019484 | Csf1r | 1.091703591 |
| Chil1 | 1.116830989 | Dusp1 | 1.322693115 | Csf1r | 4.288526998 | Chil1 | 0.938964292 | C1qa | 1.052635974 |
| Ncan | 0.975982145 | Cspg4 | 1.045106162 | C3 | 3.937282706 | Ptprz1 | 0.936350131 | Trem2 | 1.031743705 |
| Olig2 | 0.96716124 | Sparc | 0.892103618 | C1qc | 3.931028223 | Tnfrsf1a | 0.929343818 | C1qb | 1.010171155 |
| Aqp4 | 0.899950664 | Ctsl | 0.835569137 | C1qb | 3.859161933 | Slc7a11 | 0.815703597 | Il6 | 0.92837326 |
| Lss | 0.849981006 | Tnfrsf1a | 0.82829953 | C1qa | 3.779523581 | Apoe | 0.792242593 | C1qc | 0.867431606 |
| Glul | 0.827654709 | Picalm | 0.648628825 | Il6 | 3.388437117 | Glul | 0.733239435 | Cx3cr1 | 0.793401716 |

| O1 GENE | O1 LOG2FC | O2 GENE | O2 LOG2FC |
|---|---|---|---|
| Mog | 6.304489909 | Olig2 | 4.810690453 |
| Mag | 5.643015268 | Cspg4 | 4.078568635 |
| Tf | 3.406940764 | Gpnmb | 3.270600316 |
| Apod | 3.375605087 | Olig1 | 3.07436206 |
| Plp1 | 2.435888202 | Mmp2 | 2.940125754 |
| Slc44a1 | 2.433183874 | Ptprz1 | 2.601828286 |
| Megf10 | 1.534282218 | Slc44a1 | 1.356168712 |
| Olig1 | 1.446591931 | Ncan | 0.98233569 |
| Mbp | 0.955133981 | Tnfrsf1a | 0.923437469 |
| Best1 | 0.676923749 | Megf10 | 0.886483419 |

| NC1 GENE | NC1 LOG2FC | NC2 GENE | NC2 LOG2FC | NC3 GENE | NC3 LOG2FC | NC4 GENE | NC4 LOG2FC |
|---|---|---|---|---|---|---|---|
| Brinp1 | 1.179008916 | Mmp9 | 1.228406149 | Gad1 | 3.106857974 | Gad1 | 2.366938549 |
| Nefm | 1.16232053 | Kalrn | 1.190536518 | Slc32a1 | 2.718368438 | Slc32a1 | 2.255108636 |
| Kalrn | 1.135521498 | Brinp1 | 1.058210397 | Gria1 | 1.412342714 | Gria1 | 1.265603622 |
| Nefh | 0.924082058 | Nptxr | 0.959407535 | Map4 | 0.731236691 | Nefh | 1.147962603 |
| Nefl | 0.809757814 | Grin1 | 0.850735342 | Cltb | 0.70681758 | Map4 | 0.780382669 |
| Gria1 | 0.717198071 | Slc6a17 | 0.788868218 | Slc6a17 | 0.69510258 | Cltb | 0.772314127 |
| Grin1 | 0.654888431 | Stx1a | 0.788017144 | Stxbp1 | 0.608423154 | Mbp | 0.655735279 |
| Dst | 0.621584846 | Nrgn | 0.637829491 | Lgmn | 0.567516209 | Nefm | 0.602372711 |
| Slc6a17 | 0.539817638 | Cnksr2 | 0.624574208 | Best1 | 0.536494767 | Cox6b1 | 0.526761383 |
| Camta1 | 0.493712824 | Ncdn | 0.511035625 | Dst | 0.535752936 | Sod2 | 0.427241353 |

**Supplementary Figure 21**: Representative confocal images of whole section immunofluorescence stains registered to Xenium-generated cellular clusters of a naïve sample paired with the top 10 differentially expressed genes within each cluster for (A) reactive glia, (B) modulatory glia, (C) oligodendrocytes, and (D) neurons.

**A.**

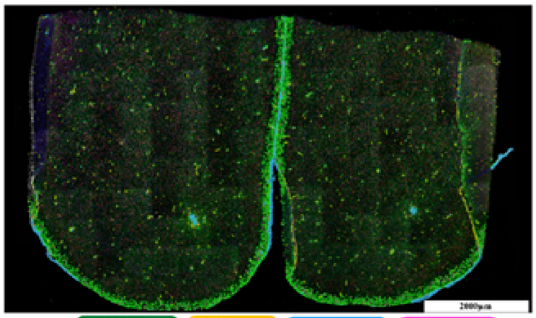


| RG1 GENE | RG1 LOG2FC | RG2 GENE | RG2 LOG2FC | RG4 GENE | RG4 LOG2FC | RG5 GENE | RG5 LOG2FC |
|---|---|---|---|---|---|---|---|
| Gfap | 3.298617162 | Vim | 3.683053885 | Serping1 | 5.505311014 | Cx3cr1 | 3.69294639 |
| Aqp4 | 2.858451147 | Serping1 | 3.674567943 | Vim | 4.9045925 | C1qa | 3.688198099 |
| Best1 | 2.060279151 | Lcn2 | 3.057513765 | Gpnmb | 3.303761406 | C1qc | 3.676509486 |
| Slc7a11 | 1.940810241 | Apod | 2.429264453 | Tlr4 | 3.255590192 | C1qb | 3.59918237 |
| Ncan | 1.792570487 | Tlr4 | 2.370818905 | Apod | 3.245394857 | Tlr2 | 3.564229848 |
| Gpnmb | 1.78302157 | Cspg4 | 2.180501638 | Grn | 3.137788015 | Itgam | 3.508415775 |
| Chil1 | 1.698740486 | Tln1 | 2.100257172 | Tln1 | 3.070330692 | Csf1r | 3.456892977 |
| Serping1 | 1.672591117 | Tnfrsf1a | 2.064644695 | Tnfrsf1a | 3.032406404 | C3 | 3.441183924 |
| Ptprz1 | 1.446363908 | Gpnmb | 2.036933361 | Mmp2 | 2.95370493 | Trem2 | 3.160220477 |
| Tnf | 1.347889813 | Mmp2 | 1.936217624 | Slc7a11 | 2.924290263 | Ctss | 3.017415604 |

**B.**

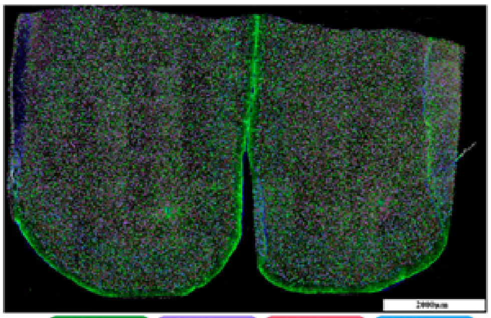


| MG1 GENE | MG1 LOG2FC | MG2 GENE | MG2 LOG2FC | MG3 GENE | MG3 LOG2FC | MG4 GENE | MG4 LOG2FC |
|---|---|---|---|---|---|---|---|
| Megf10 | 1.816827274 | Lcn2 | 2.126552844 | Tlr2 | 4.368013486 | Lcn2 | 1.284411293 |
| Mertk | 1.457080084 | Nes | 1.568031431 | Cx3cr1 | 4.224270851 | Tnf | 1.190060589 |
| Aqp4 | 1.372251177 | Tln1 | 1.102648088 | Trem2 | 4.140246081 | Nes | 1.133627829 |
| Ncan | 1.189097344 | Cspg4 | 0.964341567 | Itgam | 4.055850867 | Cxcl2 | 1.007801039 |
| Chil1 | 1.101970488 | Mbp | 0.865058092 | Csf1r | 3.936703302 | Camk2a | 0.793779362 |
| Ptprz1 | 1.094588032 | Ctsl | 0.735737547 | C3 | 3.78791884 | Best1 | 0.749115217 |
| Tnfrsf1a | 0.980192715 | Vim | 0.706251924 | C1qc | 3.558956121 | Sparc | 0.692820283 |
| Apoe | 0.963751299 | Tnfrsf1a | 0.683201632 | C1qb | 3.485521211 | Shank2 | 0.690022579 |
| Slc7a11 | 0.926827621 | Sparc | 0.673352112 | C1qa | 3.456193706 | Tln1 | 0.550253755 |
| Glul | 0.855927612 | Nefm | 0.627098719 | Ctss | 2.982626819 | Slc7a11 | 0.526588615 |

**C.**

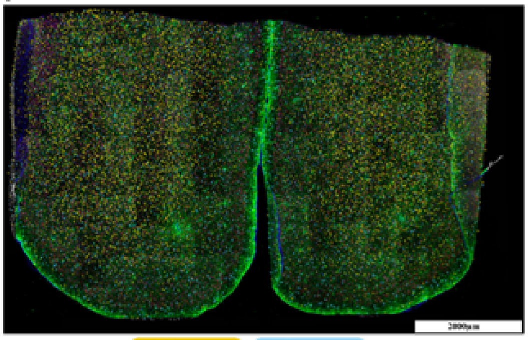


| O1 GENE | O1 LOG2FC | O2 GENE | O2 LOG2FC |
|---|---|---|---|
| Mog | 5.835550402 | Olig2 | 4.862618137 |
| Mag | 5.020375098 | Cspg4 | 3.878586085 |
| Apod | 3.316127832 | Gpnmb | 3.01056204 |
| Tf | 2.931632568 | Olig1 | 2.899889686 |
| Slc44a1 | 2.408703052 | Mmp2 | 2.877443399 |
| Plp1 | 2.019763926 | Ptprz1 | 2.552241708 |
| Megf10 | 1.720209436 | Slc44a1 | 1.075572928 |
| Olig1 | 1.415832881 | Tnfrsf1a | 1.04613136 |
| Best1 | 1.223835937 | Megf10 | 0.910584477 |
| Mbp | 0.928987323 | Ncan | 0.856023454 |

**D.**

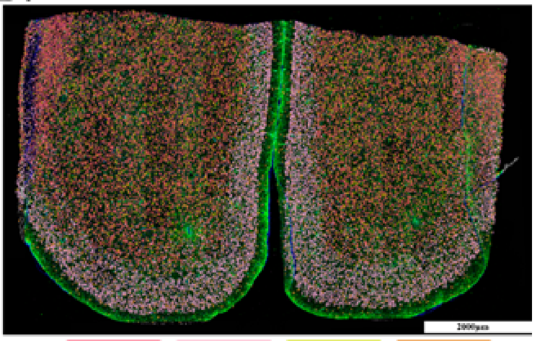


| NC1 GENE | NC1 LOG2FC | NC2 GENE | NC2 LOG2FC | NC3 GENE | NC3 LOG2FC | NC4 GENE | NC4 LOG2FC |
|---|---|---|---|---|---|---|---|
| Brinp1 | 1.356641925 | Kalrn | 1.037146617 | Slc32a1 | 3.165286957 | Gad1 | 3.030344248 |
| Kalrn | 1.337575733 | Brinp1 | 1.016661199 | Gad1 | 3.056873742 | Slc32a1 | 2.800347579 |
| Nefm | 1.028192327 | Nptxr | 1.00882127 | Gria1 | 1.724714562 | Gria1 | 1.183290861 |
| Dusp1 | 0.926553755 | Gria1 | 0.978255533 | Nefh | 1.339331882 | Best1 | 0.889031317 |
| Gria1 | 0.826440423 | Stx1a | 0.935586237 | Map4 | 1.047211783 | Cltb | 0.791677868 |
| Slc6a17 | 0.76858557 | Mmp9 | 0.88732838 | Cltb | 0.93690505 | Stxbp1 | 0.671193354 |
| Nefh | 0.767514671 | Slc6a17 | 0.684822306 | Slc6a17 | 0.803771659 | Map4 | 0.636221183 |
| Dst | 0.756546399 | Cnksr2 | 0.677845951 | Sod2 | 0.63826824 | Sod1 | 0.588179703 |
| Grin1 | 0.732017773 | Nrgn | 0.641482537 | Cox6b1 | 0.62240972 | Slc6a17 | 0.568925446 |
| Nefl | 0.669910958 | Nedn | 0.611087871 | Nefm | 0.605651111 | Cxcl2 | 0.505808451 |

**Supplementary Figure 22:** Representative confocal images of whole-section immunofluorescence stains registered to Xenium-generated cellular clusters of a bilateral polyimide sample 1-week post-implantation paired with the top 10 differentially expressed genes within each cluster for (A) reactive glia, (B) modulatory glia, (C) oligodendrocytes, and (D) neurons.

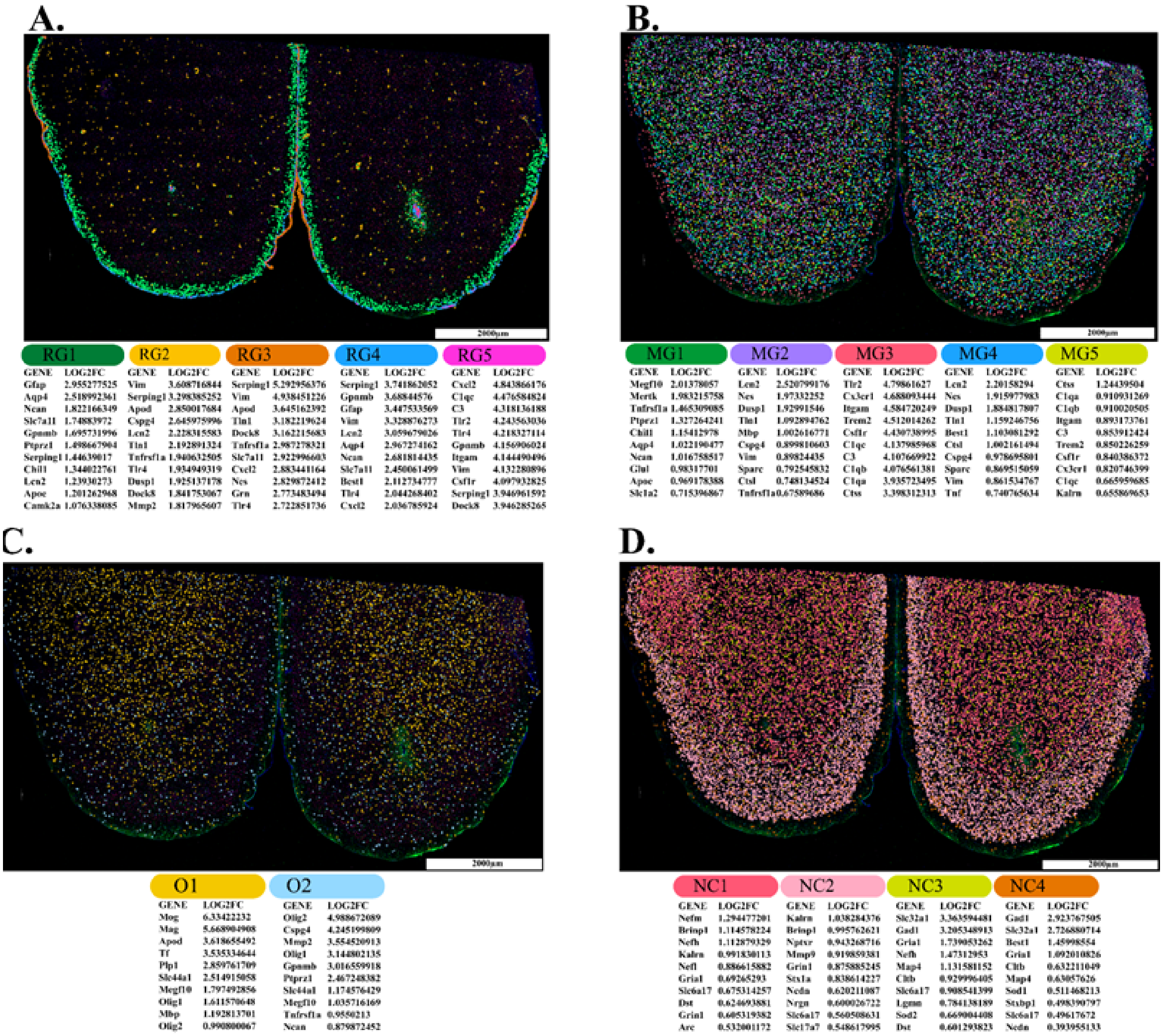


**Supplementary Figure 23:** Representative confocal images of whole section immunofluorescence stains registered to Xenium-generated cellular clusters of a bilateral polyimide sample 6 weeks post-implantation paired with the top 10 differentially expressed genes within each cluster for (A) reactive glia, (B) modulatory glia, (C) oligodendrocytes, and (D) neurons.

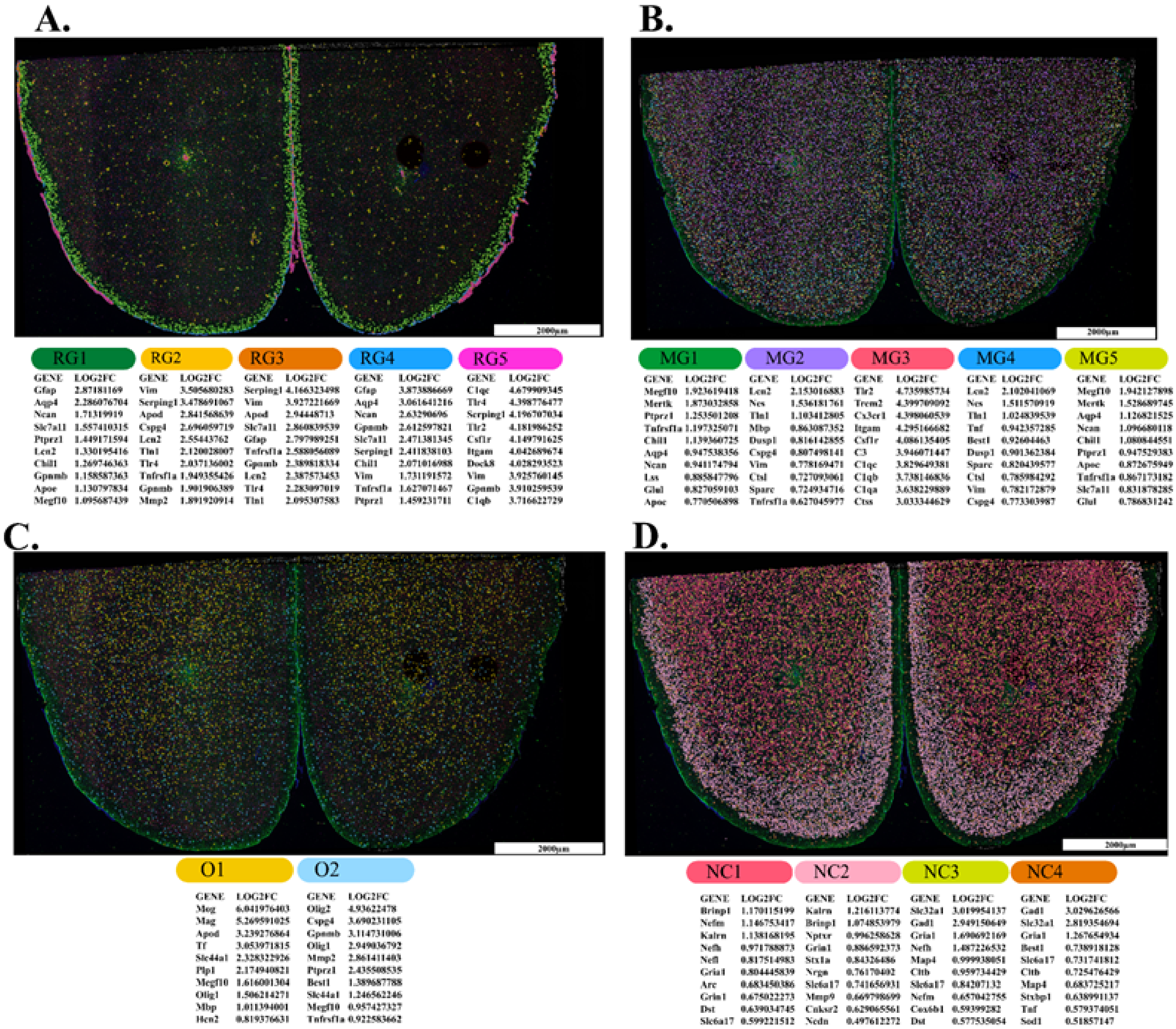


**Supplementary Figure 24:** Representative confocal images of whole section immunofluorescence stains registered to xenium-generated cellular clusters of a bilateral silicon sample 1-week post-implantation paired with the top 10 differentially expressed genes within each cluster for (A) reactive glia, (B) modulatory glia, (C) oligodendrocytes, and (D) neurons.

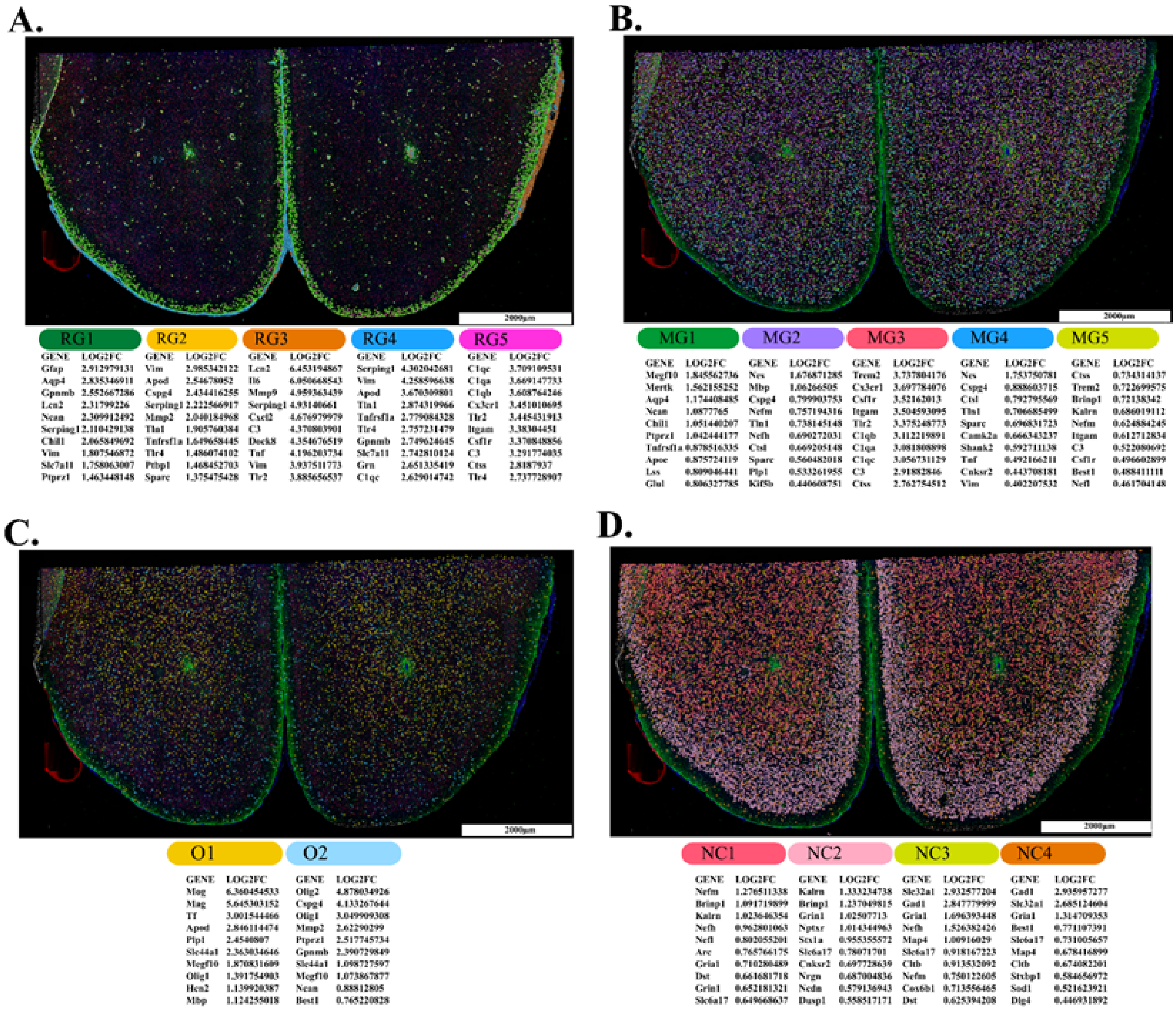


**Supplementary Figure 25:** Representative confocal images of whole-section immunofluorescence stains registered to Xenium-generated cellular clusters of a bilateral silicon sample 6 weeks post-implantation paired with the top 10 differentially expressed genes within each cluster for (A) reactive glia, (B) modulatory glia, (C) oligodendrocytes, and (D) neurons.

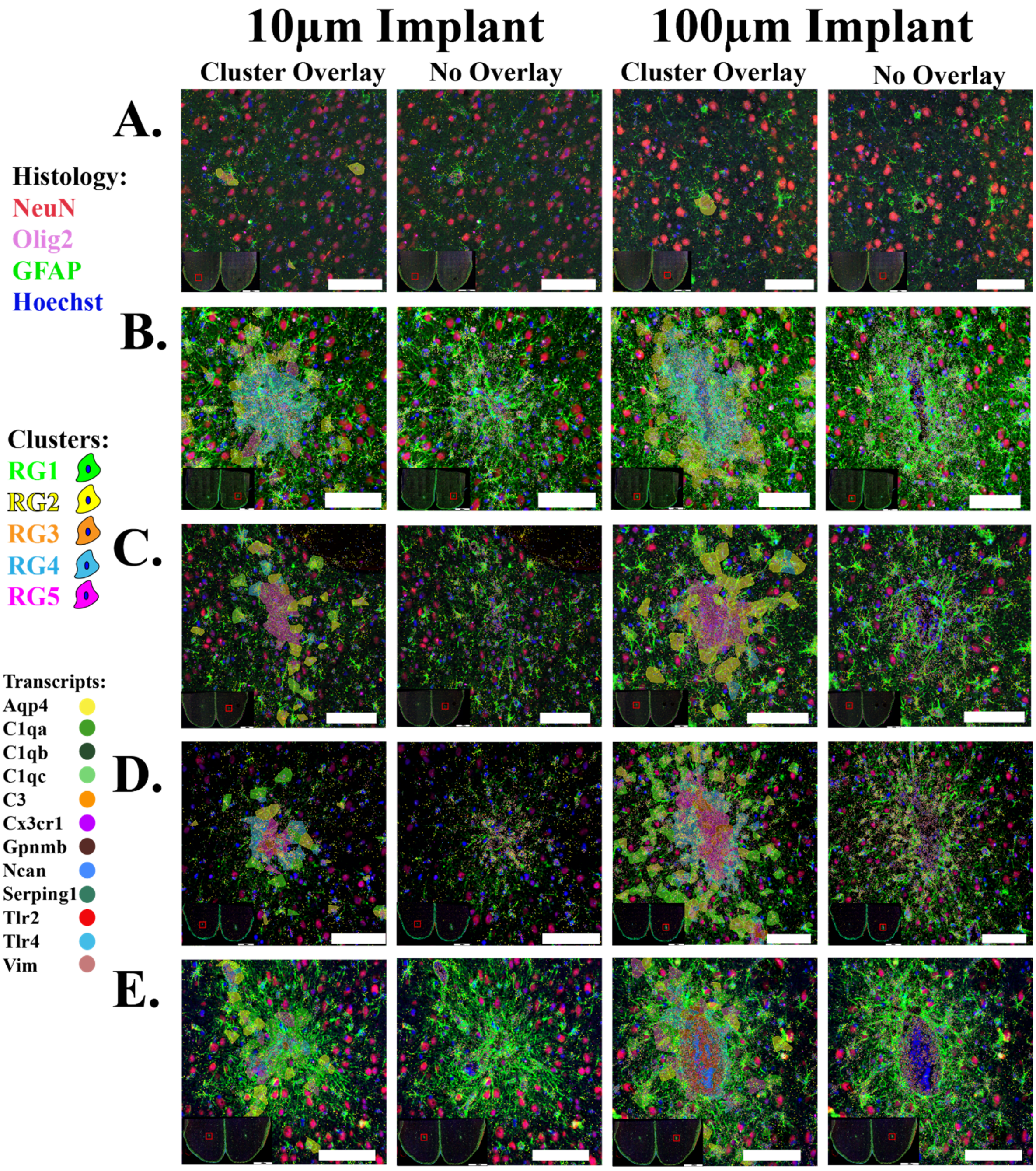


**Supplementary Figure 26:** Representative immunofluorescence images registered with Xenium data. Common reactive glia clusters (colored cell overlays) and top differentially expressed genes (transcripts identified as colored puncta) from (A) naive, (B) 1-week polyimide, (C) 1-week silicon, (D) 6-week polyimide, and (E) 6-week silicon-implanted tissue.

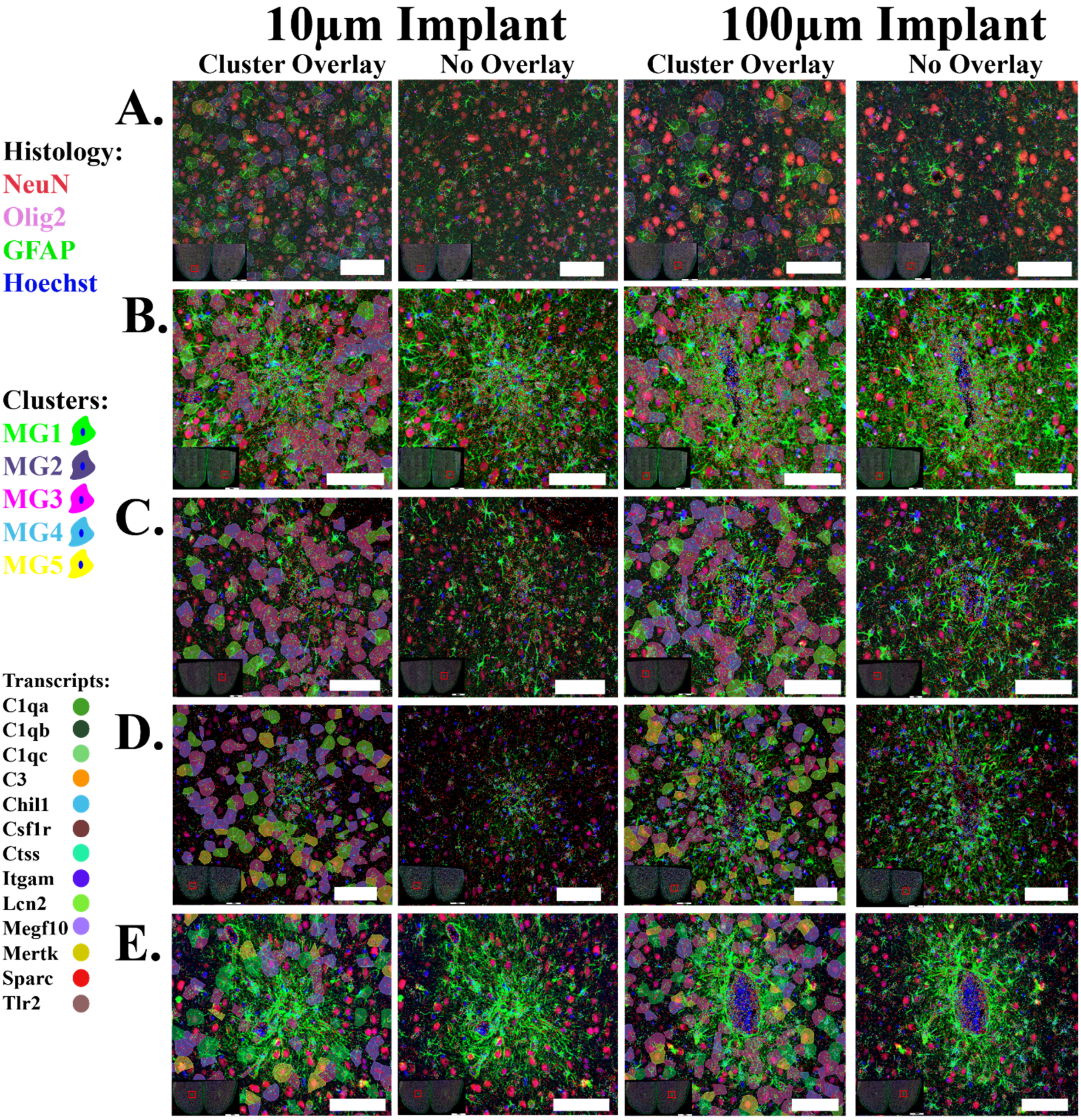


**Supplementary Figure 27:** Representative immunofluorescence images registered with Xenium data. Common modulatory glia clusters (colored cell overlays) and top differentially expressed genes (transcripts identified as colored puncta) from (A) naive, (B) 1-week polyimide, (C) 1-week silicon, (D) 6-week polyimide, and (E) 6-week silicon-implanted tissue.

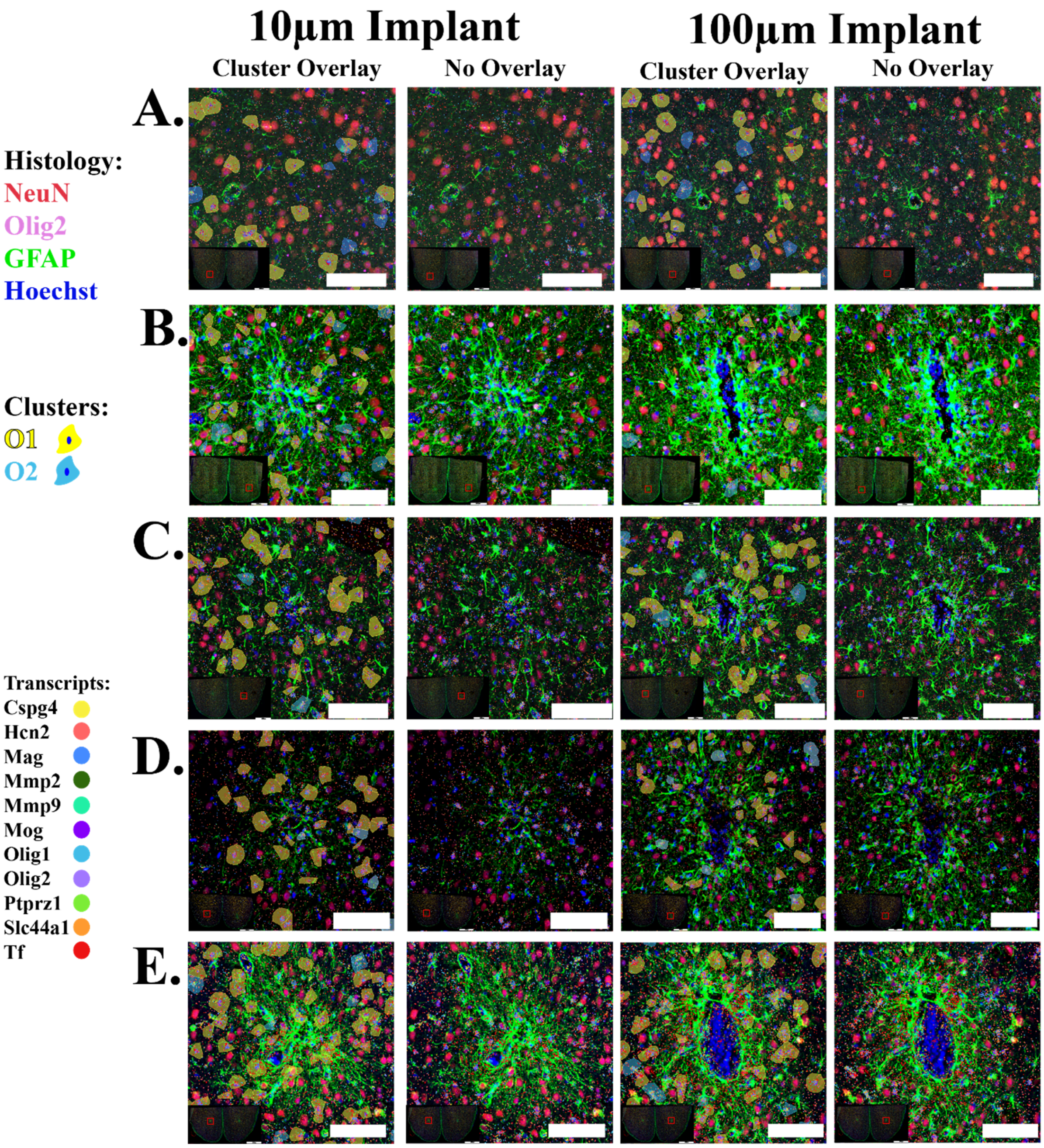


**Supplementary Figure 28**: Representative immunofluorescence images registered with Xenium data. Common oligodendrocyte clusters (colored cell overlays) and top differentially expressed genes (transcripts identified as colored puncta) from (A) naive, (B) 1-week polyimide, (C) 1-week silicon, (D) 6-week polyimide, and (E) 6-week silicon-implanted tissue.

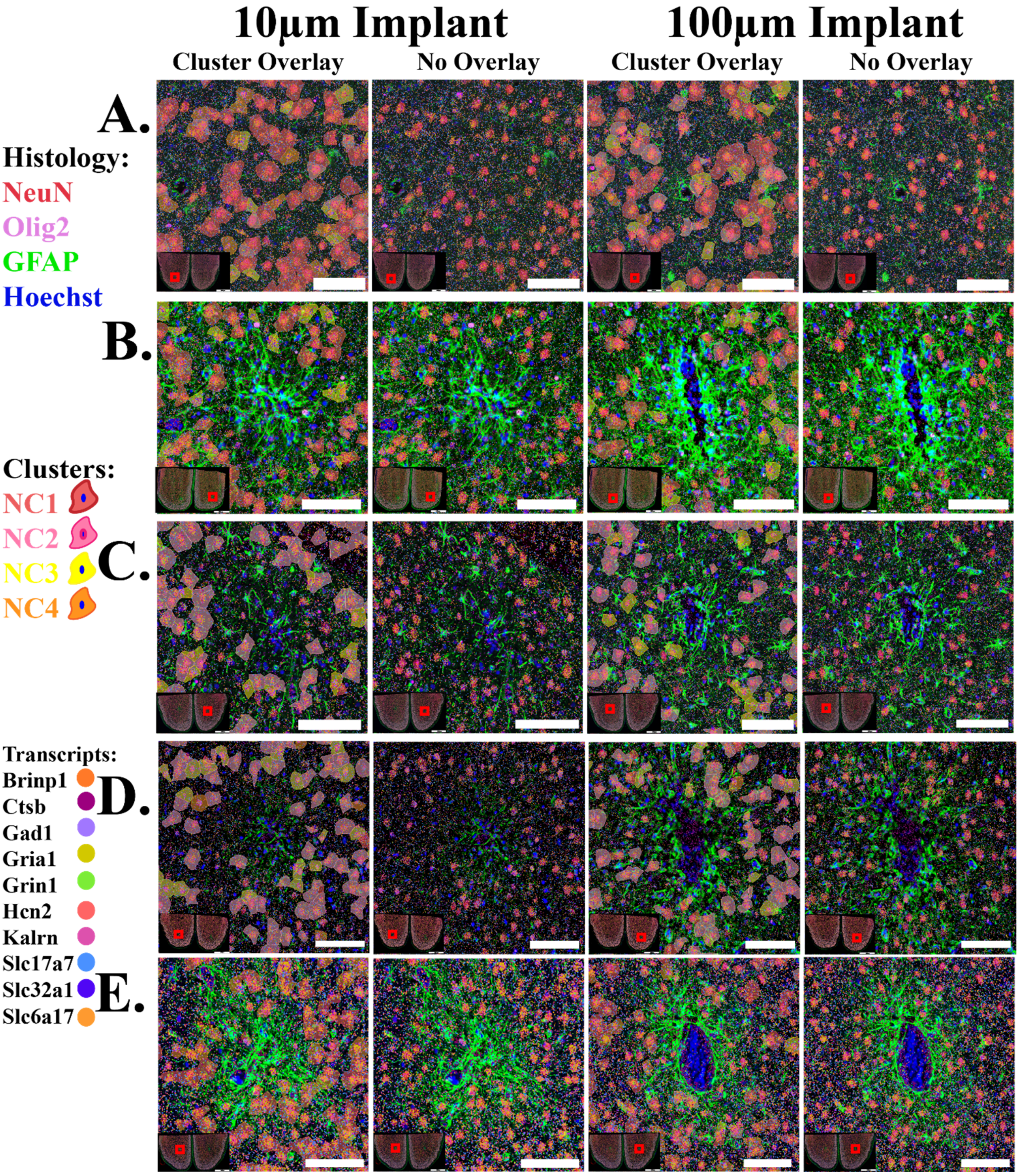


**Supplementary Figure 29:** Representative immunofluorescence images registered with Xenium data. Common neuronal clusters (colored cell overlays) and top differentially expressed genes (transcripts identified as colored puncta) from (A) naive, (B) 1-week polyimide, (C) 1-week silicon, (D) 6-week polyimide, and (E) 6-week silicon-implanted tissue.

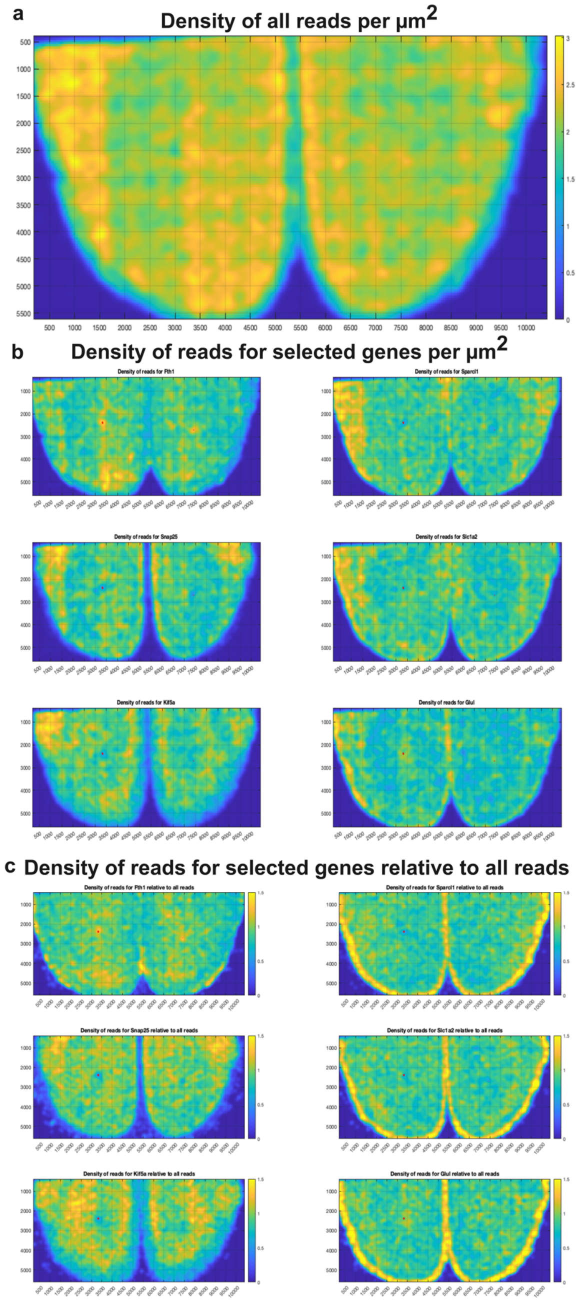


**Supplementary Figure 30:** (a) A spatial map of the overall read density (reads per μm$^2$) across the tissue section, showing significant variation in read density and a grid-like pattern consistent with an artifact from the stitching of individual fields of view during Xenium image reconstruction. (b) Spatial density maps of six highly abundant genes, which exhibit grid-like patterns similar to those observed in the overall read density. (c) Gene densities normalized by the total read density, revealing biological spatial patterns while largely removing the grid-like artifact. Implant locations are indicated by red dots.

**Supplementary Figure 31: Normalized gene density across distinct cell types in peri-implant and naïve tissues.** Box plots display gene expression density (reads/μm$^2$) for individual cell types across three regions: 100 μm from the device, 10 μm from the device, and naïve control tissue. Data groups include 1-week silicon (blue), 1-week polyimide (orange), 6-week polyimide (green), 6-week silicon (purple), and naïve tissue (grey). The horizontal line within each box represents the median; the diamond marker indicates the mean, and whiskers extend to the lowest and highest values within 1.5 × IQR of the first and third quartiles.

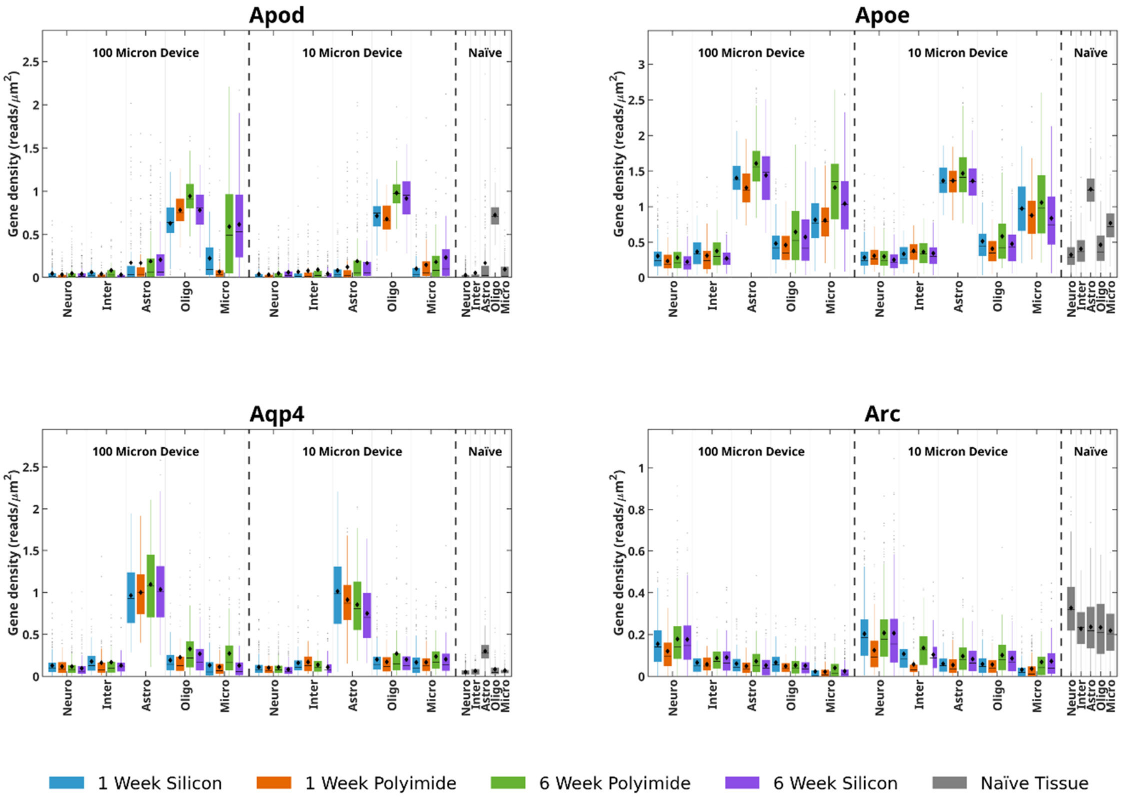

#### Supplementary Figure 31 (Contd.)

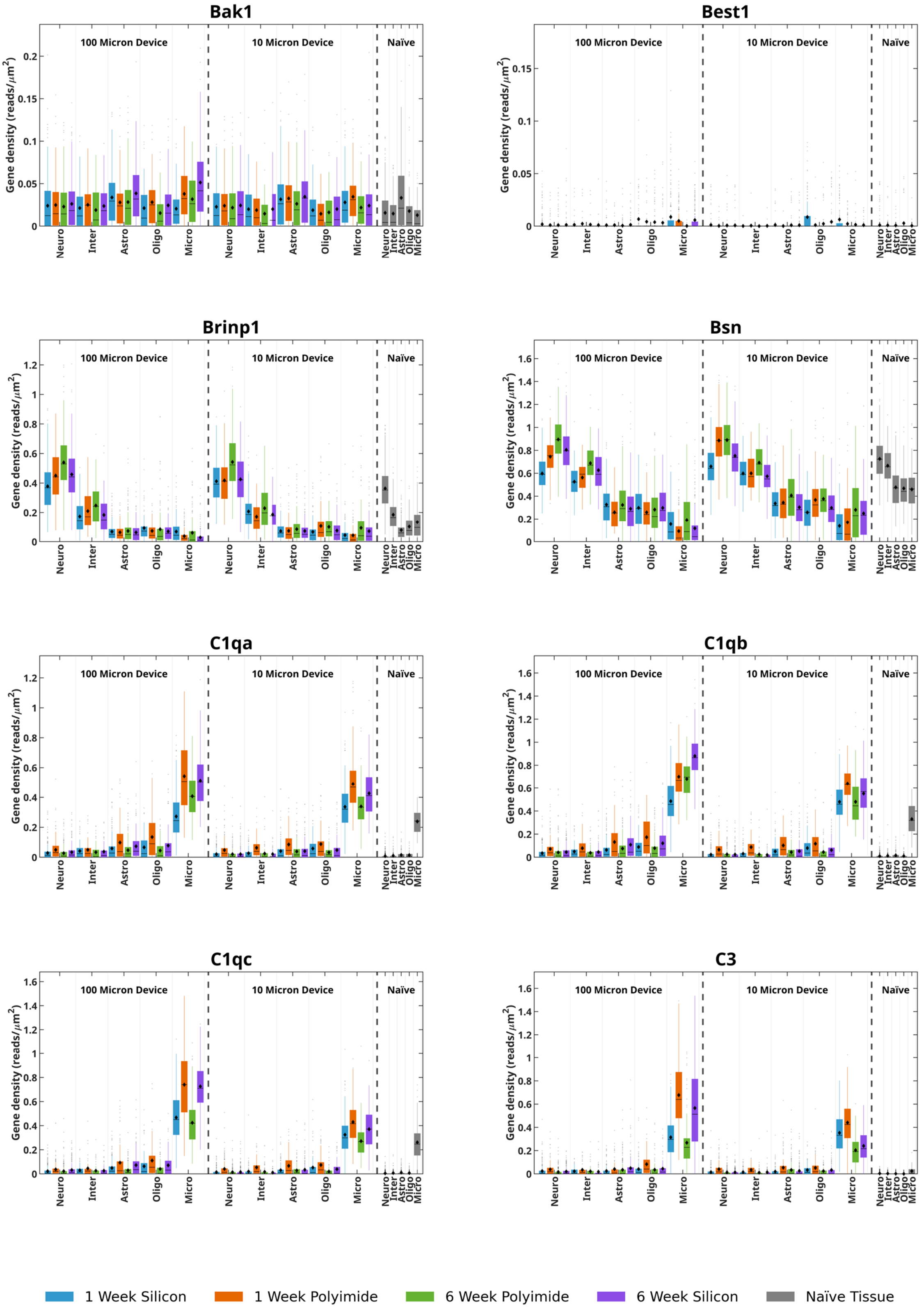

#### Supplementary Figure 31 (Contd.)

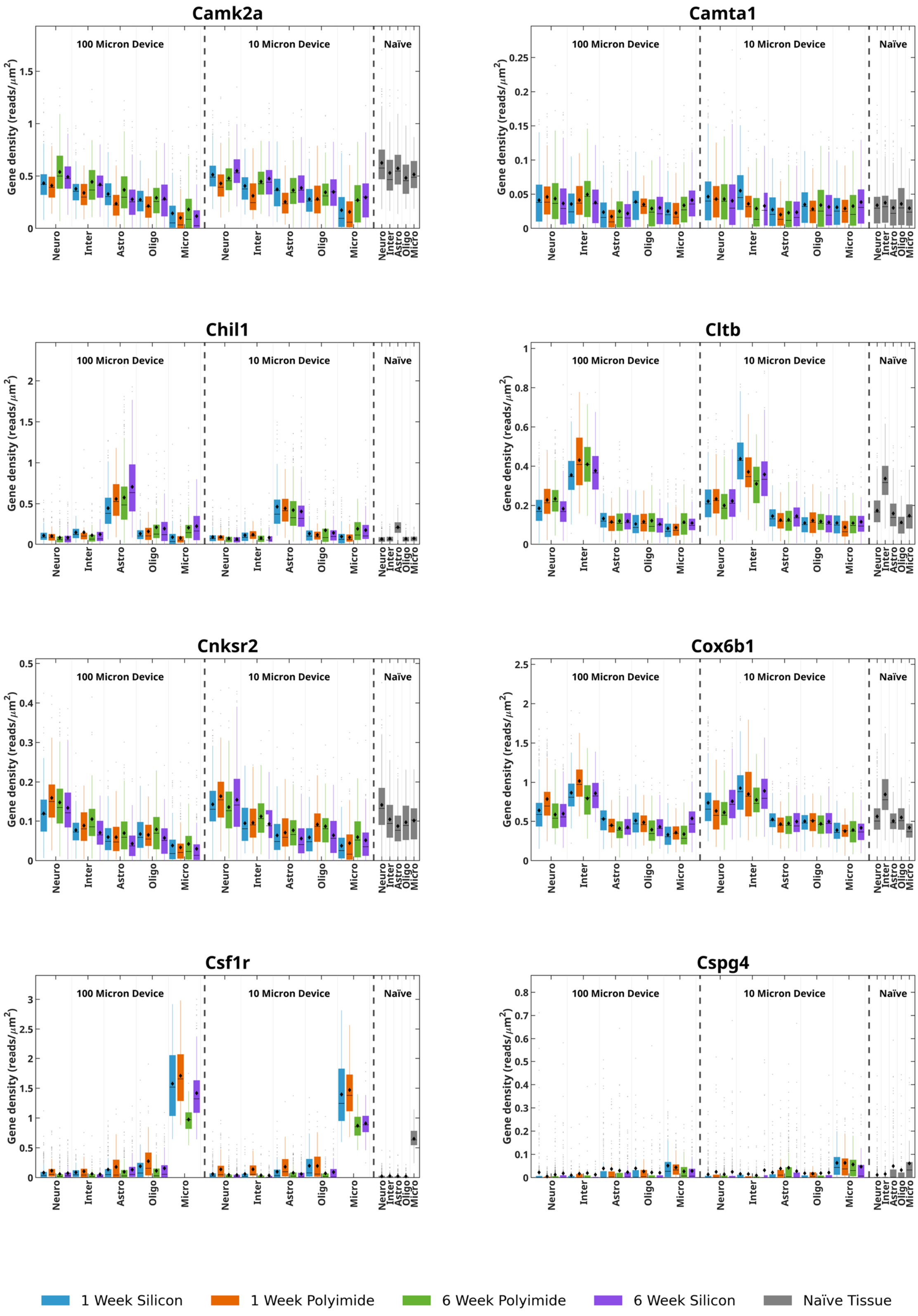

#### Supplementary Figure 31 (Contd.)

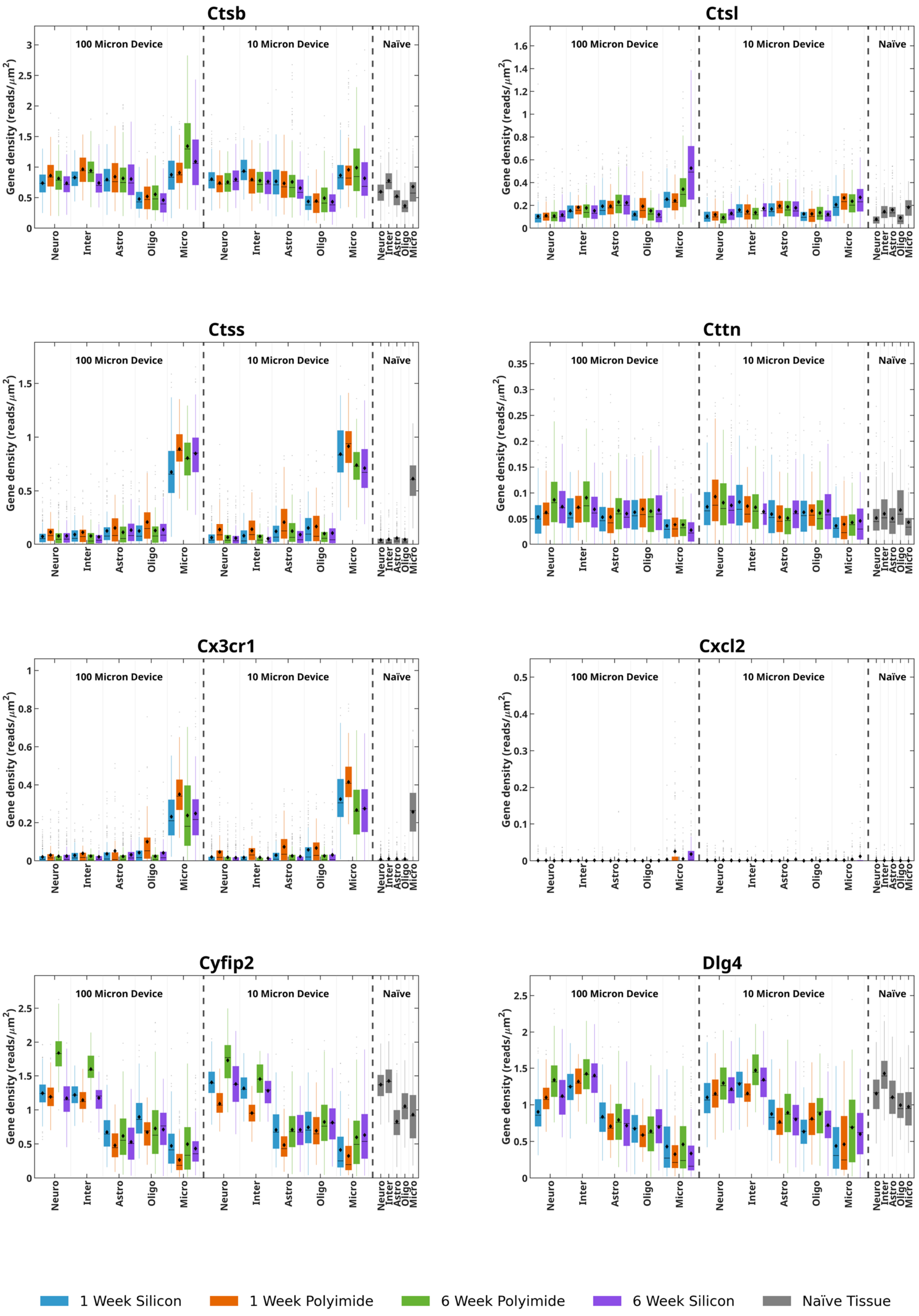

#### Supplementary Figure 31 (Contd.)

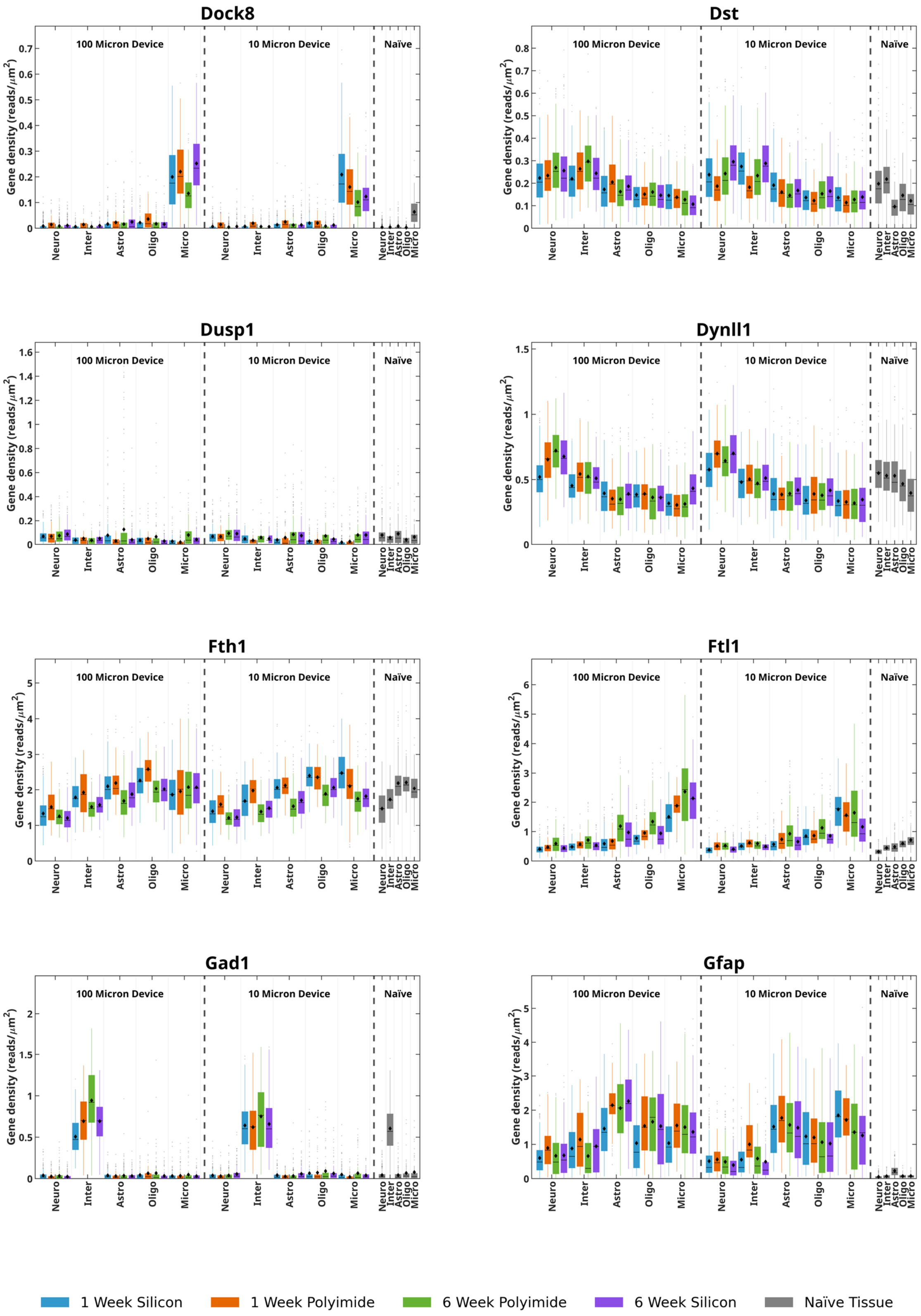

### Supplementary Figure 31 (Contd.)

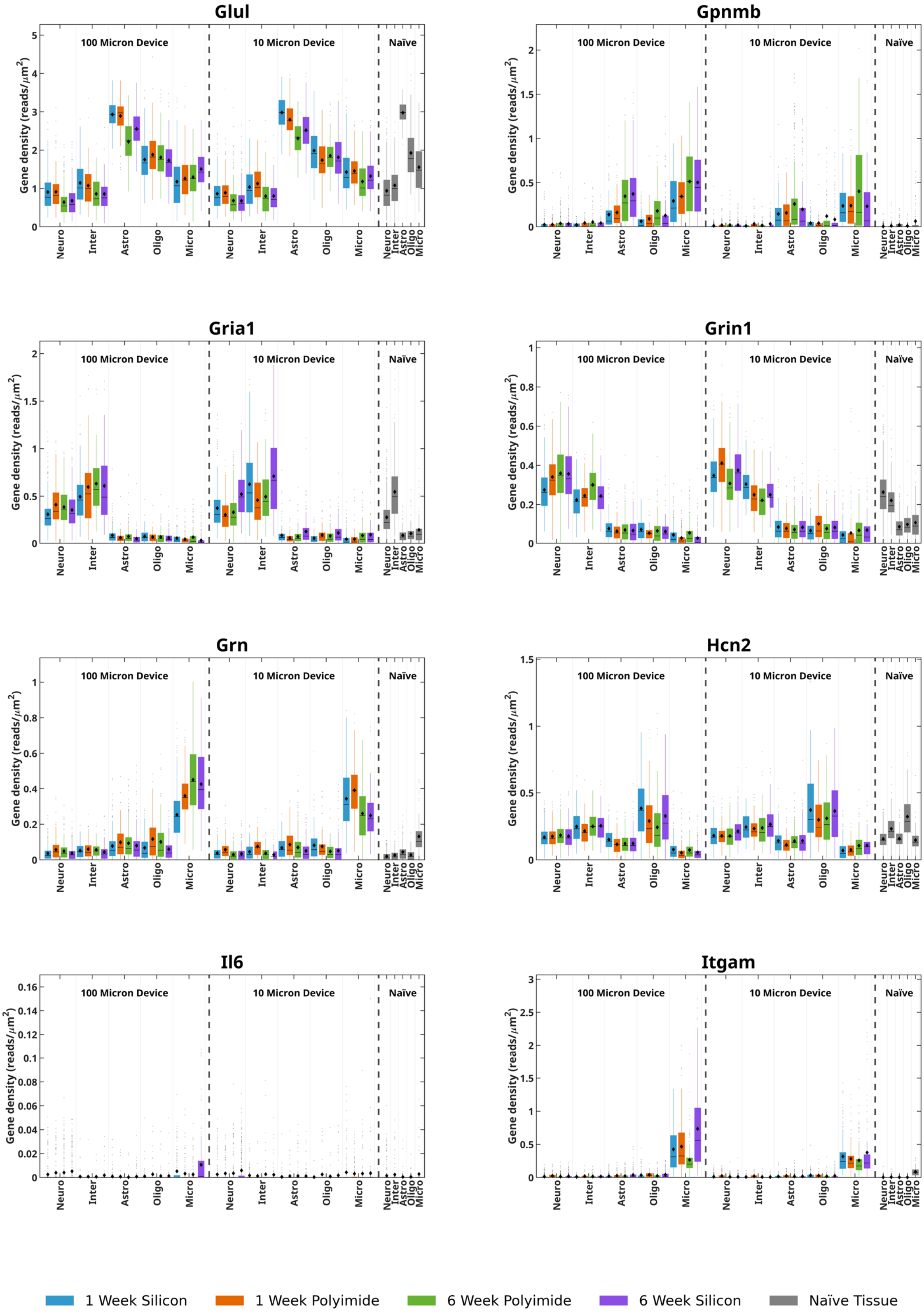

#### Supplementary Figure 31 (Contd.)

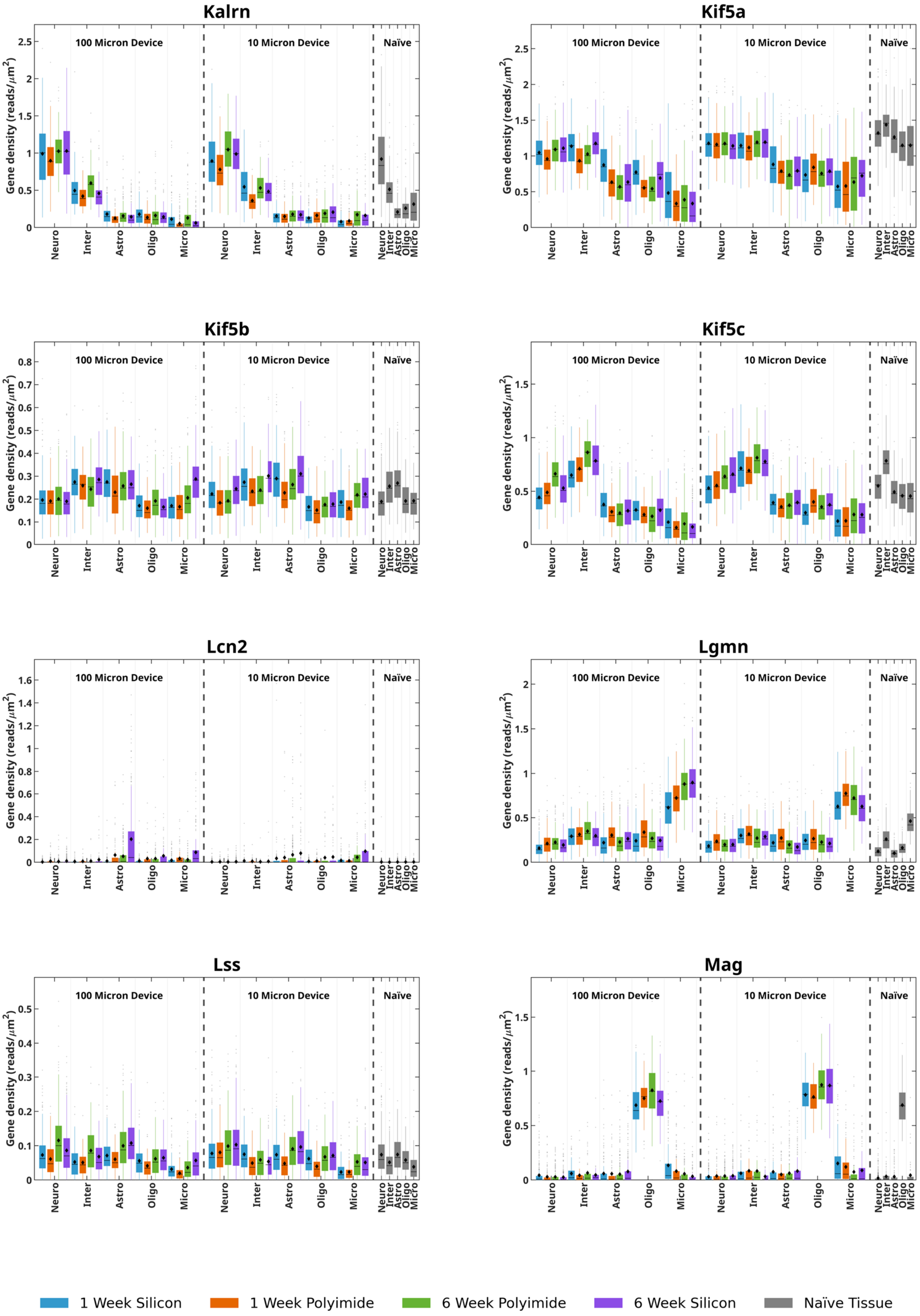

#### Supplementary Figure 31 (Contd.)

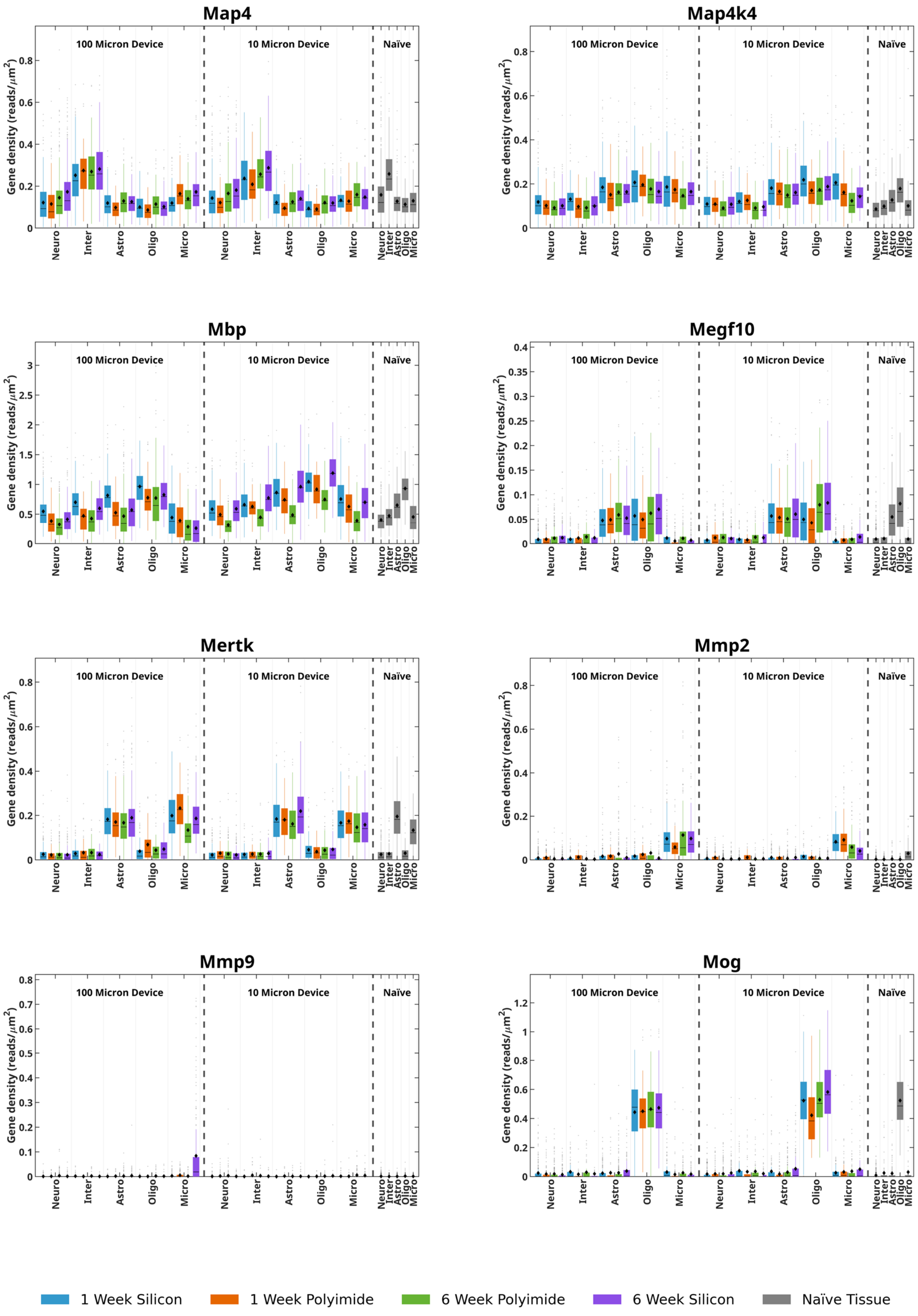

#### Supplementary Figure 31 (Contd.)

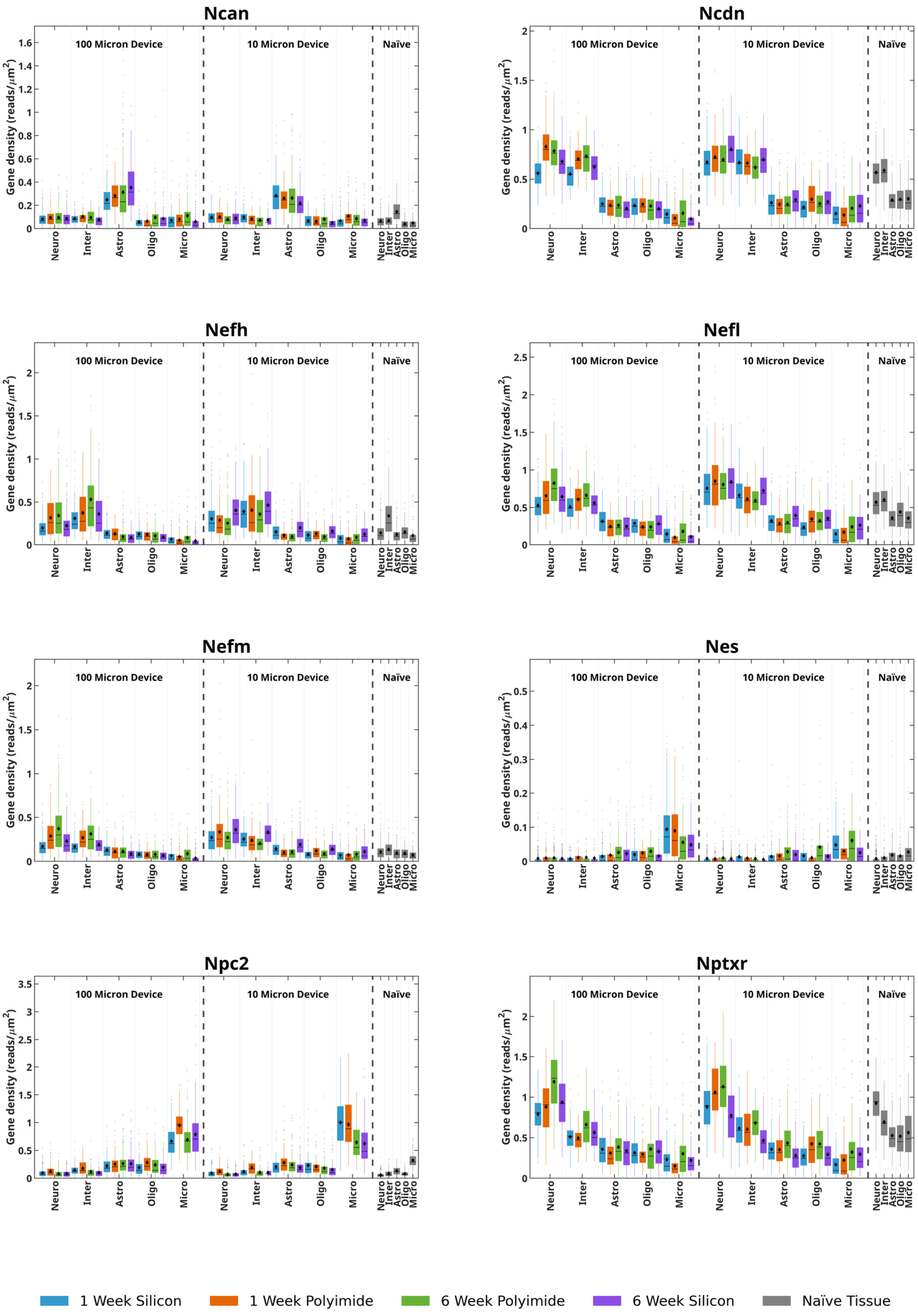

#### Supplementary Figure 31 (Contd.)

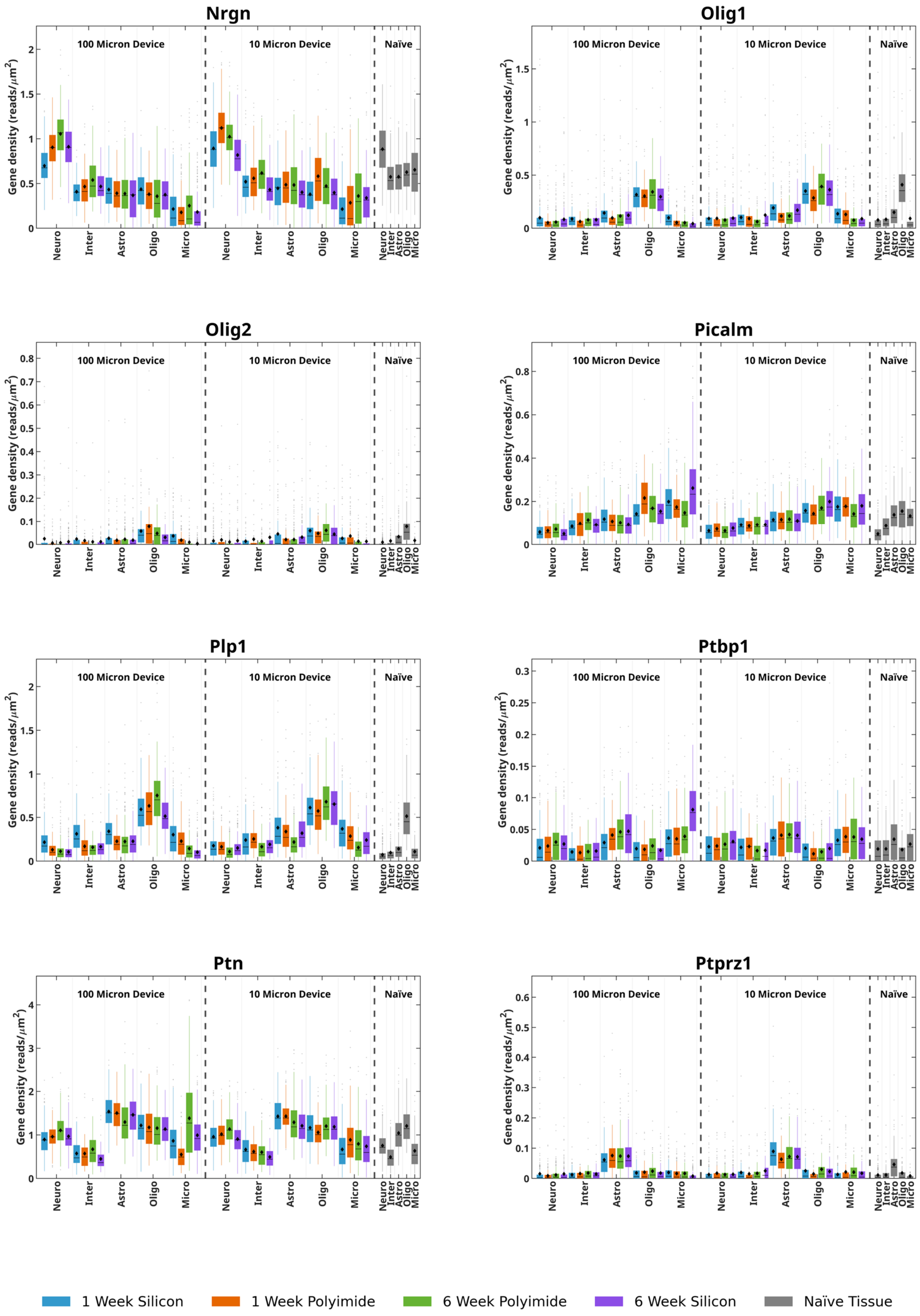

#### Supplementary Figure 31 (Contd.)

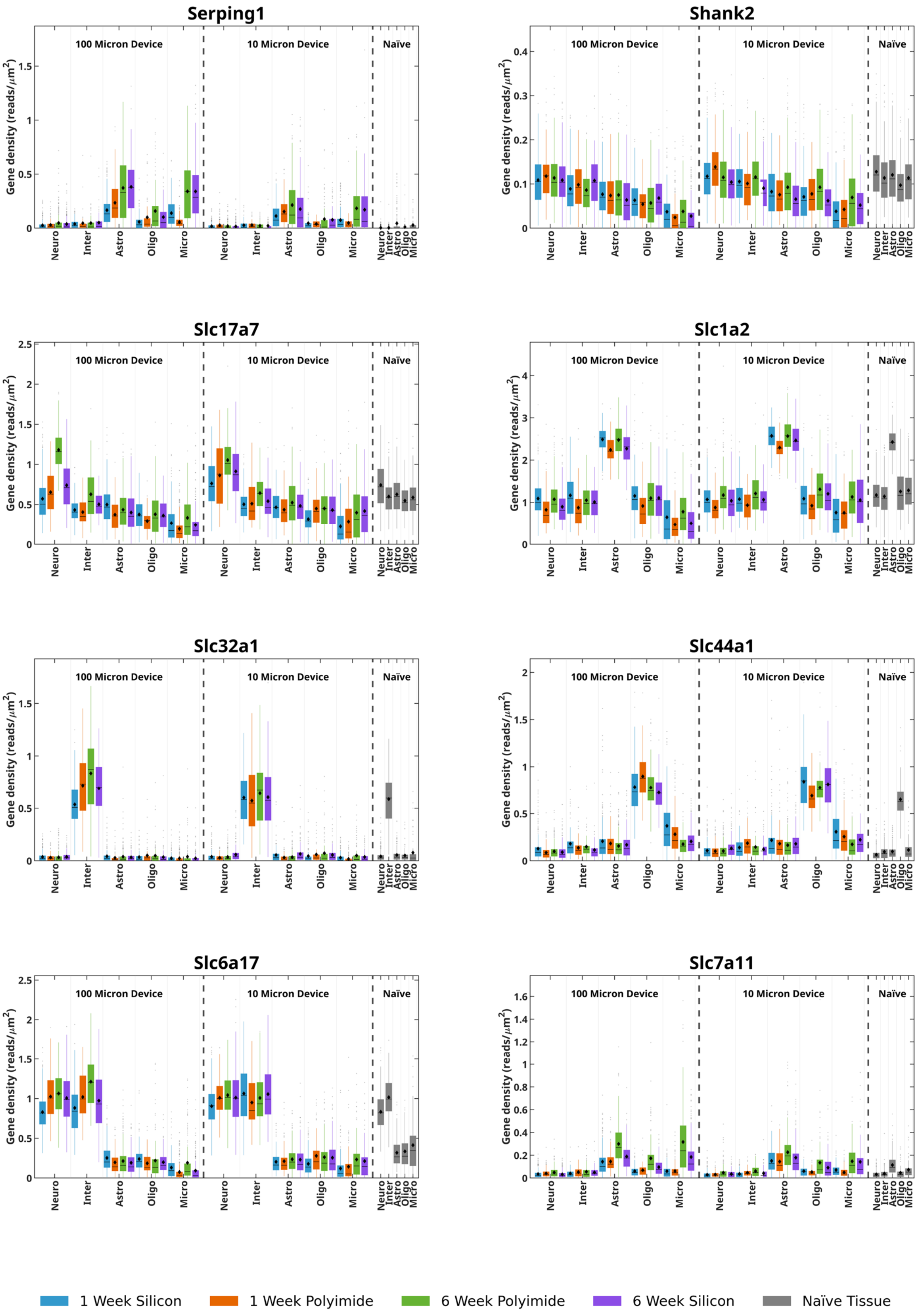

#### Supplementary Figure 31 (Contd.)

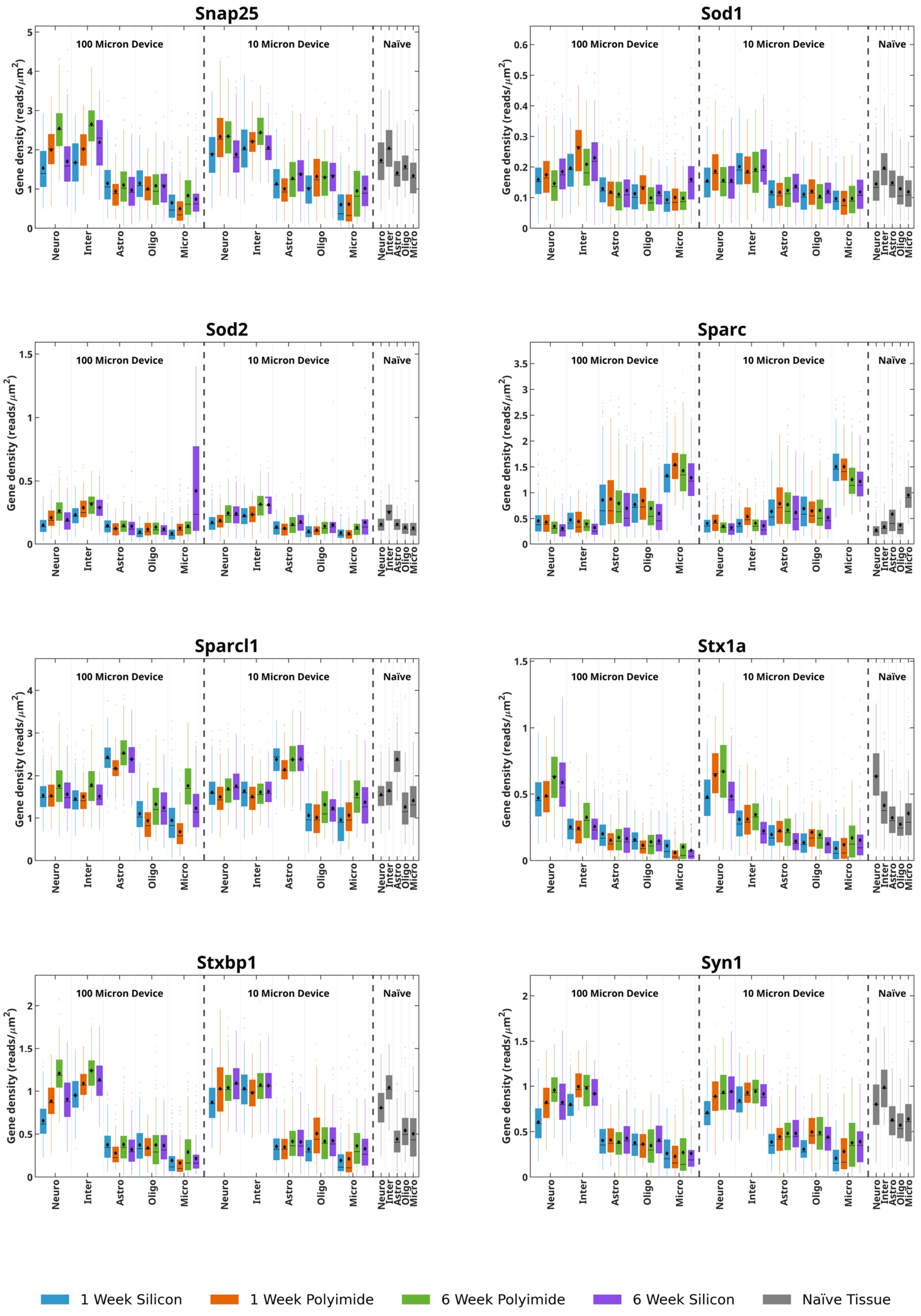

#### Supplementary Figure 31 (Contd.)

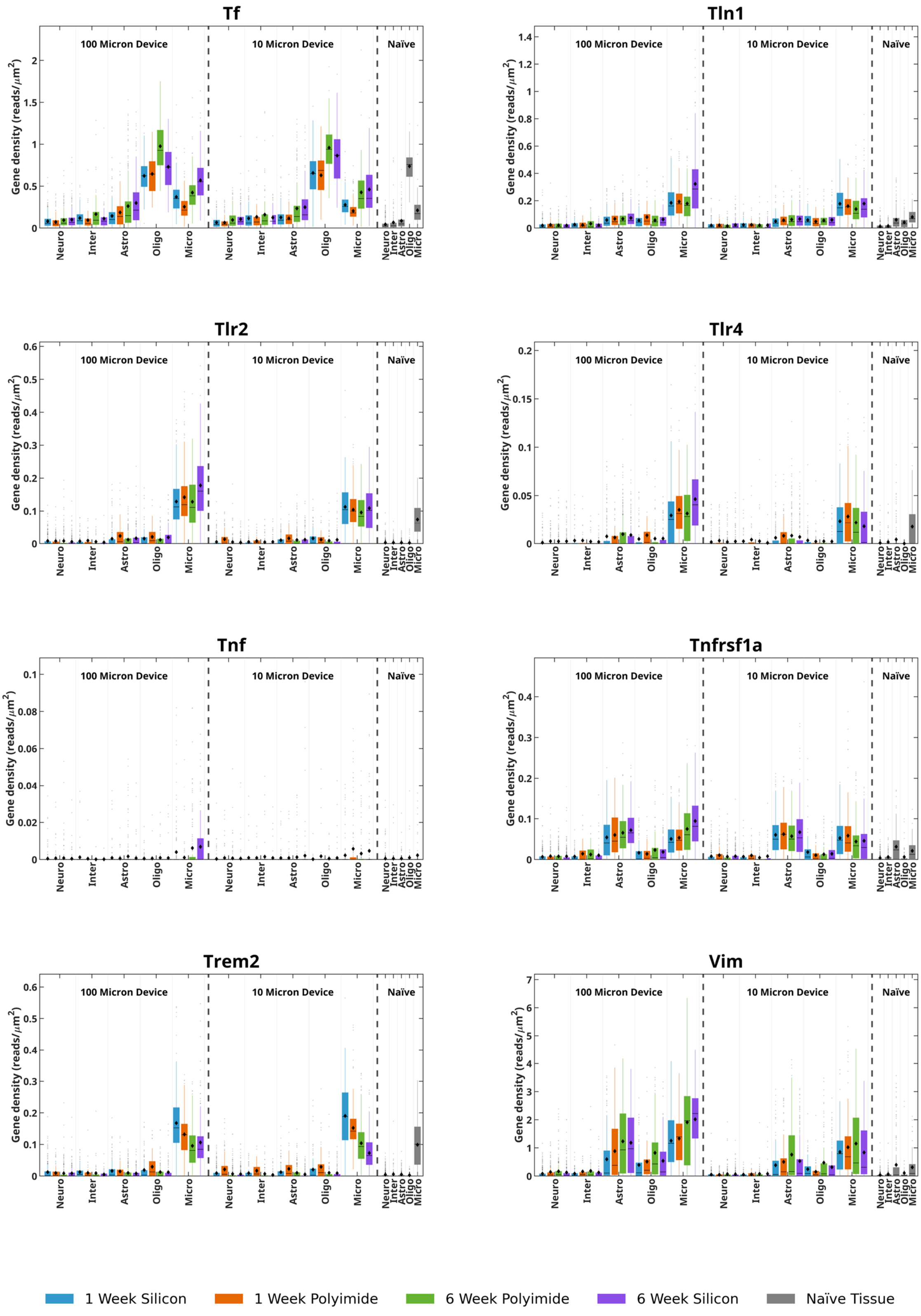

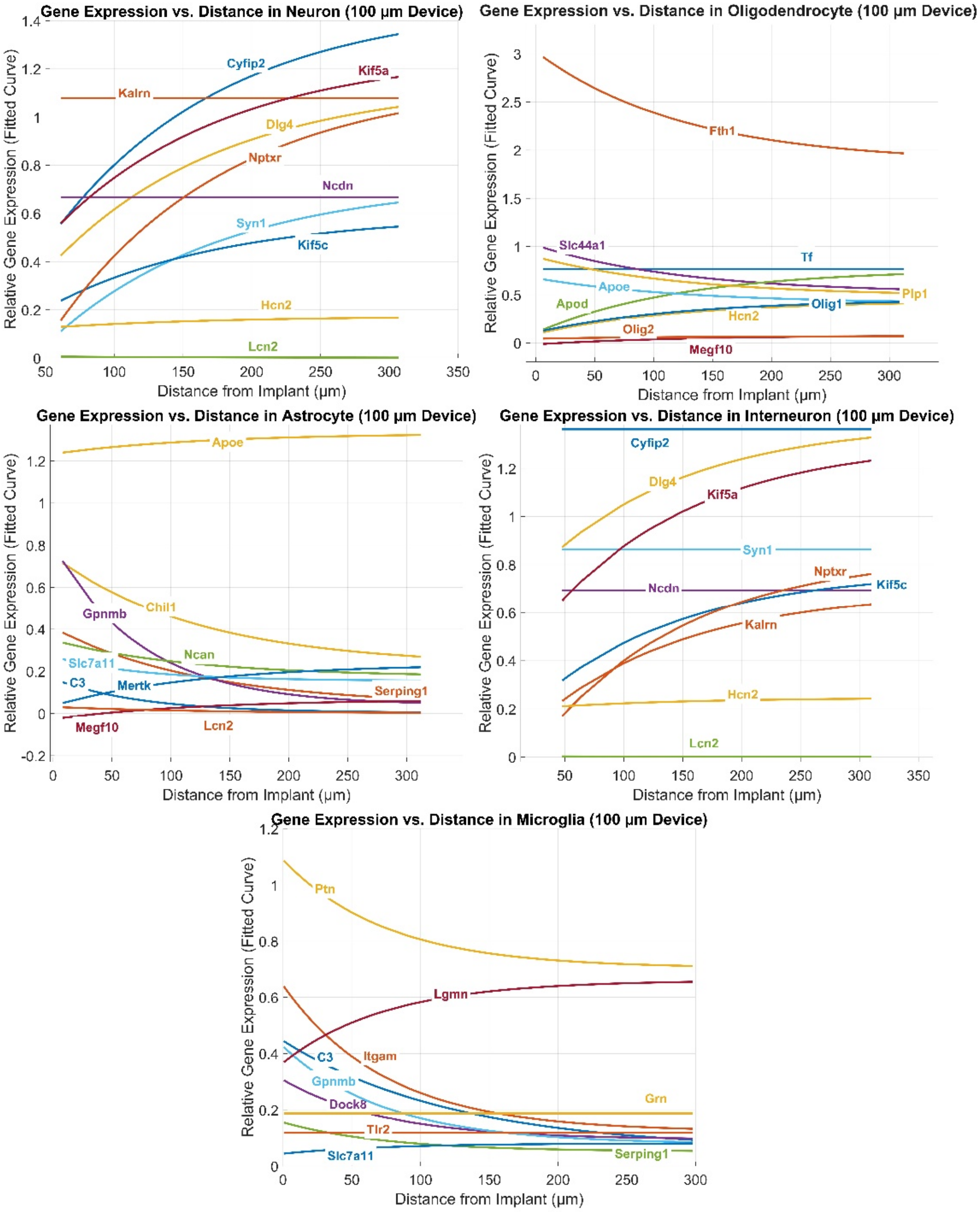


**Supplementary Figure 32:** Distance-dependent changes in relative gene expression for representative genes in neurons, interneurons, microglia, astrocytes, and oligodendrocytes surrounding 100 μm implants for Silicon 1-week ROI A (sample 1). Smoothed curves show fitted trends of normalized gene density as a function of distance from the device interface.

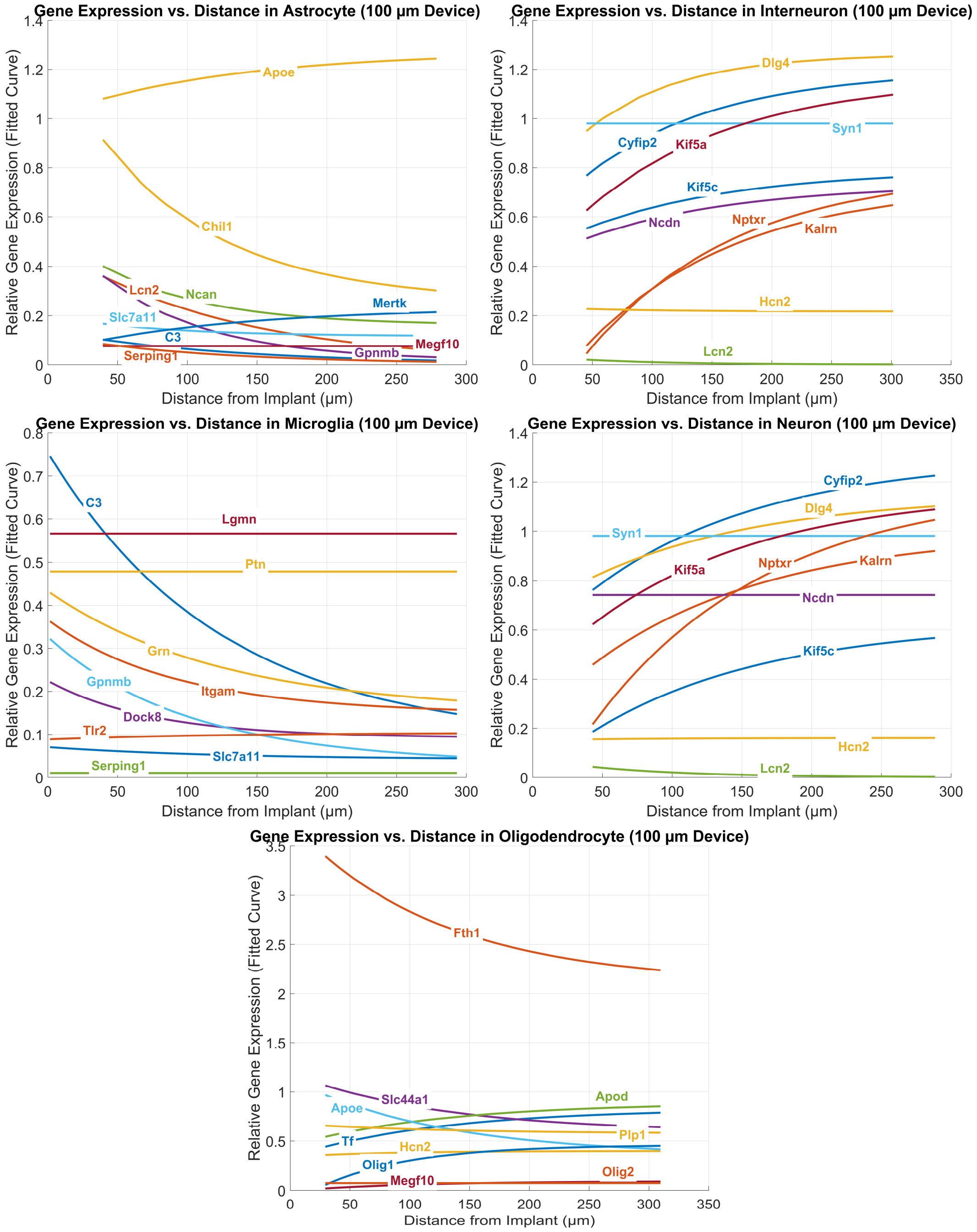


**Supplementary Figure 33:** Distance-dependent changes in relative gene expression for representative genes in neurons, interneurons, microglia, astrocytes, and oligodendrocytes surrounding 100 µm implants for Polyimide 1-week ROI B (sample 2). Smoothed curves show fitted trends of normalized gene density as a function of distance from the device interface.

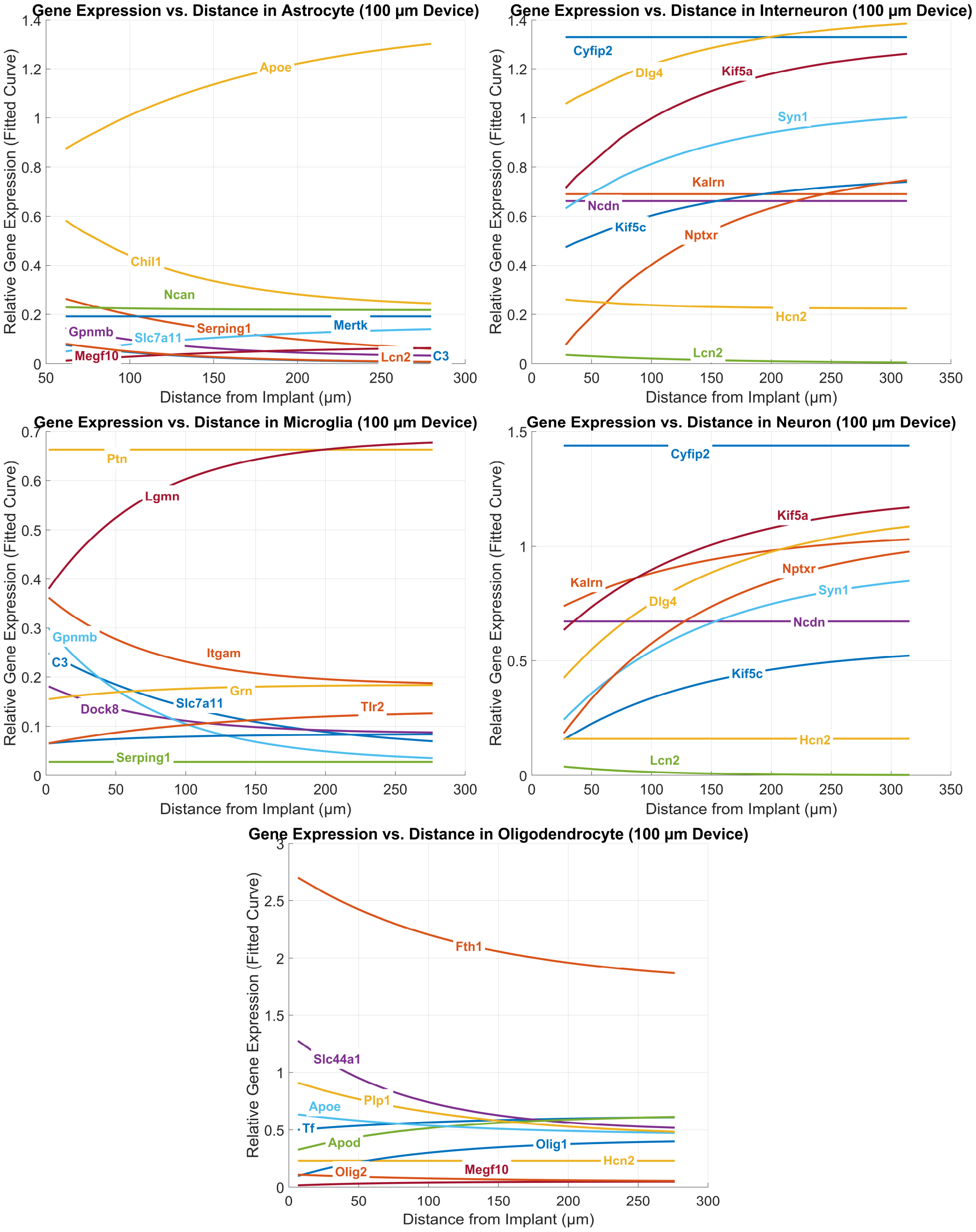


**Supplementary Figure 34:** Distance-dependent changes in relative gene expression for representative genes in neurons, interneurons, microglia, astrocytes, and oligodendrocytes surrounding 100 µm implants for Silicon 1-week ROI B (sample 3). Smoothed curves show fitted trends of normalized gene density as a function of distance from the device interface.

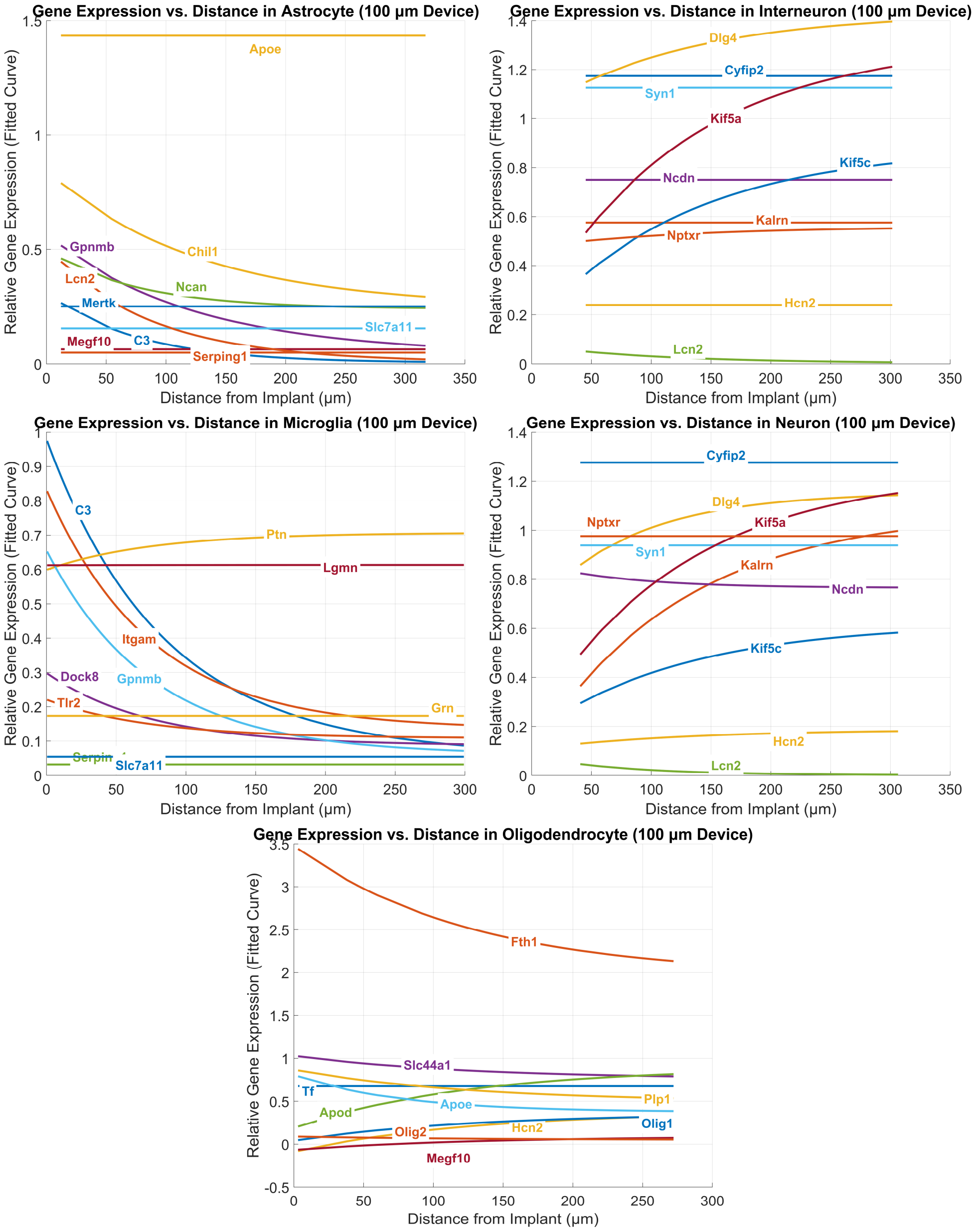


**Supplementary Figure 35:** Distance-dependent changes in relative gene expression for representative genes in neurons, interneurons, microglia, astrocytes, and oligodendrocytes surrounding 100 μm implants for Polyimide 1-week ROI C (sample 3). Smoothed curves show fitted trends of normalized gene density as a function of distance from the device interface.

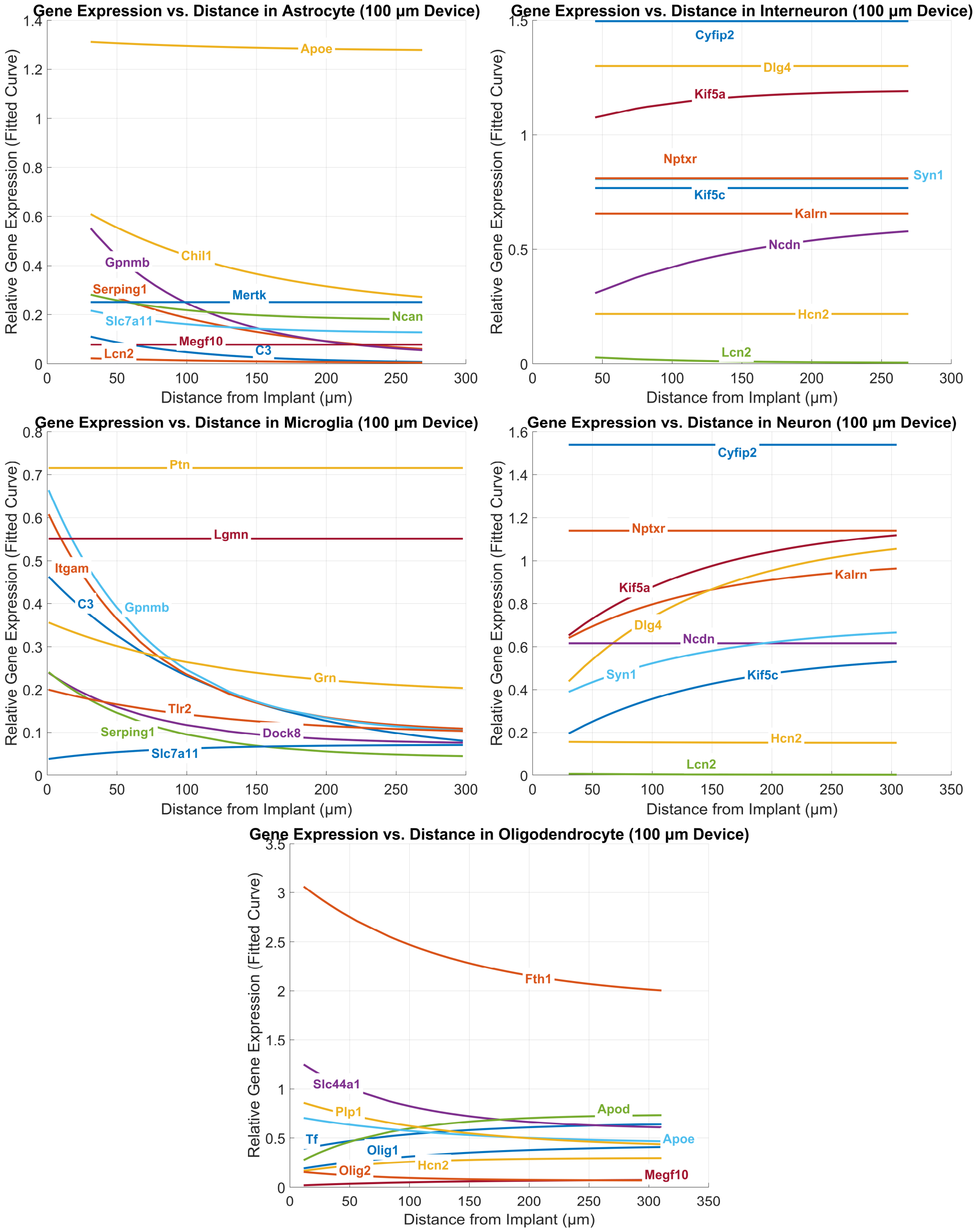


**Supplementary Figure 36:** Distance-dependent changes in relative gene expression for representative genes in neurons, interneurons, microglia, astrocytes, and oligodendrocytes surrounding 100 μm implants for Silicon 1-week ROI C (sample 3). Smoothed curves show fitted trends of normalized gene density as a function of distance from the device interface.

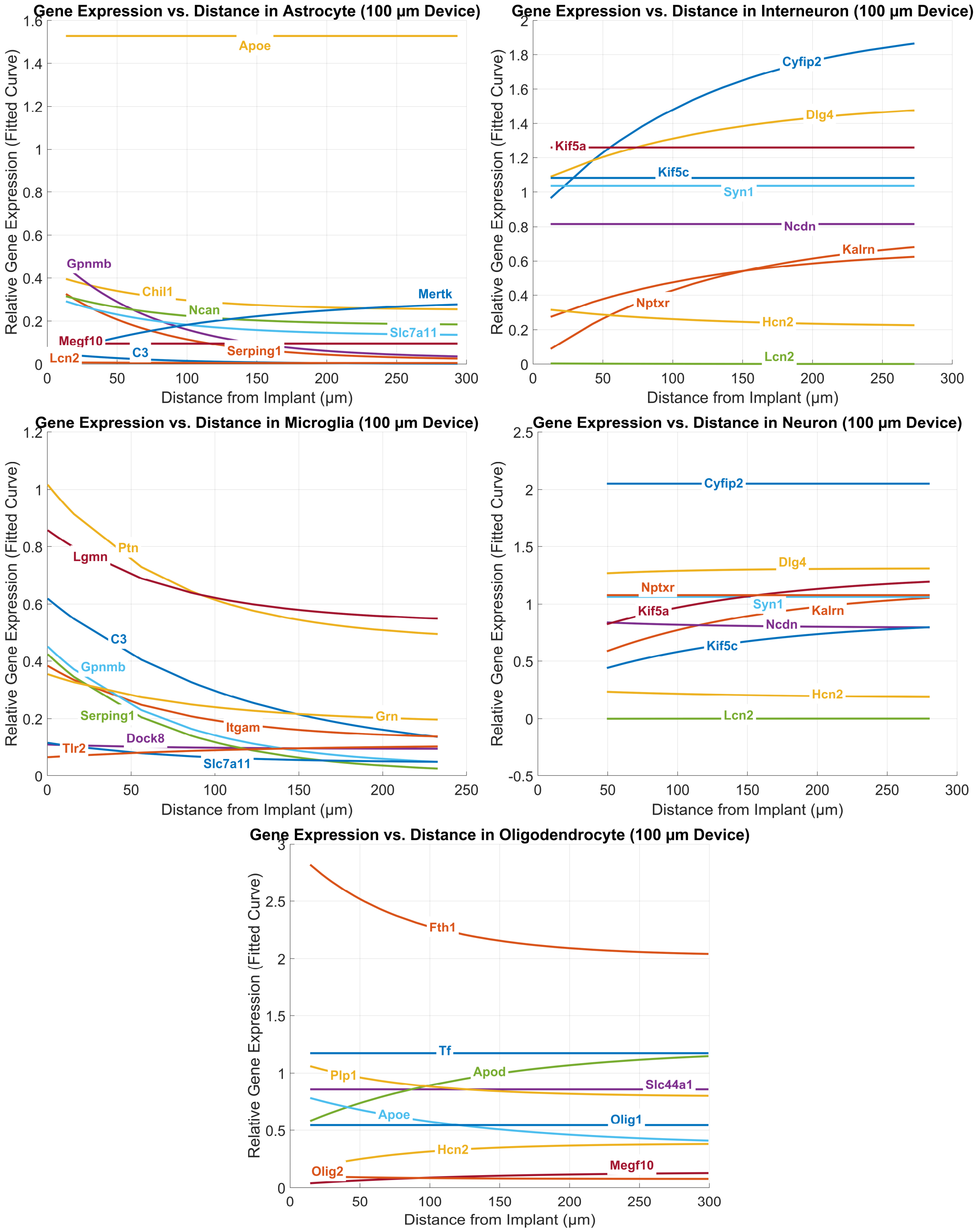


**Supplementary Figure 37:** Distance-dependent changes in relative gene expression for representative genes in neurons, interneurons, microglia, astrocytes, and oligodendrocytes surrounding 100 μm implants for Polyimide 6-week ROI A (sample 1). Smoothed curves show fitted trends of normalized gene density as a function of distance from the device interface.

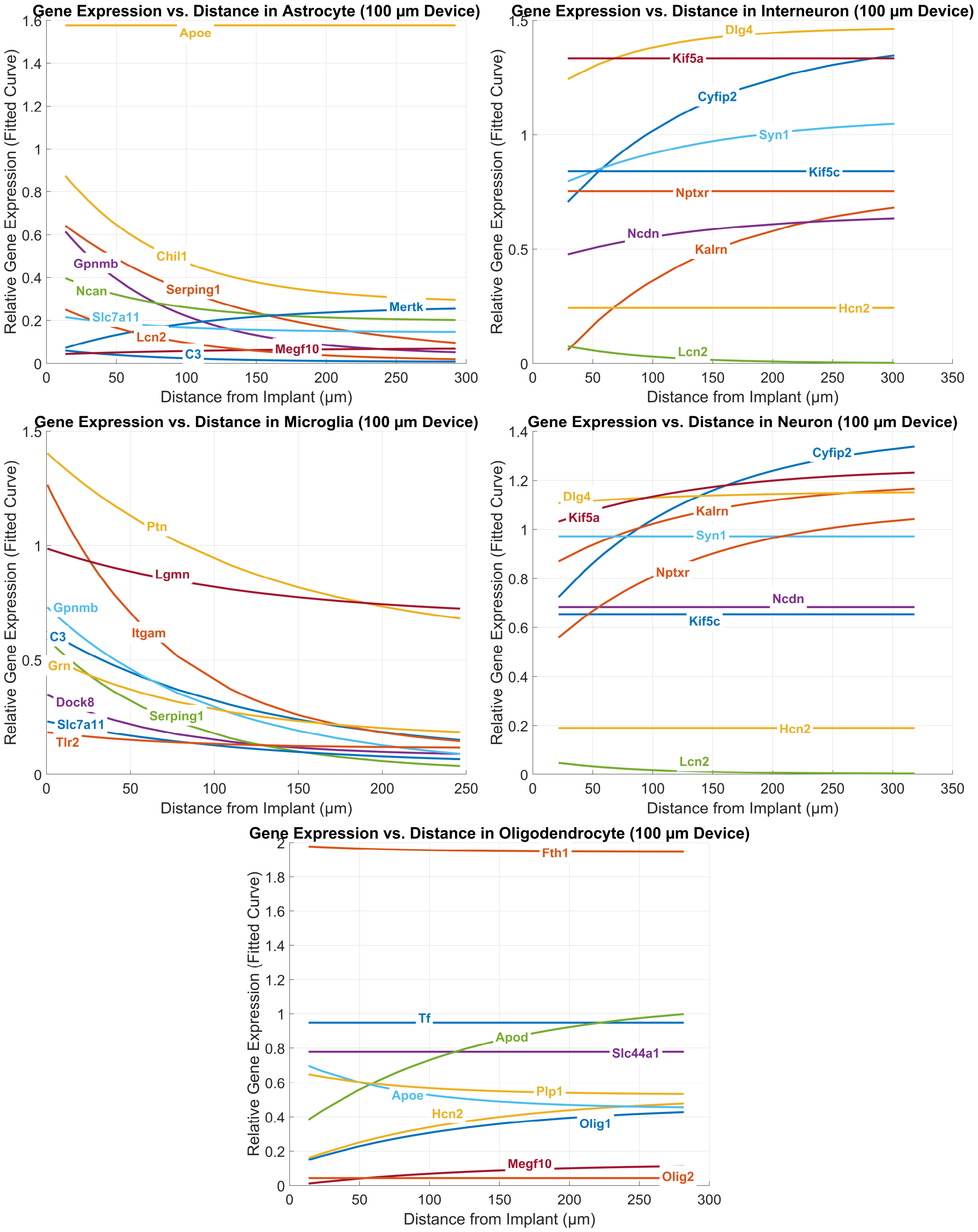


**Supplementary Figure 38:** Distance-dependent changes in relative gene expression for representative genes in neurons, interneurons, microglia, astrocytes, and oligodendrocytes surrounding 100 μm implants for Silicon 6-week ROI A (sample 1). Smoothed curves show fitted trends of normalized gene density as a function of distance from the device interface.

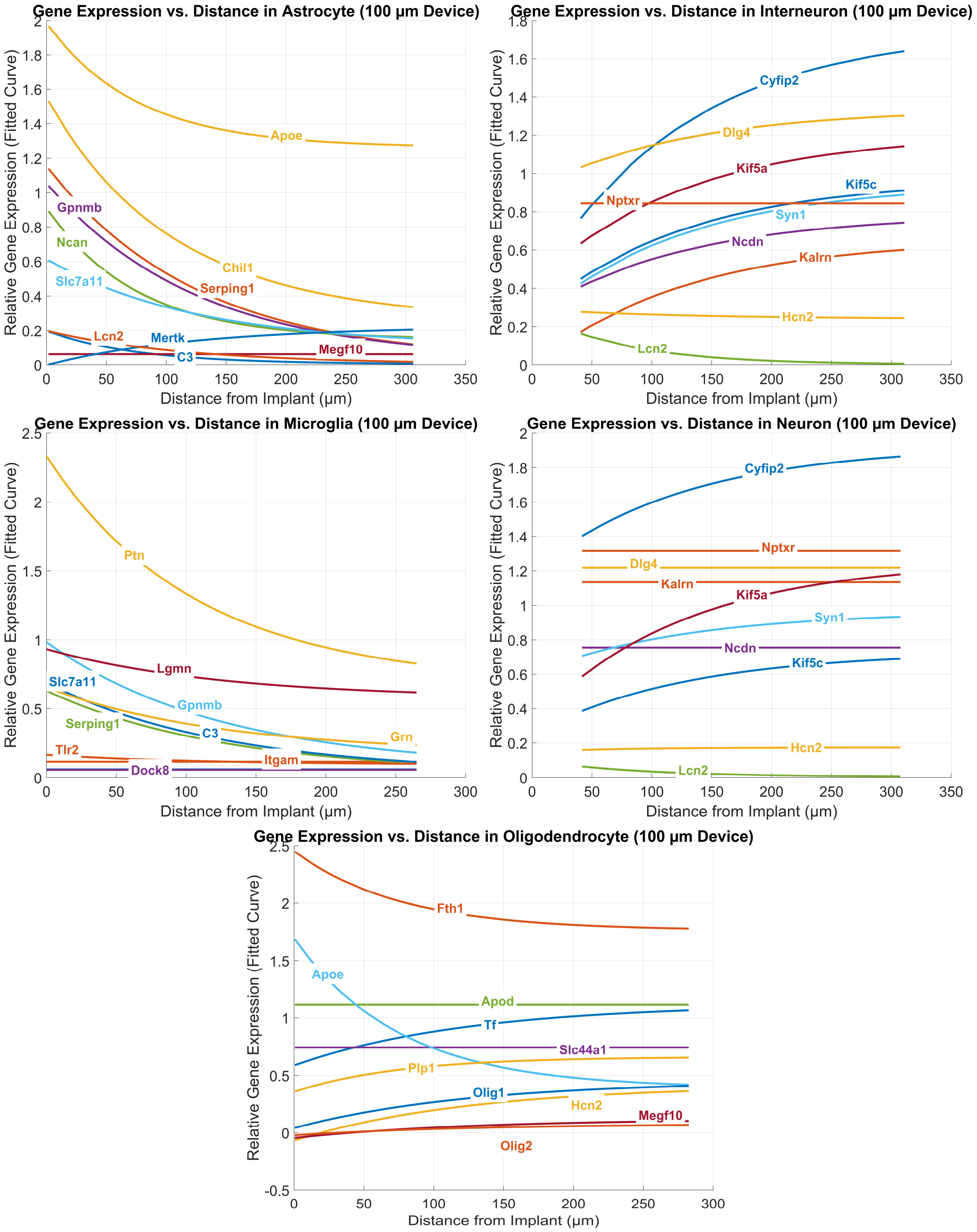


**Supplementary Figure 39:** Distance-dependent changes in relative gene expression for representative genes in neurons, interneurons, microglia, astrocytes, and oligodendrocytes surrounding 100 μm implants for Polyimide 6-week ROI B (sample 2). Smoothed curves show fitted trends of normalized gene density as a function of distance from the device interface.

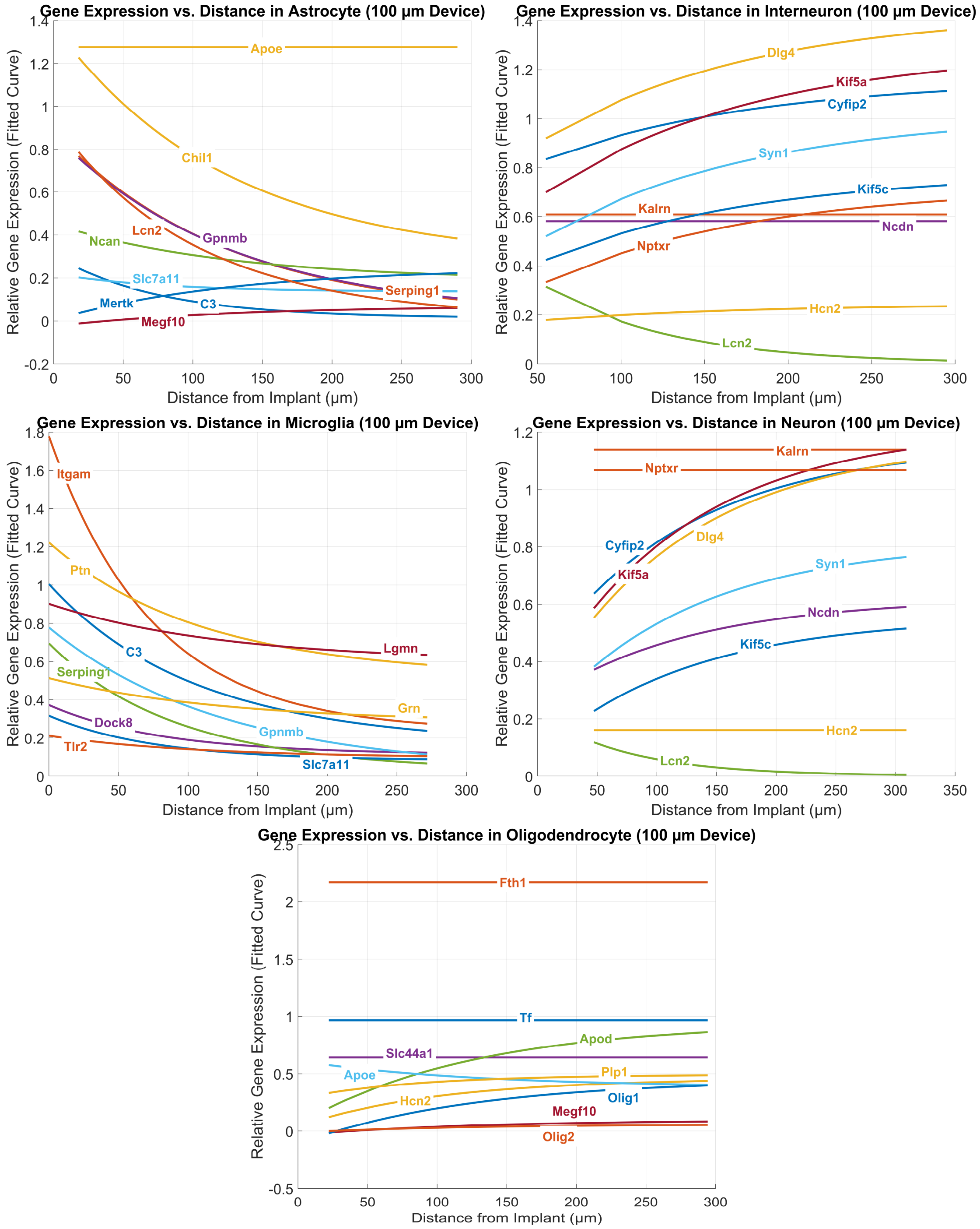


**Supplementary Figure 40:** Distance-dependent changes in relative gene expression for representative genes in neurons, interneurons, microglia, astrocytes, and oligodendrocytes surrounding 100 μm implants for Silicon 6-week ROI B (sample 2). Smoothed curves show fitted trends of normalized gene density as a function of distance from the device interface.

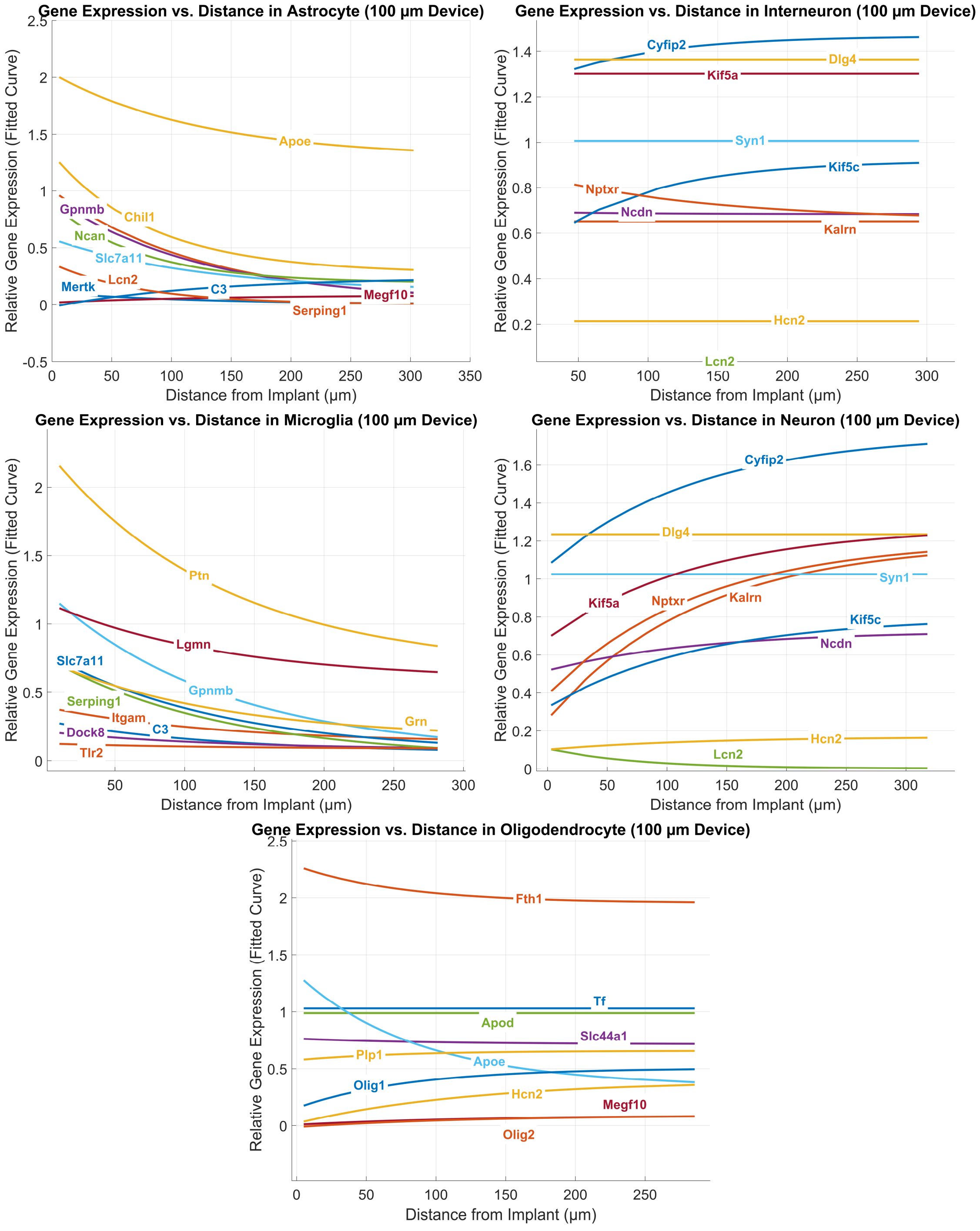


**Supplementary Figure 41:** Distance-dependent changes in relative gene expression for representative genes in neurons, interneurons, microglia, astrocytes, and oligodendrocytes surrounding 100 μm implants for Polyimide 6-week ROI C (sample 3). Smoothed curves show fitted trends of normalized gene density as a function of distance from the device interface.

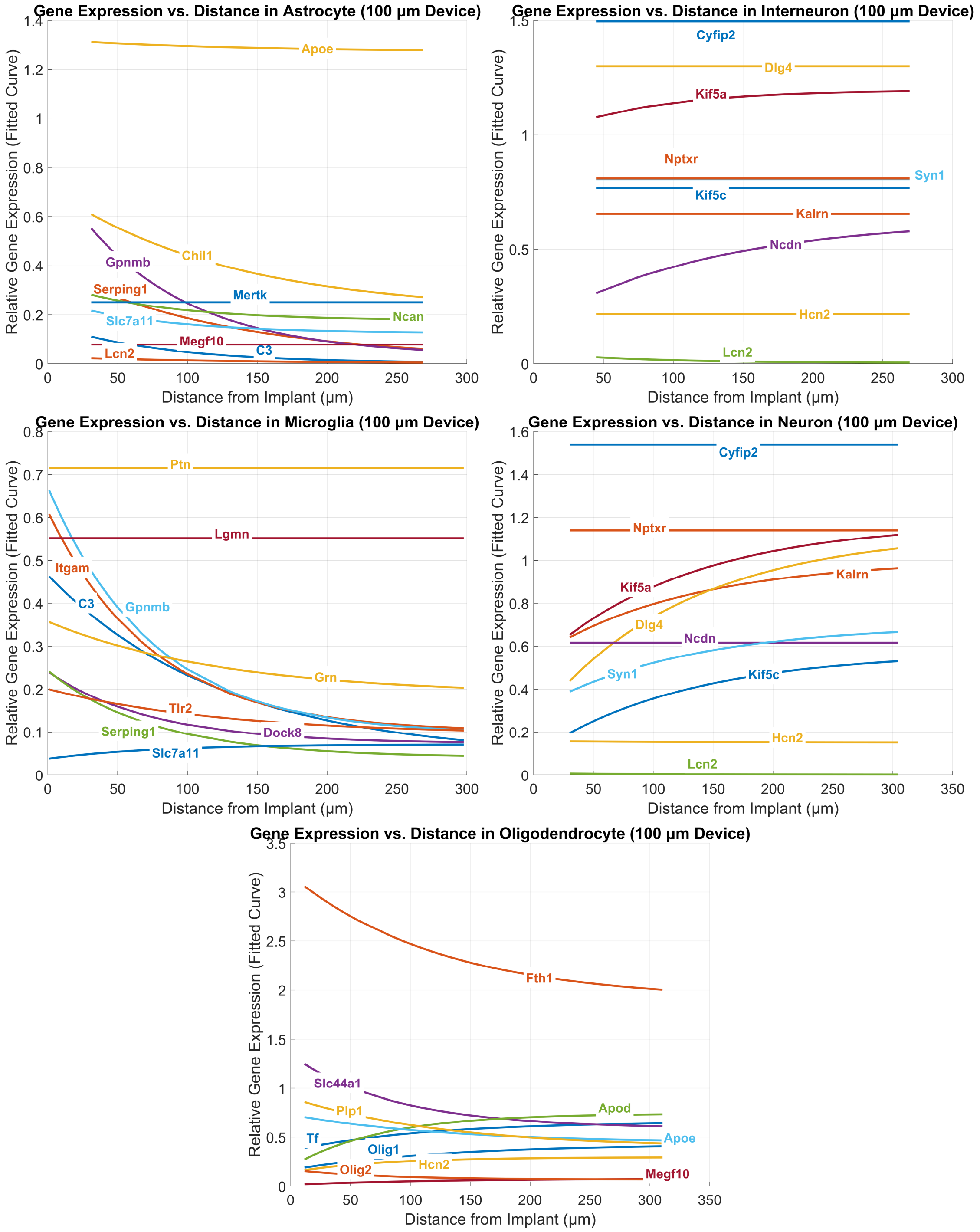


**Supplementary Figure 42:** Distance-dependent changes in relative gene expression for representative genes in neurons, interneurons, microglia, astrocytes, and oligodendrocytes surrounding 100 µm implants for Silicon 6-week ROI C (sample 3). Smoothed curves show fitted trends of normalized gene density as a function of distance from the device interface.

**Supplementary Figure 43:** Relative expression of genes expressed in oligodendrocytes within 60 µm of electrode implants compared against an equivalent area of tissue 150 µm from implanted electrodes. A paired t-test was used to evaluate the significance between 60 µm and 150 µm tissue.

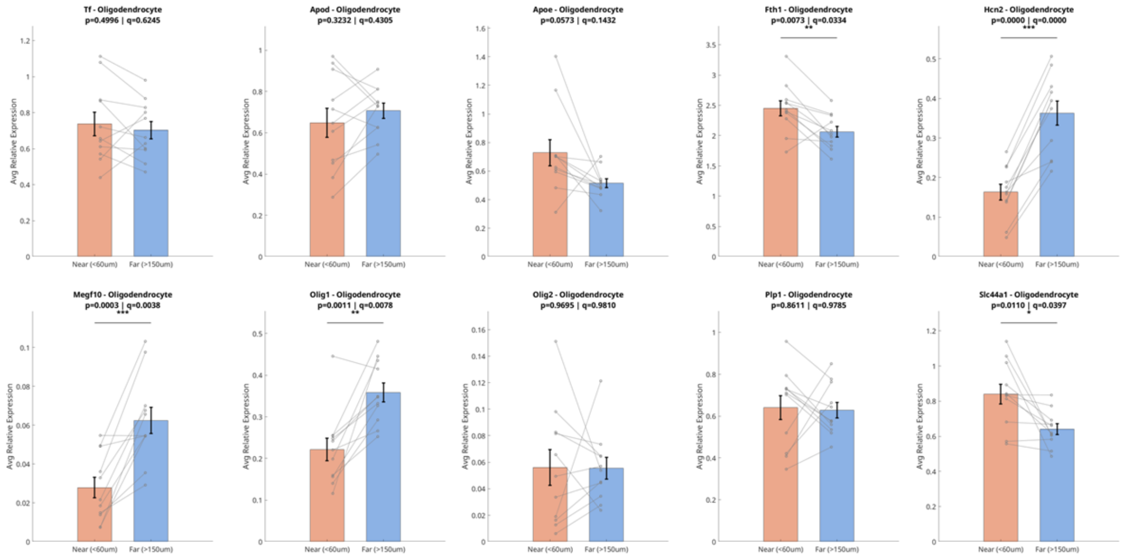


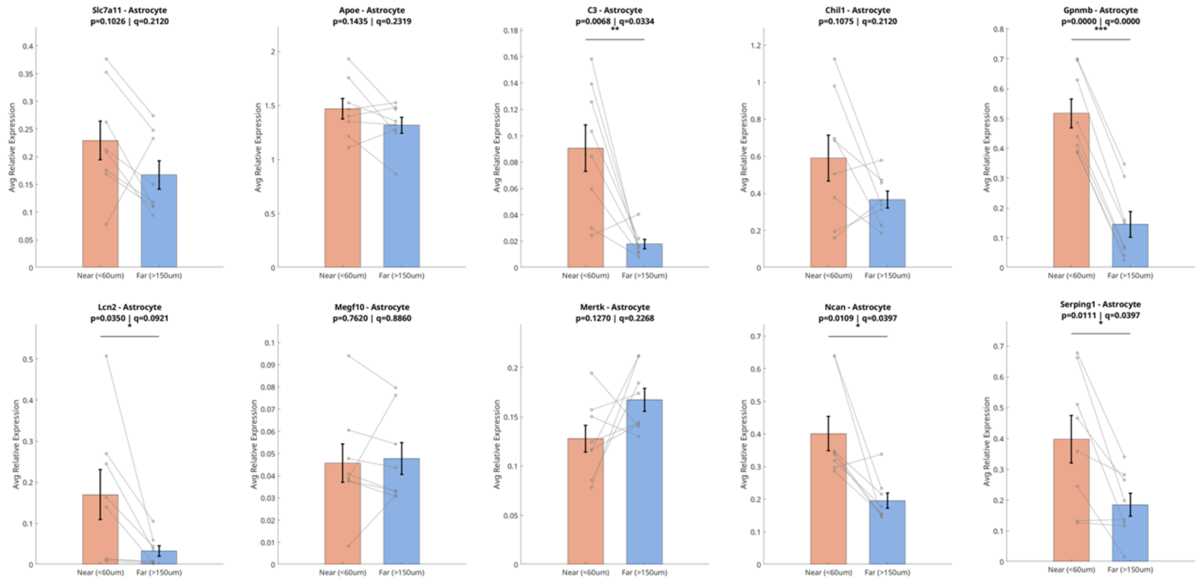

#### Supplementary Figure 43 (Contd.)

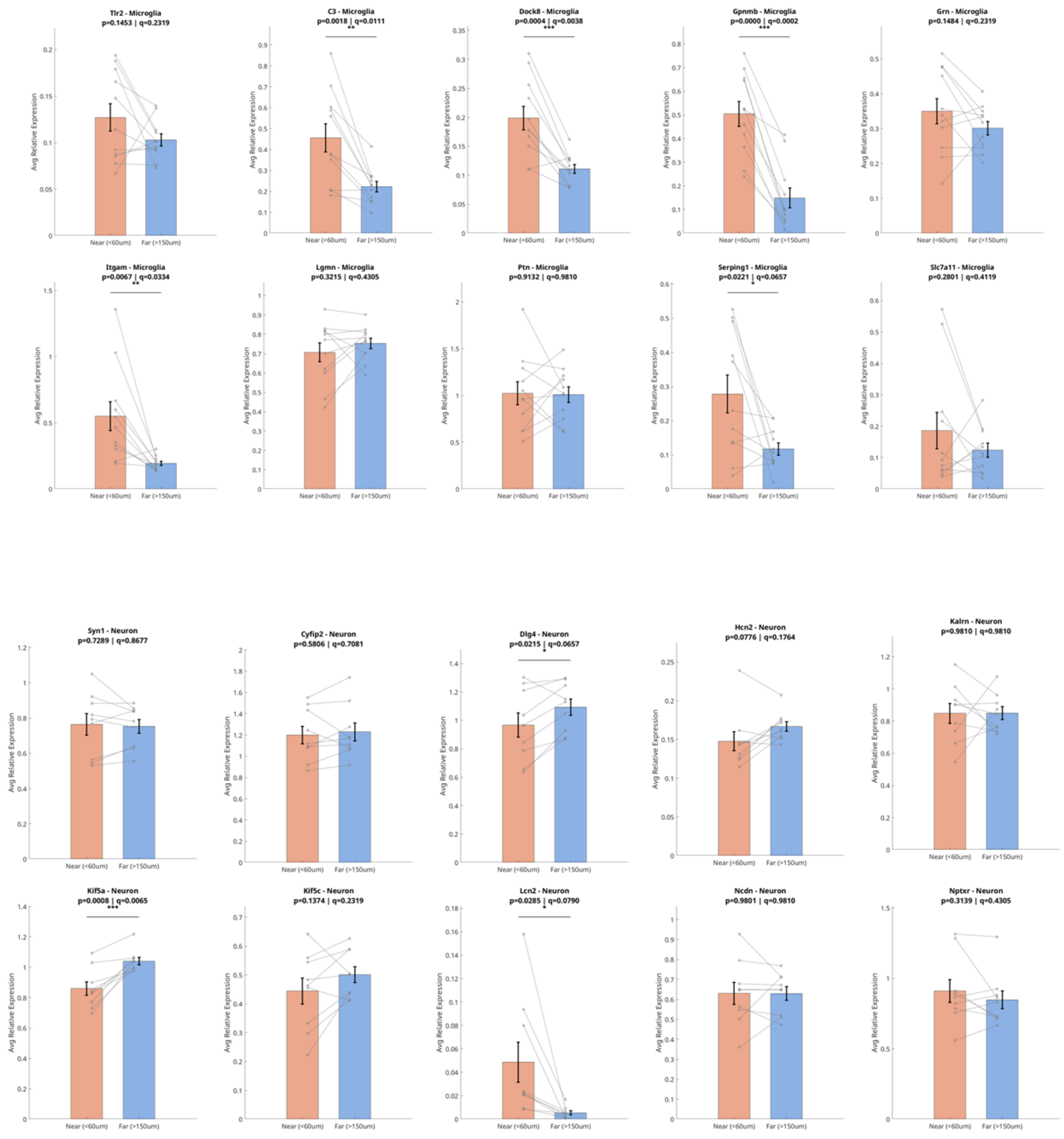

**Supplemental Discussion**

#### Gene expression UMAP analysis reveals distinct cell type-associated clusters at the device interface

The results of this study have generated UMAP-based cellular clusters, and quantitative differential expression of genes between clusters has identified possible phenotypes of individual cells in the observed cortical regions and at the implant interface based on the custom 100-gene panel used in this study. Based on the differential expression profiles of the UMAP-generated clusters, we have sorted the clusters into 4 distinct groups (Reactive Glia (RG1, RG2, RG3, RG4, and RG5); Modulatory Glia (MG1, MG2, MG3, MG4, and MG5); Oligodendrocytes (O1 and O2); and Neurons (NC1, NC2, NC3, and NC4)) by gene expression and the spatial distribution of these clusters around the implant injury (Supplementary Figures 35-39). Within individual xenium analyses, these clusters may have been assigned different numerical identifiers. However, the gene expression profiles of these clusters are highly conserved across samples, and we have renamed them under a uniform identifier for ease of understanding and greater simplicity in the results. The original cluster identifiers, as well as their individual gene expression profiles, can be referred to in the supplementary figures 21-25. Reactive astrocytes and microglia often progressively encapsulate implanted devices over time. In this study, we identified five clusters of reactive glia (RG1–RG5) that localize to the device interface, to perivascular regions and to the glia limitans (Supplementary Figure 26). These clusters appear stratified in a manner that mirrors the organization of the glia limitans. RG5 is tightly organized at the core of the device injury and shows high expression of innate immune and complement genes (*Cxcl2*, *C1qa*, C1qb, C1qc, *C1qc*, *C3*, *Itgam*, *Csf1r*). Immediately beyond RG5, we observed a mixture of two reactive clusters—RG4 (*Serping1*, *Gpnmb*, *GFAP*, *Vim*, *Lcn2*, *Aqp4*, *Ncan*) and RG3 (*Serping1*, *Vim*, *Apod*, *Tln1*, *Cxcl2*, *Dock8*)—that display distinct profiles of reactive astrocyte and microglial genes. RG1 lies peripheral to RG3–RG5, constitutes much of the glial scar and the glia limitans, and exhibits a comparatively less reactive transcriptional profile (*GFAP*, *Aqp4*, *Ncan*, *Slc7a11*, *Gpnmb*, *Chil1*). Interspersed within the scar and throughout the cortex tex is RG2, which is found exclusively around cortical vasculature and is characterized by the expression of *Vim*, *Serping1*, *Apod*, *Cspg4* and *Lcn2*. These clusters are not only present at the device interface but are also highly enriched in perivascular niches and at the glia limitans, which delineates the boundary between the meninges, vasculature and neural parenchyma. Beyond the reactive glial core at the device interface, we identified five clusters of modulatory glia (MG1–MG5) interspersed within the reactive glial scar and throughout the imaged brain regions (Supplementary Figure 27). Unlike RG1–RG5, the MG clusters are generally less localized to the implant injury and display mixed modulatory and reactive gene programs. MG1 cells primarily expressed *Megf10*, *Mertk*, *Tnfrsf1a*, *Ptprz1*, *Chil1*, *Aqp4* and *Ncan*. MG2 expressed *Lcn2*, *Nes*, *Dusp1*, *Tln1*, *Mbp* and *Cspg4*. MG3 showed high expression of inflammation-associated genes, including *Tlr2*, *Cx3cr1*, *Itgam*, *Trem2*, *Csf1r*, *C1qa*, *C1qb*, *C1qc* and *C3*. MG4 shared features with MG2 (e.g., *Lcn2*, *Nes*, *Dusp1*, *Tln1*) but was distinguished by expression of *Sparc*, *Vim*, *Tnf*, *Shank2*, *Cnksr2* and *Grn*. MG5 overlapped transcriptionally with MG3 (*Tlr2*, *Cx3cr1*, *Itgam*, *Trem2*, *Csf1r*, *C1qa*, *C1qb*, *C1qc*, *C3*) but uniquely included *Kalrn*, *Brinp1*, *Stx1a*, *Grin1*, *Nptxr* and *Npc2*, suggesting proximity to neurons. Based on spatial correspondence between defined clusters and immunofluorescence, RG2 and RG4 did not align with $NeuN^+$ or $Olig2^+$ nuclei; instead, these clusters coincided with GFAP$^+$ nuclei and other Hoechst$^+$ cells. MG3 cells localized broadly to Hoechst$^+$ nuclei across the section and in proximity to NeuN$^+$ cells, whereas MG5 was almost exclusively associated with NeuN$^+$ cells and Hoechst$^+$ nuclei adjacent to neuronal somata. This study also identified two clusters of cells that are present across all samples and express genes associated with the oligodendrocyte lineage (Clusters **O1** and **O2**) (Supplementary Figure 28). Cluster O1 expresses genes associated with mature and myelinating oligodendrocytes (*Olig1*, *Olig2*, *Mog*, *Mbp*, *Plp1*, *Slc44a1*, *Hcn2*, *Apod*), in addition to high relative expression of iron metabolism genes *Tf*, *Fth1*, and *Ftl1*. Cluster O2 does not express *Mog*, *Mag*, *Mbp*, or *Plp1*, and instead expresses genes associated with both OPCs and mature oligodendrocytes (*Cspg4*, *Olig1*, *Olig2*, *Ptprz1*, *Slc44a1*). Both O1 and O2 appear evenly distributed throughout the cortex and the implantation injury, with no apparent changes in density near or far from the device interface. The results of this study indicated four consistently identified neuronal clusters that distinctly express genes associated with excitatory and inhibitory neurons and are highly associated with NeuN$^+$ nuclei (**NC1**, **NC2**, **NC3**, **NC4**) (Supplementary Figure 29). Clusters NC1 and NC2 highly express genes associated with excitatory neurons (*Slc17a7*), as well as pan-neuronal genes related to axonal transport (*Kif5a*, *Kif5b*, *Kif5c*, *Dynll1*), synaptic transmission (*Dlg4*, *Bsn*, *Snap25*, *Syn1*, *Stxbp1*, *Ncdn*, *Stx1a*, *Nrgn*, *Slc6a17*, *Camk2a*, *Nptxr*), cytoskeletal structure (*Nefh*, *Nefm*, *Nefl*, *Dst*, *Map4*), and dendritic spine maintenance (*Kalrn*, *Cyfip2*, *Shank2*). Clusters NC3 and NC4 express genes associated with inhibitory interneurons (*Slc31a1*, *Gad1*), as well as pan-neuronal genes related to axonal transport (*Kif5a*, *Kif5b*, *Kif5c*, *Dynll1*), synaptic transmission (*Dlg4*, *Bsn*, *Snap25*, *Syn1*, *Stxbp1*, *Ncdn*, *Stx1a*, *Nrgn*, *Slc6a17*, *Grin1*,

*Gria1*, *Camk2a*), cytoskeletal structure (*Nefh*, *Nefm*, *Nefl*, *Dst*, *Map4*), and dendritic spine maintenance (*Kalrn*, *Cyfip2*, *Shank2*). We also observed *Mmp9* expression within some excitatory neurons, as well as *Sod2* within inhibitory neurons throughout the tissue section and at the device interface.
